# SMYD5 is a regulator of the mild hypothermia response

**DOI:** 10.1101/2023.05.11.540170

**Authors:** Salvor Rafnsdottir, Kijin Jang, Sara Tholl Halldorsdottir, Meghna Vinod, Arnhildur Tomasdottir, Katrin Möller, Katrin Halldorsdottir, Tinna Reynisdottir, Laufey Halla Atladottir, Kristin Elisabet Allison, Kevin Ostacolo, Jin He, Li Zhang, Frances J. Northington, Erna Magnusdottir, Raul Chavez-Valdez, Kimberley Jade Anderson, Hans Tomas Bjornsson

**Author notes:** Program in Genetic Counseling, Stanford University (Arnhildur Tomasdottir).

## Abstract

The mild hypothermia response (MHR) maintains organismal homeostasis during cold exposure and is thought to be critical for the neuroprotection documented with therapeutic hypothermia. To date, little is known about the transcriptional regulation of the MHR. We utilize a forward CRISPR-Cas9 mutagenesis screen to identify the histone lysine methyltransferase SMYD5 as a regulator of the MHR. SMYD5 represses the key MHR gene *SP1* at euthermia. This repression correlates with temperature-dependent levels of H3K36me3 at the *SP1*-locus and globally, indicating that the mammalian MHR is regulated at the level of histone modifications. We have identified 37 additional SMYD5 regulated temperature-dependent genes, suggesting a broader MHR-related role for SMYD5. Our study provides an example of how histone modifications integrate environmental cues into the genetic circuitry of mammalian cells and provides insights that may yield therapeutic avenues for neuroprotection after catastrophic events.

## Main Text

In clinical practice, physicians actively lower core temperature with the use of Targeted Temperature Management (TTM) also called therapeutic hypothermia. TTM is a therapeutic strategy to minimize neurological damage following neonatal hypoxic-ischemic brain injury and cardiac arrest^1–8^. The lack of increased benefit for temperatures below 33.5°C as well as the observed upregulation of several genes at mild hypothermic temperatures suggests that this pathway mediates the neuronal benefits of TTM ^9–11^. In support of this, researchers have found several genes (e.g., *CIRBP/CIRP*, *RBM3*) which consistently show increased expression during mild hypothermia^12^. Two of these, *SP1* and *RBM3,* appear to have roles in decreasing neuronal death in disease states^13–17^. Therefore, it is important for researchers and clinicians to understand both the mechanisms and the extent of this response.

SP1, a transcription factor, binds to the promoter region of *CIRBP*^18^ upon cold exposure at a sequence called the Mild Cold Response Element (MCRE). Its binding leads to increased expression of *CIRBP* at 32°C indicating regulation at the transcriptional level. Similarly, RBM3 is known to regulate at least *one* downstream gene, *RTN3*, a neuroprotective factor in a neurodegeneration model^16^. This suggests that mammalian cells have a pathway culminating in SP1/CIRBP and RBM3/RTN3. Little has been done to elucidate the transcriptional regulation of this pathway and relatively little knowledge is available regarding the mammalian MHR compared to heat shock responses, which have been extensively characterized^19^.

Histone modifications have previously been found to help integrate the effects of cold temperature exposure in diverse organisms, but there are no known mammalian mechanisms. In plants, vernalization is the basis of cold exposure influence on the rate of flowering and is mediated through a Trithorax-Polycomb switch^20^. Some reptiles such as the red eared slider turtle (*Trachemys Scripta Elegans*), use temperature-dependent sex determination and this regulation acts through a histone methylation switch^21^. A detailed understanding of the mammalian MHR is important to begin charting 1) the impact of climate change on mammals and 2) how temperature can systematically bias results in biology and medicine. An obvious example of the latter involves the process of harvesting tissue from human cadavers: these are uniformly kept at a low temperature, which may activate the mild hypothermia response in the cells being harvested. Similarly, countless researchers store samples at room temperature between experimental steps, which in some scenarios may activate the MHR. Knowing which genes may be influenced by temperature would constitute crucial information for many investigators studying genome-wide expression levels or protein levels.

Here, we use unbiased strategies to gain clues into the MHR upstream of *SP1*. Specifically, we use the *SP1* <u>m</u>ild <u>h</u>ypothermia indicator (MHI) containing the *SP1* promoter (SP1-MHI) to perform genome-scale CRISPR-Cas9 forward mutagenesis screen^22–24^, reading out *SP1*-promoter activity using the SP1-MHI. Our study yielded multiple candidate regulators of the MHR. We focused on one of these regulators, *SMYD5*, a histone methyltransferase. We demonstrate that SMYD5 binds to and directly represses SP1 by various independent experiments. We further show that total protein levels of SMYD5 are decreased at 32°C compared to 37°C through proteasomal degradation, and that a subset of other SMYD5-bound and repressed genes at 37°C show significant differential expression upon mild hypothermia. Our insights elucidate how an external cue (mild hypothermia) specifically mediates its effects on the mammalian genome through the actions of histone modifications. This work paves the way towards the development of therapeutic means to treat neonatal hypoxic-ischemic brain injury, a major cause of mortality and morbidity in otherwise healthy newborns^25^, without TTM.

## Results

### Mild hypothermia indicators show increased fluorescence at 32°C

Core temperature is kept at a very tight range in humans (euthermia, 36.5-37.8°C)^26^. By definition, mild hypothermia involves temperature ranges from 32-36°C (**Fig. S1**, bold). To directly visualize the promoter activity of the three genes previously shown^9,12,18^ to respond to mild hypothermia (*CIRBP*, *RBM3*, *SP1*), we have designed eight distinct MHIs (**table S1** and **file S1**). Each of our MHI has a promoter material with or without upstream MCRE sequences driving the expression of either a Green Fluorescent Protein (GFP) or mCherry (**Fig. 1A**). When transfected into HEK293 cells, the MHIs demonstrate increased fluorescence at 32°C compared to 37°C (**Fig. 1B–1C**). These findings are mirrored by increased endogenous SP1 levels in a Western blot from our cell system collected after 16 hours (h) at 32°C (**Fig. 1D**). We assessed the temporal dynamics of promoter activity using flow cytometry analysis of a representative indicator and observed a maximal increase in fluorescence of SP1-MHI between 6-8 h after a temperature shift to 32°C (**Fig. 1E**). This finding was supported by continuous monitoring of fluorescence of three MHIs where maximal intensity was reached within 6-8 h, with SP1-MHI, rising fastest, followed by RBM3-MHI, and then CIRBP-MHI (**Fig. S2A-C**). MHIs were also transiently transfected into HEK293 cells and then exposed to five distinct temperatures (26°C, 29°C, 32°C, 37°C, and 40°C), for 16 h followed by flow cytometry (**Fig. 1F-1H; Fig. S2D-E**). Consistent with previous literature^9,12,18^ and our own validation in HEK293 cells (**Fig. 1B-D**), indicators representing all three genes (*SP1*, *CIRBP*, *RBM3)* consistently had increased fluorescence at 32°C compared to 37°C/40°C (P<0.01, **Fig. 1F-1H**). SP1- and CIRBP-MHIs do not show increased activity at lower (26°C, 29°C, **Fig. S2D-E**) or higher (40°C) temperatures (**Fig. 1F-1G**) and the response was strongest at 32°C (**Fig. 1F-1H**). SP1-MHI is not activated after exposing HEK293 cells to the pro-oxidant agent H_2_O_2_ (**Fig. S2F**), suggesting that neither metabolic derangement nor apoptosis is able to activate the response. These data support the notion that our MHIs specifically reflect the MHR and are not activated by general cell stress (i.e., moderate hypothermia, hyperthermia, or apoptosis). Furthermore, we observe comparable results in six other distinct lines (**Fig. S3**). These data suggest that all three genes are regulated at the transcriptional level emphasizing the importance of understanding the transcriptional regulation of this response. The time-lag in the response upon hypothermia is measured in hours rather than minutes, indicating that there must be additional regulators operating upstream of these genes. Since many of the key studies in the field have used HEK293^16,18,27^ and these cells have neuronal characteristics^28^, we used this cell line for our screening approaches. However, when needed, we cross-validated our findings using either human or murine neural progenitor cells or *in vivo* tissues.

**Fig. 1.**
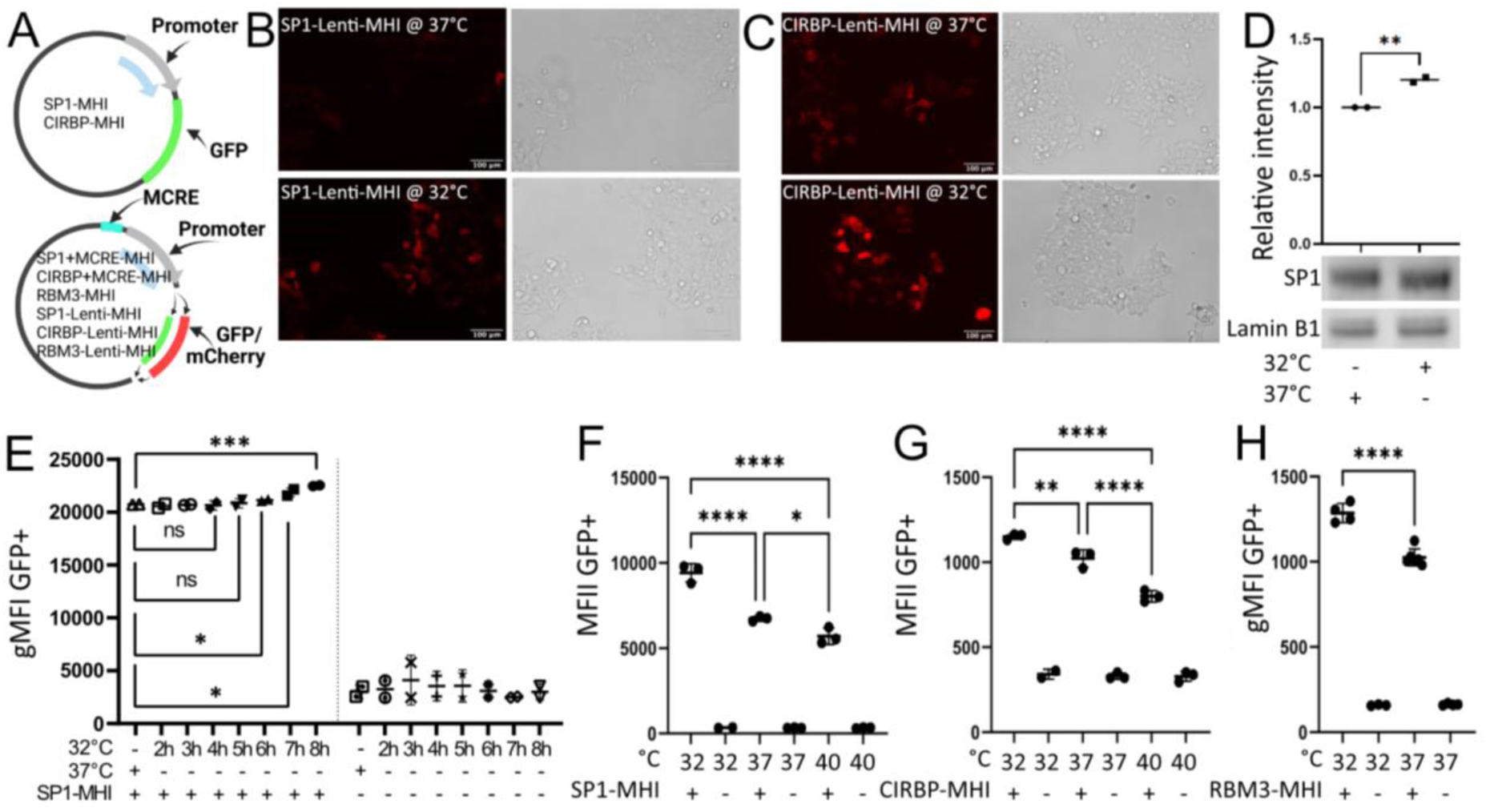
MHIs allow single cell fluorescent quantification of MHR activation. (**A**) A schematic of MHI structures. (**B-C**) Representative fluorescent images of SP1-Lenti-MHIs and CIRBP-Lenti-MHI, respectively, at 37°C and 32°C. (**D**) A Western blot demonstrating increased SP1 after 16 h at 32°C compared to 37°C. Each data point is a biological replicate (n=2), and their mean is depicted. Significance levels were calculated with an unpaired two-tailed t-test. (**E**) geometric Mean Fluorescence Intensity (gMFI, flow cytometry) after varying lengths of hypothermia exposure for SP1-MHI. Each data point is a technical replicate (n=2), with mean and standard deviation (SD) where applicable. Significance levels were calculated with an unpaired one-tailed t-test. (**F-G**) Mean fluorescence (MFI, flow cytometry) of SP1-MHI and CIRBP-MHI, respectively (16 h at 32°C, 37°C, 40°C). Each point is a technical replicate (n=3), with mean and SD where applicable. Significance levels were calculated with Šidák‘s multiple comparison test. (**H**) gMFI of RBM3-MHI (16 h at 32°C and 37°C). Each point is a technical replicate (n=4-5), with mean and SD where applicable. Significance levels were calculated with an unpaired one-tailed t-test. All significance levels in this figure were calculated in GraphPad Prism, * = P<0.05, ** = P<0.01, *** = P<0.001, **** = P<0.0001.

### CRISPR-Cas9 Knockout Screen with SP1 -MHIs reveals candidate regulators

To map regulators of the cooling response (*SP1*) in an unbiased manner, we used an available genome-coverage lentiGuide-Puro pooled sgRNA library^22–24^ in combination with our SP1-MHI to uncover transcriptional regulators of *SP1*. We chose to use the SP1-MHI, as our and others prior data indicate that this gene regulates *CIRBP* at the transcriptional level^18^ and it is the MHR gene that is the first to achieve maximal response at 32°C (**Fig. S2A-C**). We performed the screen on HEK293WT+Cas9 cells expressing SPI-MHI (HEK293WT+Cas9+SP1), transducing the sgRNA library at ∼0.3 MOI sgRNA per cell. After puromycin selection and exposure to hypothermia (32°C) for 16 h (to activate the MHR), we sorted out the 5% most and least fluorescent cells (**Fig. 2A**) to capture repressors and activators of *SP1*, respectively. Some would argue that storing cells on ice (i.e., during fluorescent activating cell sorting (FACS)) could possibly lower the temperature of cells to the hypothermic range and activate the MHR. We found that prolonged storage on ice (8 h) at room temperature was unable to activate the MHI (**Fig. S4)**. The screen yielded a consistent shift towards lower fluorescence for the transduced HEK293WT+Cas9+SP1-MHI (green) compared to the positive control (non-transduced HEK293WT+Cas9+SP1-MHI, grey, **Fig. 2B**). The observation of such a large fluorescent shift of cells towards less SP1-MHI fluorescence at hypothermia may suggest the involvement of many sgRNAs. This indicates a large number of putative genes that are upstream of *SP1*, potentially more activators than repressors. We amplified guides for each sorted group and a negative control (transduced HEK293WT+Cas9) and performed next-generation sequencing. To identify guides enriched in the sorted populations, we used the MaGeCK pipeline^29^ to computationally predict genes with enrichment (**Fig. 2C-D**). We looked at all genes that had a - Log10(Robust Rank Aggregation (RRA)) score above 3.5 and an FDR less than 0.25 to avoid missing possible regulators (**table S2**). We were left with a list of 74 genes for the SP1-repressors and 60 for SP1-activators. Of note, there were two microRNAs on the top of the SP1 repressor (*hsa-mir-4314*) and activator (*hsa-mir-6739*) lists. Interestingly, both are predicted to target SP1 (mirdb.org).

**Fig. 2.**
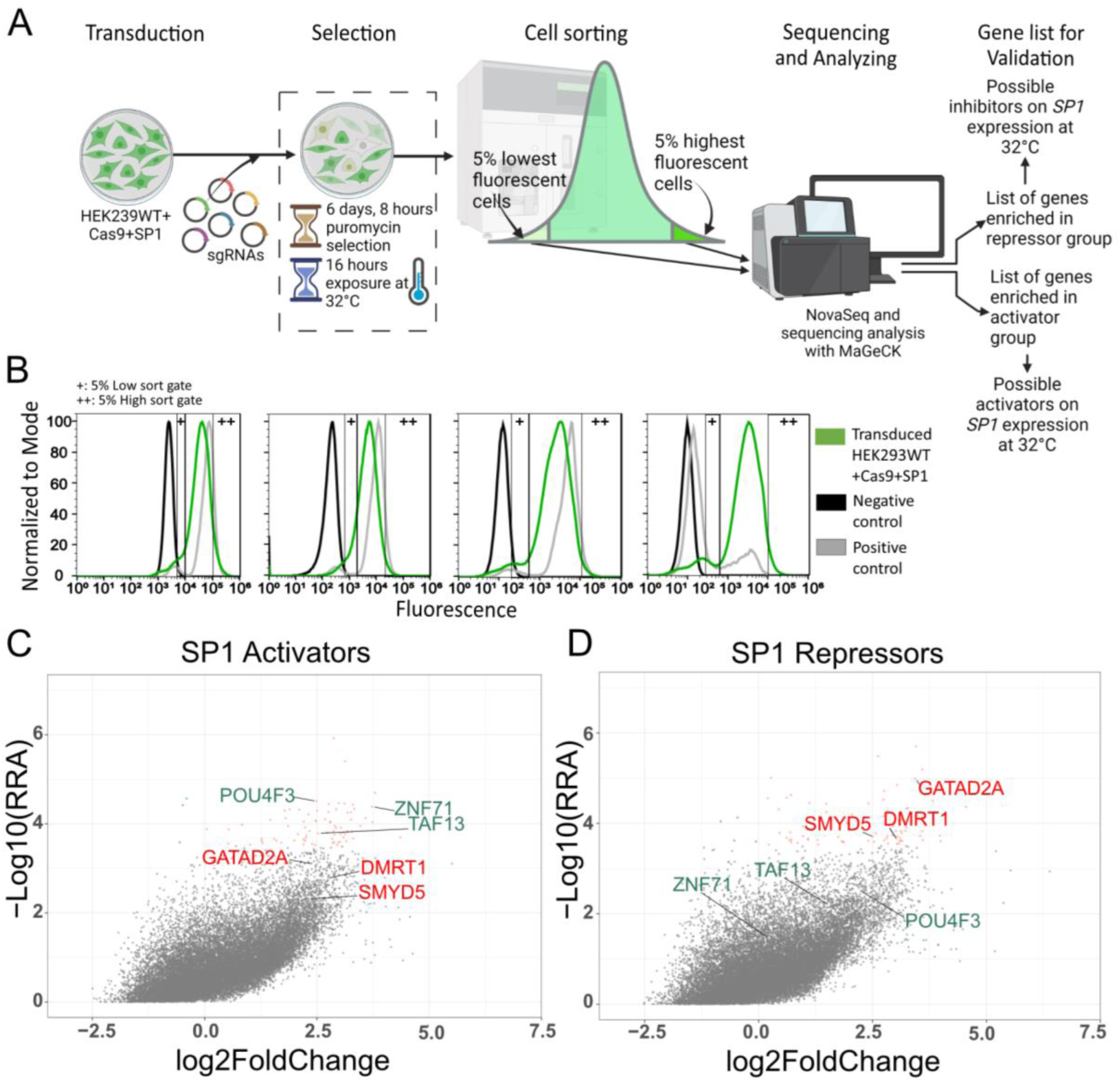
Genome-scale CRISPR-Cas9 knock out screen on SP1-MHI reveals multiple potential inhibitors and activators of the MHR. (**A**) Overview of the genome-scale CRISPR-Cas9 knock out approach for the HEK293WT+Cas9+SP1 cell line. (**B**) Fluorescence measurements and sort gates of the 4 replicates of transduced HEK293WT+Cas9+SP1 cells (green), negative control (HEK293WT, black) and positive control (HEK293WT+Cas9+SP1, grey). (**C-D**) Genes marked in red are transcription regulators that have a -Log10(RRA) score >3.5 and a known repressive function. Genes marked in green are transcription regulators that have a known activating function and a -Log10(RRA) score >3.5. Colored dots represent genes that have a -Log10(RRA) score >3.5, where the orange dots indicate genes that have a positive log fold change (LFC) and the blue dots indicate genes that have a negative LFC, from either the SP1 activator (C) or repressor (D) screen. -Log10(RRA) score and LFC was calculated with MAGeCK^29^.

We were motivated to find a possible transcriptional regulator of the MHR. We decided to look further at known transcriptional regulators which could either repress or activate in a way which was concordant with the shift of fluorescence (repressors with more, activators with less fluorescence). There were six genes (*GATAD2*, *POU4F3*, *ZNF71*, *SMYD5*, *DMRT1*, *TAF13*) from our lists that fulfilled these criteria, including *SMYD5*, encoding a histone methyltransferase. SMYD5 is known to catalyze the placement of H4K20, H3K9, and H3K36 trimethylation modifications^30,31^. Histone modification has previously been known to dictate temperature-dependent sex determination in reptiles and fish through KDM6B^21,32^, as well as vernalization in plants^20^. Interestingly, SMYD5 is a likely direct interactor with KDM6B in fish^33^. SMYD5 is also a known resident of stress granules like CIRBP and RBM3^34^.

### SMYD5 is a direct repressor of SP1

To further explore the role of *SMYD5*, we took advantage of a recently published dataset^30^ from a CUT&TAG assay in mouse Embryonic Stem Cells (mESC) with an overexpressed SMYD5-Flag plasmid. Upon re-analysis of these data, we noticed that SMYD5 has strong peaks at the promoters of both *Sp1* (130K) and *Cirbp* (70K), but not *Rbm3* (**Fig. 3A**). These peaks occur at CpG islands (**Fig. 3A**). We reanalyzed an RNASeq dataset from mESC with a *Smyd5* knock out (KO) from the same study ^30^. We observed approximately equal amounts of upregulated and downregulated differentially expressed genes (DEGs) following *Smyd5* KO among SMYD5 bound loci (**Fig. 3B**). This supports prior reports that histone marks deposited by SMYD5 can either be associated with active or repressive chromatin^30,31^ and H3K36me3 itself has been observed on both repressed and actively transcribed genetic sites^35^. When looking at the three loci (*Sp1*, *Cirbp*, *Rbm3*), we found that *Smyd5*-KO yielded increased expression of *Sp1* but not *Cirbp* or *Rbm3* (**Fig. 3C**). The direct binding and the increased expression suggest that SMYD5 acts as a direct repressor in a murine system. In the same dataset, *Cirbp* showed low expression compared to *Sp1* and *Rbm3* (**Fig. S5**), which may impede interpretation. To validate these findings in a human system, we knocked down (KD) *SMYD5* with siRNA in our HEK293WT+Cas9+SP1 cell line and observed an increased fluorescence of the SP1-MHI at 32°C and 37°C (**Fig. 3D**). *SMYD5*-KO with a guide RNA (gRNA) and Cas9 significantly decreased amounts of *SMYD5* mRNA expression as evaluated by qRT-PCR (**Fig. 3E**) and Western blot (**Fig. 3F**) using a previously validated antibody against SMYD5^36^. Both of which were rescued upon overexpression of a gRNA-resistant Flag-SMYD5 plasmid (Flag-SMYD5 sgRNAres plasmid [labelled SMYD5sgRNA#6res], **Fig. 3E and 3F**). The same experiment revealed de-repression of the *SP1* mRNA in KO cells by RT-qPCR, with repression restored in rescued cells (**Fig. 3G**). Furthermore, we observed a significant increase of SP1 and CIRBP but not RBM3 following *SMYD5*-KO at 37°C using a Western blot of the endogenous loci (**Fig. 3H-J**). Together these data suggest that SMYD5 is a direct regulator of *SP1/Sp1* in human and murine systems.

**Fig. 3.**
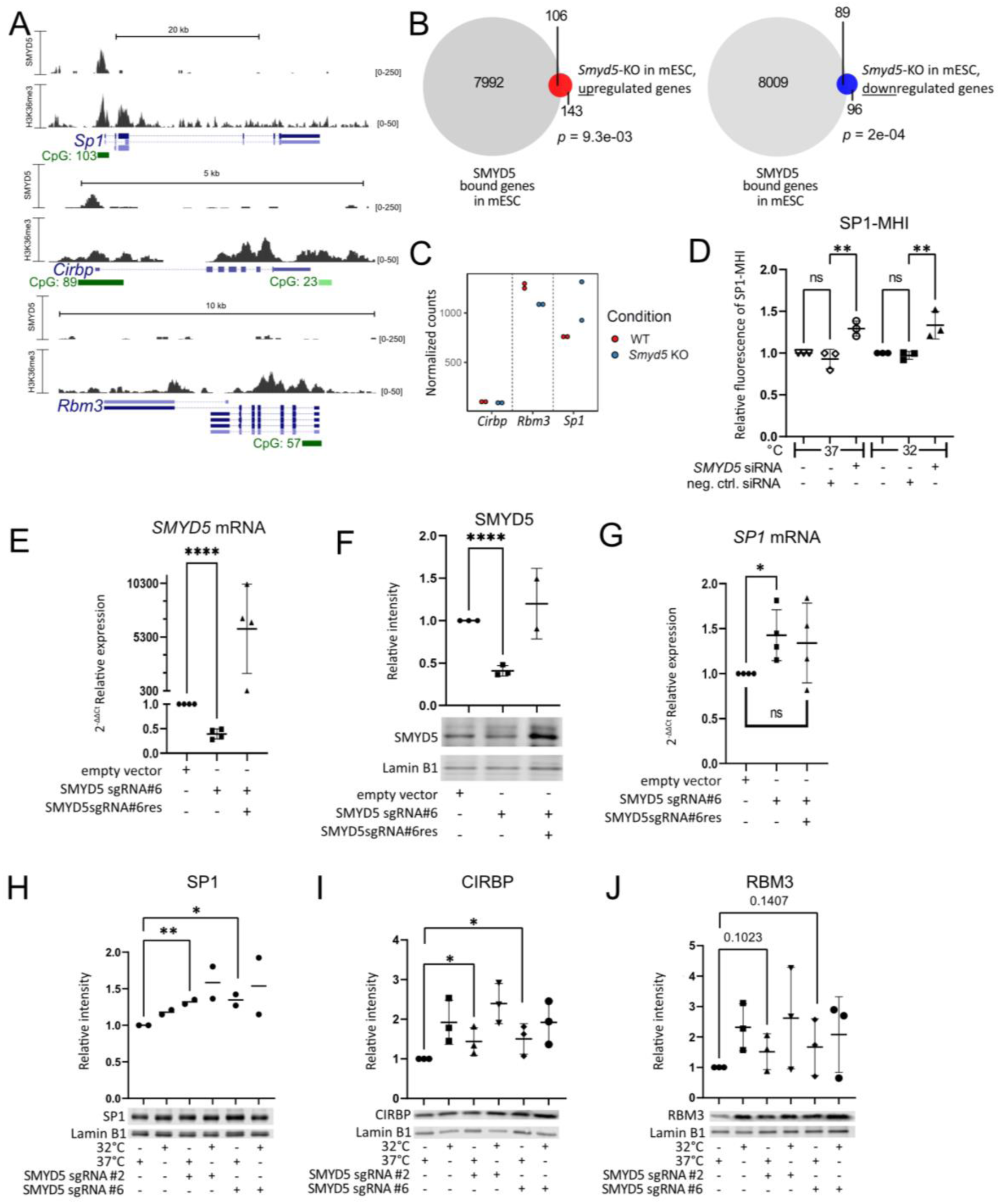
SMYD5 is a direct repressor of SP1 at 37°C. (**A**) Overexpressed FLAG-tagged SMYD5 (n=2) in mESc binds at promoters of *Sp1* and *Cirbp* but not of *Rbm3*. H3K36me3 peaks over *Sp1*, *Cirbp* and *Rbm3* promoter and gene body regions. Data from Zhang et. al.^30^ (**B**) Venn graph showing both up- and downregulated SMYD5-bound genes in an RNASeq of *Smyd5*-KO cells (n=2). Statistical analysis was done with GeneOverlap^57^ package in R using Fisher’s Exact test. (**C**) *Smyd5*-KO leads to increased mRNA expression of *SP1* but not *RBM3* or *CIRBP* in mESCs. (**D**) *SMYD5*-KD by siRNA yields higher levels of fluorescence of SP1-MHI at 32°C and 37°C compared to empty vector control. Each data point is a biological replicate (n=3), that has been normalized against the same non-transfected HEK293WT+Cas9+SP1-MHI cell line, mean and SD are depicted where applicable. Significance levels were calculated with Šidák‘s multiple comparisons test in GraphPad Prism. (**E**) Relative expression of *SMYD5* mRNA compared to *GAPDH* from *SMYD5*-KO cells, measured by RT-qPCR at 37°C, with or without rescue with Flag-SMYD5 sgRNAres plasmid (labelled SMYD5 sgRNA#6res). Each data point is a biological replicate (n=4), with mean and SD where applicable. Significance levels were calculated with an unpaired one-tailed t-test in GraphPad Prism. (**F**) Western blot using antibodies against SMYD5 and Lamin B in a *SMYD5*-KO HEK293 cell line at 37°C with and without rescue with Flag-SMYD5 sgRNAres plasmid (labelled SMYD5 sgRNA#6res). *SMYD5*-KO was successful at the protein level at 37°C, (n=2-3). Data shown as in (E). (**G**) Relative expression of *SP1* mRNA compared to *GAPDH* from *SMYD5*-KO cells, measured by RT-qPCR at 37°C, with or without rescue with Flag-SMYD5 sgRNAres plasmid (labelled SMYD5 sgRNA#6res), (n=4). Data shown as in (E). (**H-J**) Western blot quantification with and without 16 h incubation at 32°C and representative examples using antibodies against SP1, CIRBP and RBM3, respectively, in *SMYD5*-KO cells. Each data point is a biological replicate (n=2-3), with mean and SD where applicable. Significance levels were calculated with an unpaired one-tailed t-test in GraphPad Prism. * = P<0.05, ** = P<0.01, *** = P<0.001, **** = P<0.0001.

### SMYD5 total protein levels are decreased at 32°C by the proteasome

To test whether SMYD5 itself shows temperature-dependent levels or distribution in cells, we performed cellular immunofluorescence staining. We observed nuclear and cytoplasmic staining of SMYD5 at 37°C (SMYD5: white or green, DAPI: purple, **Fig. 4Aa, Fig S6A-B left**) and at 32°C (**Fig. 4Ab, Fig S6A-B, right**). Quantified total protein expression of SMYD5 was decreased at 32°C compared to 37°C (P<0.05, **Fig. 4A-B** and **Fig. S6A**). We did not observe a significant difference in nuclear to whole cell SMYD5 mean intensity ratio at the two temperatures (**Fig. 4C**). This indicates that SMYD5 is not being sequestered outside of the nucleus in response to hypothermia. We further performed a Western blot against SMYD5^36^ that also showed decreased SMYD5 at 32°C (P<0.01, **Fig. 4D**). The difference seen between the two temperatures is not observed at the transcriptional level (**Fig. 4E**), indicating that SMYD5 is primarily regulated post-transcriptionally. Since SMYD5 is known to bind to some ubiquitin ligases per BioGRID (e.g. TRIM25), we inhibited the proteasome with MG132. With proteasomal inhibition we observed significantly increased SMYD5 levels at 32°C compared to non-exposed SMYD5 levels at 32°C (**Fig. 4F**). This supports the hypothesis that SMYD5 is actively recruited to the proteasome at 32°C. Finally, to demonstrate that this also takes place *in vivo*, we used sagittal brain slices (**Fig. 4G**) from neonatal mice (P10) injured with cerebral HI and treated with normothermia (37°C) or cooling (32°C)^37–39^. We observe a similar temperature-dependent decrease in SMYD5 but not NeuN after 6 h of cooling. This is observed both in the somatosensory cortex (**Fig. 4Hd**, **Fig. 4Hh** and **Fig. 4I**) and in the subiculum (Sub), CA1 and CA3 subfields of the hippocampus (**Fig. 4Ha-c, Fig 4e-g**, **Fig 4JG** and **Fig. S6C-D**). The temperature dependent behavior of SMYD5 indicates that SMYD5 might be a useful biomarker of adequate TTM in the clinical setting.

**Fig. 4.**
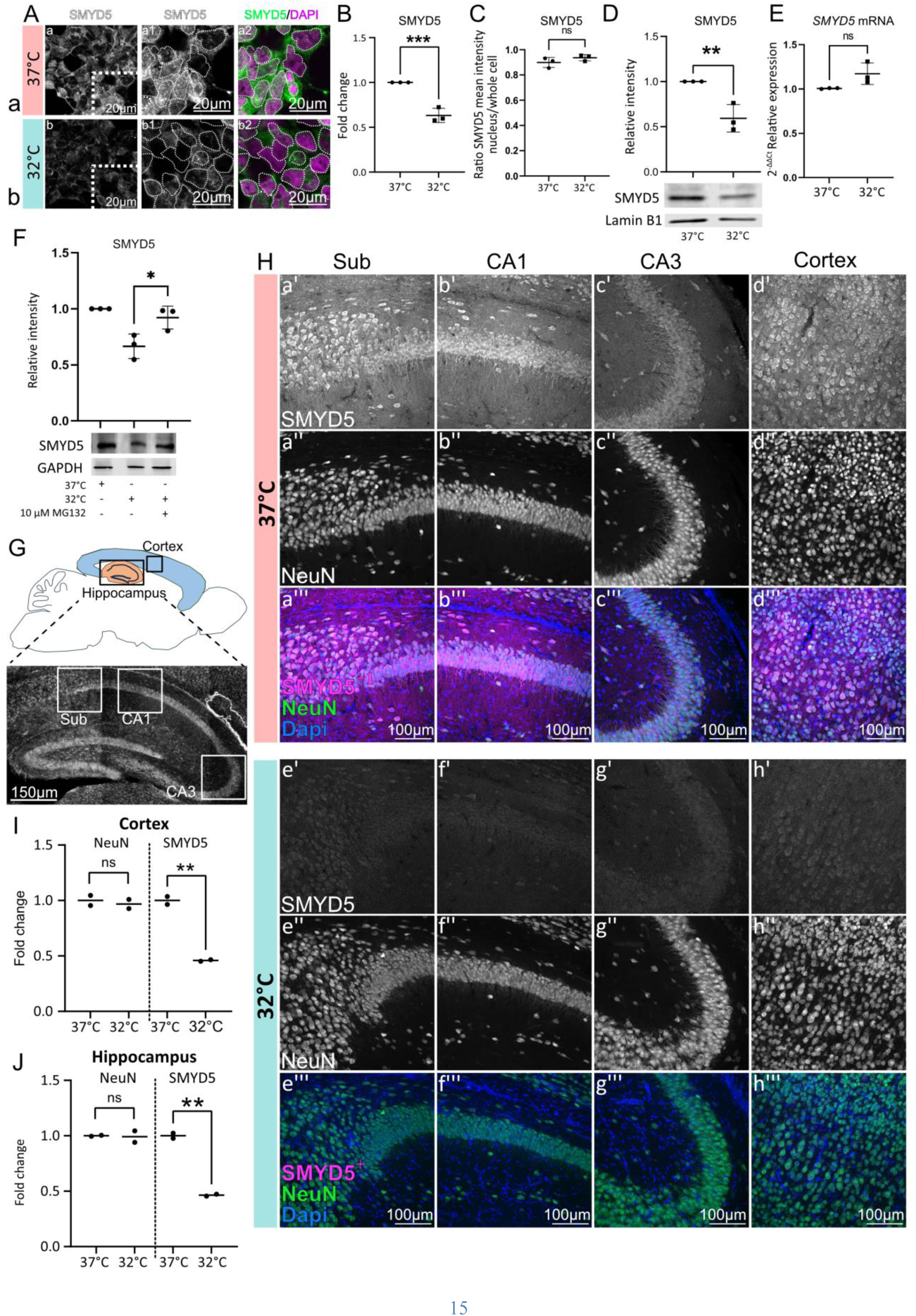
SMYD5 is depleted at 32°C *in vitro* by the proteasome and *in vivo*. (**A**) Representative image of intensity of endogenous SMYD5 at 37°C (top, a) and 6 h incubation at 32°C (bottom, b). Nuclei are marked with a dotted line. Other images can be found in **Fig. S6**. (**B**) Quantification of cellular endogenous SMYD5 mean intensity levels from immunocytochemistry, mean fold change between conditions is depicted. Each data point is a biological replicate (n=3), mean and SD shown where applicable, significance levels were calculated with an unpaired one-tailed t-test. (**C**) The ratio of nuclear to whole cell SMYD5 mean intensity, (n=3). Data shown as in (B) (**D**) Western blot using a SMYD5 antibody at 37°C and 6 h incubation at 32°C, (n=3). Significance levels calculated with an unpaired two-tailed t-test otherwise data shown as in (B). (**E**) qRT-PCR results for SMYD5 expression at 37°C and 32°C after 6 h incubation, (n=3). Significance levels calculated with an unpaired one-tailed t-test. (**F**) SMYD5 levels at 37°C and at 32°C with and without the exposure of the proteasomal inhibitor MG132, (n=3). Data shown as in B. (**G**) A schematic overview of sagittal sections of a P10 mouse brain. Boxes indicate areas that were used in (H-J). (**H**) Representative SMYD5, DAPI and NeuN staining in brain sections of mice kept at euthermia (37°C; a-d) after neonatal hypoxic-ischemic injury compared to those treated with cooling at 32°C, for 6 h (e-h). Sagittal brain sections are from the Sub (a and e), CA1 (b and f), CA3 (c and g) and somatosensory cortex (d and h). ^+^ Background for SMYD5 in merged images has been adjusted to emphasize the specific staining of SMYD5. (**I-J**) Quantification of NeuN and SMYD5 staining from (H) (n=2), where hippocampus included Sub, CA1 and CA3 regions. Each data point is a biological replicate, that is an average of two technical replicates, the mean of the replicates is shown. Mean fold change between conditions is depicted. Significance level calculated with an unpaired one-tailed t-test. All significance levels for this figure were calculated in GraphPad Prism, * = P<0.05, ** = P<0.01, *** = P<0.001, **** = P<0.0001.

### SMYD5 regulates additional mild hypothermia responsive genes

To identify additional genes that are both regulated by SMYD5 and responsive to mild hypothermia exposure, we performed RNASeq at the two temperatures from three sources: material from mouse neural progenitor cells (mNPCs), and on hippocampal and cortical tissues from mice (same mice as depicted and analyzed in **Fig. 4**). This yielded 5396 upregulated DEGs at 32°C, of which 348 were common in all three RNASeq datasets (**Fig. 5A**), and 4958 downregulated DEGs at 32°C, of which 617 were common in all datasets (**Fig. 5B**). This supports our hypothesis that the regulation of the MHR has a transcriptional component. We further overlapped upregulated DEGs (5396) at 32°C with SMYD5 repressed genes (106) at 37°C (SMYD5 bound and *Smyd5*-KO upregulated, **Fig. 3B**, left) and observe a significant overlap of 37 genes (P=5e-03, **Fig. 5C, table S3**). These 37 genes generally show upregulation at 32°C in all three datasets (**Fig. 5D**), and three were consistently significantly upregulated in all three RNASeq datasets (*Ewsr1*, *Hnrnpul1*, *Xrn2*).

**Fig. 5.**
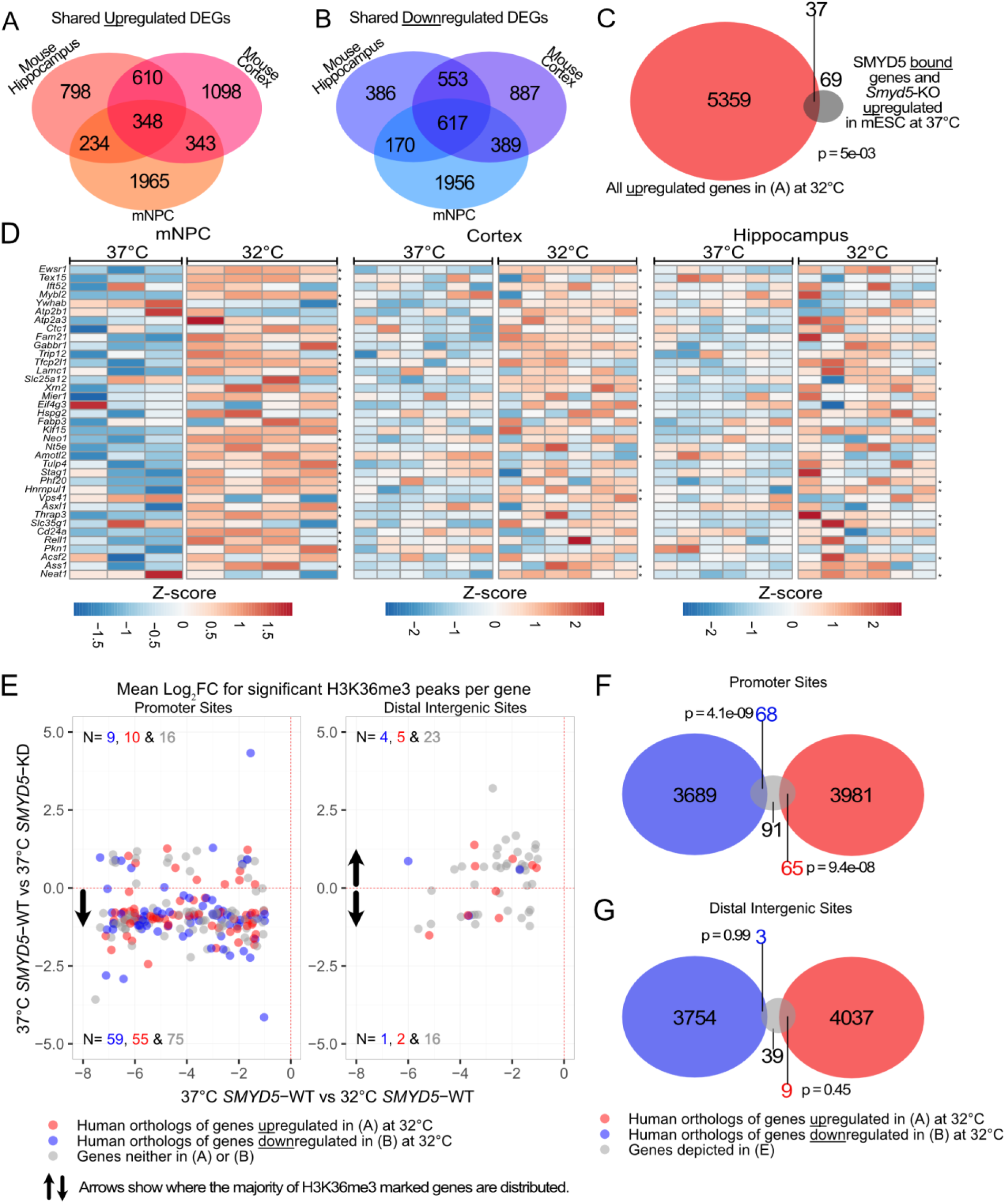
SMYD5 is a regulator of the MHR. (**A**) Overlap of upregulated DEGs *in vitro* and *in vivo* RNASeq datasets from mNPCs (n=3 at 37°C, n=4 at 32°C), hippocampal (n=6 at 37°C, n=6 at 32°C), and cortical cells (n=6 at 37°C, n=6 at 32°C), at 32°C compared to 37°C. (**B**) Overlap of downregulated DEGs *in vitro* and *in vivo* RNASeq datasets from (n=3 at 37°C, n=4 at 32°C), hippocampal (n=6 at 37°C, n=6 at 32°C), and cortical cells (n=6 at 37°C, n=6 at 32°C), at 32°C compared to 37°C. (**C**) Overlap of upregulated DEGs from *in vitro* and *in vivo* RNASeq datasets (mNPCs, hippocampal and cortical cells), at 32°C with SMYD5-bound and repressed genes in mESC. Statistical analysis was done with GeneOverlap^57^ package in R using Fisher’s Exact test. (**D**) Heatmap for the 37 genes from (C), depicting all normalized counts >10 converted to z-scores, the transformation was done with variance stabilization in DESeq2. *DEG in each dataset that have p. adjusted < 0.1 calculated with DESeq2. (**E**) Mean Log_2_FC of H3K36me3 peaks per gene over promoter sites and distal intergenic sites. Each datapoint depicted is a gene that had p value < 0.05 for both datasets and Log_2_FC < −1 for H3K36me3 modification when comparing 37°C to 32°C in *SMYD5*-WT cells (*SMYD5*-WT; n=6, *SMYD5*-KD; n=3). Red colored dots represent genes that are upregulated at 32°C in one of three RNASeq datasets presented in (A). Blue colored dots represent genes that are downregulated at 32°C in one of three RNASeq datasets presented in (B). Gray colored dots represent genes that are not upregulated at 32°C in any of the three RNASeq datasets presented in (A). (**F-G**) Overlap of genes from (E) and human orthologs of up- or downregulated genes at 32°C from at least one of the three RNASeq datasets from (A) or (B), at promoter sites and distal intergenic sites, respectively. Data shown as in (C).

To further explore the consequences on histone modification of differential SMYD5 availability at 37°C versus 32°C, we performed CUT&RUN using antibodies against H3K36me3, H4K20me3, H3K9me3 and H3K4me3 in human neural progenitor cells (NPCs) and in HEK293 cells. Only H3K36me3 and H3K4me3 show global differences between the two temperatures (**Fig. S7A-S7D**), so we performed CUT&RUN using antibodies against H3K36me3 and H3K4me3, with and without *SMYD5*-KD (**Fig. S7E**) in HEK293 cells (**Fig. S7C-S7D**). We observe a lower H3K4me3 level at 32°C compared to 37°C for both *SMYD5-*KD and -WT, which fits with previous observations of reduced protein production during cold^27^ and indicates that transcriptional response also play a role in this effect. Additionally, we see less H3K4me3 for *SMYD5*-KD compared to WT for both temperatures (**Fig. S7C**), indicating that SMYD5 also contributes to gene activation at both temperatures. In alignment with our prior data, we observe a decrease of H3K36me3 only at 37°C in *SMYD5*-KD of HEK293 cells compared to SMYD5-WT (**Fig. S7D**), indicating loss of H3K36me3 levels that are normally maintained at 37°C. This supports the hypothesis that SMYD5 regulates a subset of hypothermia responsive genes at 37°C. On the individual gene level, we observe many genes that have either increased or decreased H3K36me3 or H3K4me3 at 32°C compared to 37°C both for *SMYD5*-WT and -KD (**Fig. S7F-S7I**). Further examination of the peak distribution over *SP1* and *EWSR1* promoters revealed lower density of the H3K36me3 modification with *SMYD5*-KD at 37°C (**Fig. S7J**). The lack of complete loss of the H3K36me3 modification for *SMYD5*-KD could be due to compensation by other histone methyltransferases, a previously described phenomenon^40^. Next, we examined the mean Log_2_FoldChange (Log_2_FC) of H3K36me3 signal over promoter regions for each gene with significantly differential decreased H3K36me3 modifications at 32°C. When comparing this metric for two datasets, *SMYD5*-KD versus -WT at 37°C and *SMYD5*-WT at 32°C vs 37°C, we observe that 84% of the genes with decreased H3K36me3 at 32°C in *SMYD5*-WT also have decreased H3K36me3 with *SMYD5*-KD at 37°C (**Fig. 5E, left**). In contrast, we do not observe this same phenomenon for other regions such as distal intergenic regions (**Fig. 5E, right**). This indicates that *SMYD5*-KD at 37°C induces a H3K36me3 profile that resembles the profile seen in mild hypothermia, and that this effect is primarily limited to promoters. Fewer genes were associated with H3K36me3 peaks at distal intergenic sites, which is expected as SMYD5 is known to deposit H3K36me3 at promoters (*30*). We then compared the list of the 224 genes with significantly decreased H3K36me3 over their promoter sites to the human orthologs of the DEGs detected at 32°C (**Fig. 5A**). We observe a significant overlap of 65 upregulated (red) and 68 downregulated (blue) DEGS at 32°C in at least one of the RNASeq datasets (P=9.4e-08 and P=4.1e-09, respectively, **Fig. 5E, left** and **5F).** When examining the same overlap for our list of genes associated with decreased H3K36me3 over distal intergenic regions (51 genes), we observed a non-significant overlap of only 9 upregulated 3 downregulated DEGs (P=0.45 and P=0.99, respectively, **Fig. 5E, right** and **5G**). This is consistent with the results from the independent mESC dataset (**Fig. 3B-3C**). It is also congruent with the observed global decrease in H3K36me3 following *SMYD5*-KD at 37°C in HEK293 cells, and the global decrease in H3K36me3 at 32°C compared to 37°C in hNPCs (**Fig. S7D**). Together these data indicate that the SMYD5-dependent H3K36me3 signal at promoters mediates the differential expression of a subset of MHR genes at 32°C.

## Discussion

Environmental responses are likely to be dynamic and our MHIs allow for some temporal dissection. Currently, all MHIs harbor stable fluorescent protein, which may be particularly sensitive to capturing changes over time rather than dynamic differences at a given time point. The latter can be detected by RT-qPCR and Western blotting. Thus, our current MHIs may be most useful to map the start of the response but the exact timing of the response itself may be a key task for future studies.

Our data support the hypothesis that SP1 and RBM3 may be parts of distinct arms of the MHR as SMYD5 appears to regulate *SP1* at 37°C but not *RBM3*. We interpret the lag in the response (at least 6 hours to signal) and the obvious shift in fluorescence of sgRNA mutagenized cells to mean that there are likely many genes upstream of the currently tested point of the response (*SP1*). We provide 134 candidate regulators that can subsequently be validated by the scientific community using our MHIs or other strategies. By uncovering *SMYD5* as a repressor of *SP1* at 37°C, a key factor of the mammalian MHR, we provide an example of how histone methylation integrates temperature cues into a fundamental mammalian response. We suggest that SMYD5 exerts its effects at 37°C to ensure that *SP1*/*CIRBP* are not upregulated without a hypothermic stimulus. However, at onset of hypothermia (32°C), SMYD5 is rapidly degraded by the proteasome and SMYD5 repression is released, thus activating the MHR. A generalized decrease of protein production occurs after a cold stimulus^27^, and thus decreased translational efficiency could play a part in decreasing SMYD5 levels. However, we are unaware of biological examples of targeted protein degradation by the proteasome in response to mild hypothermia.

SMYD5 itself appears to regulate at least 37 other genes that show temperature dependence in which SMYD5 acts as a repressor (like *SP1*). There were three genes: *Ewsr1*, *Hnrnpul1* and *Xrn2* that were significantly upregulated at 32°C in all three RNASeq datasets and repressed by SMYD5 at 37°C. All three play a role in RNA-processing like CIRBP and RBM3 and have a predicted SP1-binding sites similar to *Sp1* itself (per GeneHancer in Genecards). *HNRNPUL1* is also on our list of SP1 activators (**Table 2**). There are other interesting genes on our list (**Table 3**) which includes *Gabbr1*, a gene which when knocked out leads to decreased basal body temperature in mice^41^. The list includes genes that are known to play a role in tissues of physiological mammalian temperature responses such as the thyroid (*TRIP12*, *THRAP3,*^42^), brown adipose tissue (*KLF15,*^43^) and testes (*TEX15,*^44^).

In summary, we have developed and validated tools to interrogate the MHR and used them in combination with cutting-edge technology (CRISPR-Cas9 library screen) to provide a list of 134 candidate MHR regulators. We validate SMYD5 as a temperature-dependent histone modification regulator of the MHR and show that loss of histone methylation (H3K36me3) at specific gene sites in mammals happens with MHR exposure. Our results provide a specific description of how mammalian cells use histone modification to respond to temperature cues with some evidence that this phenomenon may translate to in vivo following cooling treatment for neonatal hypoxic-ischemic brain injury.

### Limitations of Study

A caveat of our study is that our results are not fully transferable to neonatal hypoxic-ischemic brain injury as neuroprotection in that setting is achieved at 33.5°C and getting as low as 32°C is not recommended^10^. Another limitation of our study is that SMYD5 doesńt have any known intrinsic DNA binding ability^45^, thus there must be another factor that recruits SMYD5 to the right sites. This could be addressed in future studies by exploring protein interactors of SMYD5 or with a more focused CRISPR-Cas9 library screening using a library only containing known transcriptional regulators. Another limitation of this study is that we do not know the mechanism of how SMYD5 is labeled to be routed to the proteosome with mild hypothermia. This would be important to address in future studies to fully understand how the MHR repression by SMYD5 is lifted under mild hypothermia.

## Acknowledgments

We thank SCRU lab (UI) for gifting us pMD2.G (Addgene, #12259) and psPAX2 (Addgene, #12260), and H.C. Dietz and D. Valle labs (JHU) for gifting us the HCT-116, SK-N-SH, HeLa, Jurkat and K562 cell lines. We thank the Immunology department of Landspitali University Hospital for the access and assistance with the FACS sorting. We thank FUNlab at deCODE genetics for the usage of their flow cytometry machines as well as for gifting us the HEK293T cell line. We thank Kári Stefánsson and Ólafur Þ. Magnússon at deCODE Genetics for providing some of the next generation sequencing for this project. We thank Rachel Latanich for help with preparing isolating RNA for the *in vivo* RNA-Seq studies. Finally, some images in this manuscript were created with BioRender.com. Funding: Rannis Technology Development Fund (#2010588, HTB), Eimskip University Fund (SR), Fulbright Iceland (SR), Icelandic Cancer Society Research Fund (HTB), Gongum Saman Research Fund (SR), Landspitali Research Fund (HTB), Felag Haskolakvenna Research Fund (SR), Rannis (#217988, #195835, #206806, HTB), Thomas Wilson Foundation (RC-V), Johns Hopkins University Pediatric Innovation Grant (RC-V), NIH R01NS126549 (RC-V, FJN), 1R21NS123814 (RC-V, FJN), 1R01 HD110091 (RC-V, FJN), R01HD074593 (FJN), U01NS114144 (FJN)

## Author contributions

Conceptualization: HTB, SR; Methodology: HTB, SR, LZ, KJA; Software: SR, KJA, KM, STH, TR; Validation: SR, KJ, LHA, AT, MV, KO; Formal analysis: SR, KJA, KM, TR, KJ, STH, MV, EM; Investigation: SR, KJA, KJ, AT, MV, STH, LHA, RC-V, FJN, EM, TR; Resources: JH, EM, KEA; Data curation: SR, KJA, KM, TR, KJ, MV; Writing – original draft: HTB; Writing – review & editing: HTB, SR, KJA, RC-V; Visualization: SR, KJA, KM, EM, KJ, TR, HTB, AT, MV; Supervision: HTB; Project administration: HTB, SR, KJA; Funding acquisition: HTB, SR, RC-V, FJN.

## STAR★Methods

### Key resources table

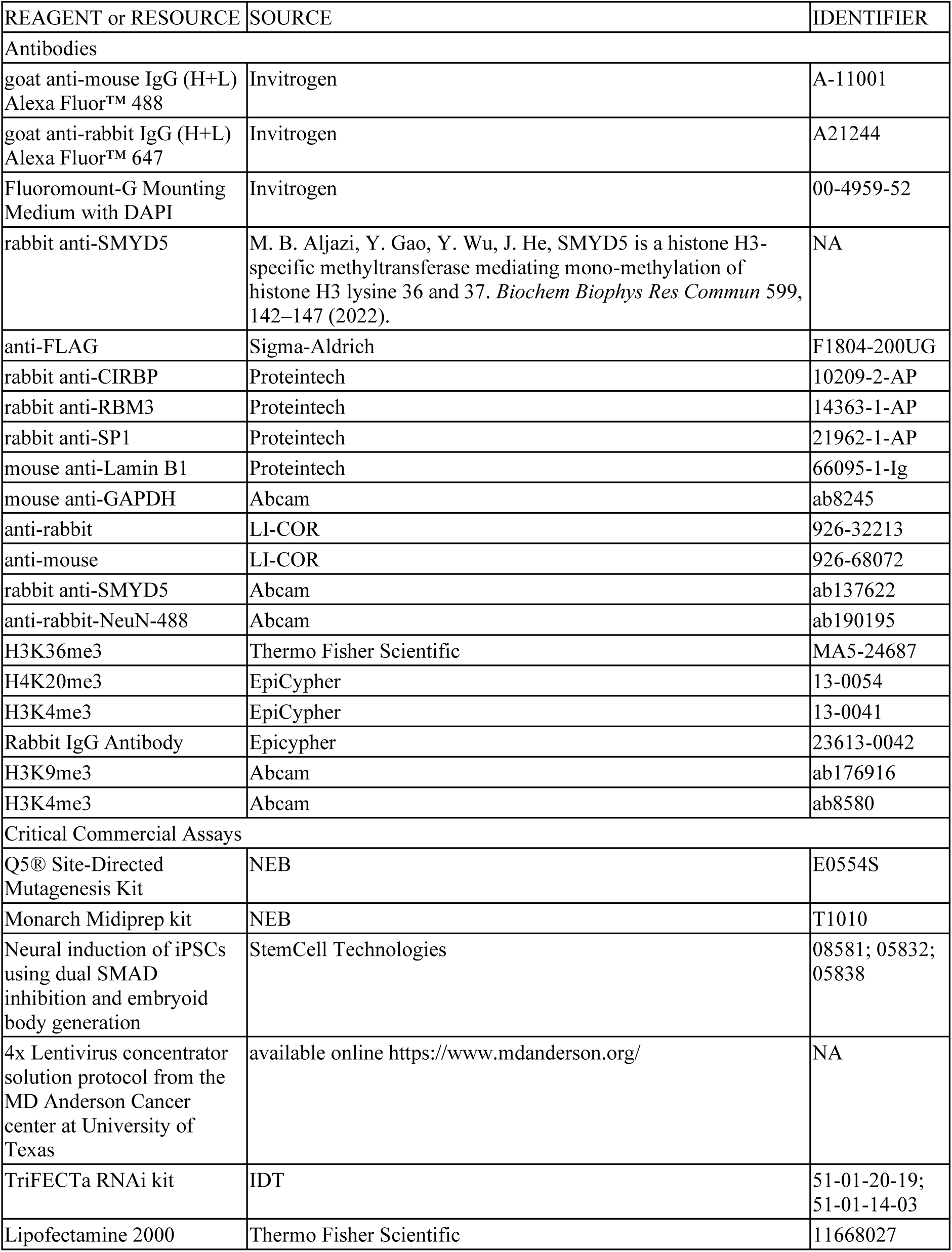
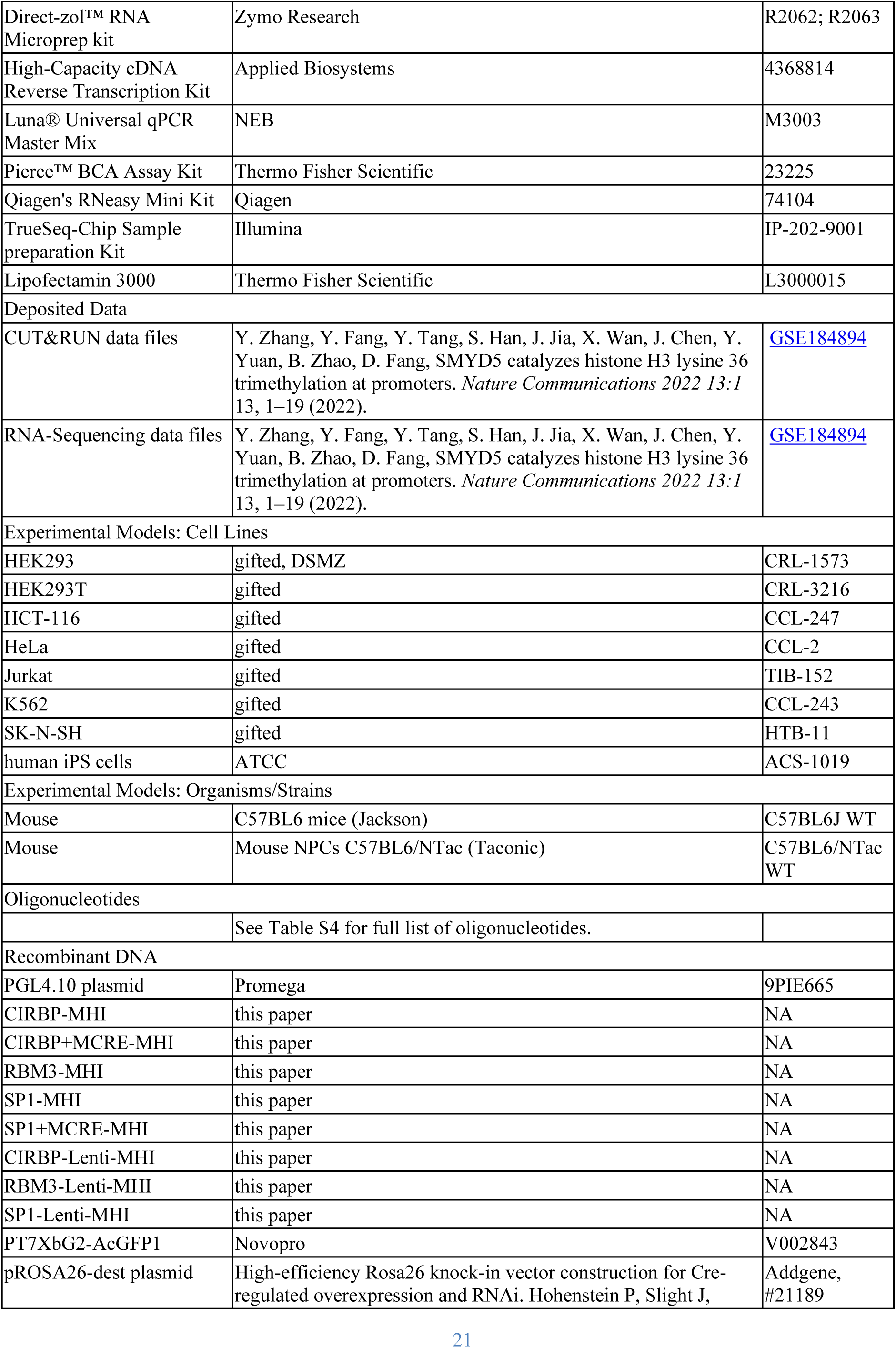
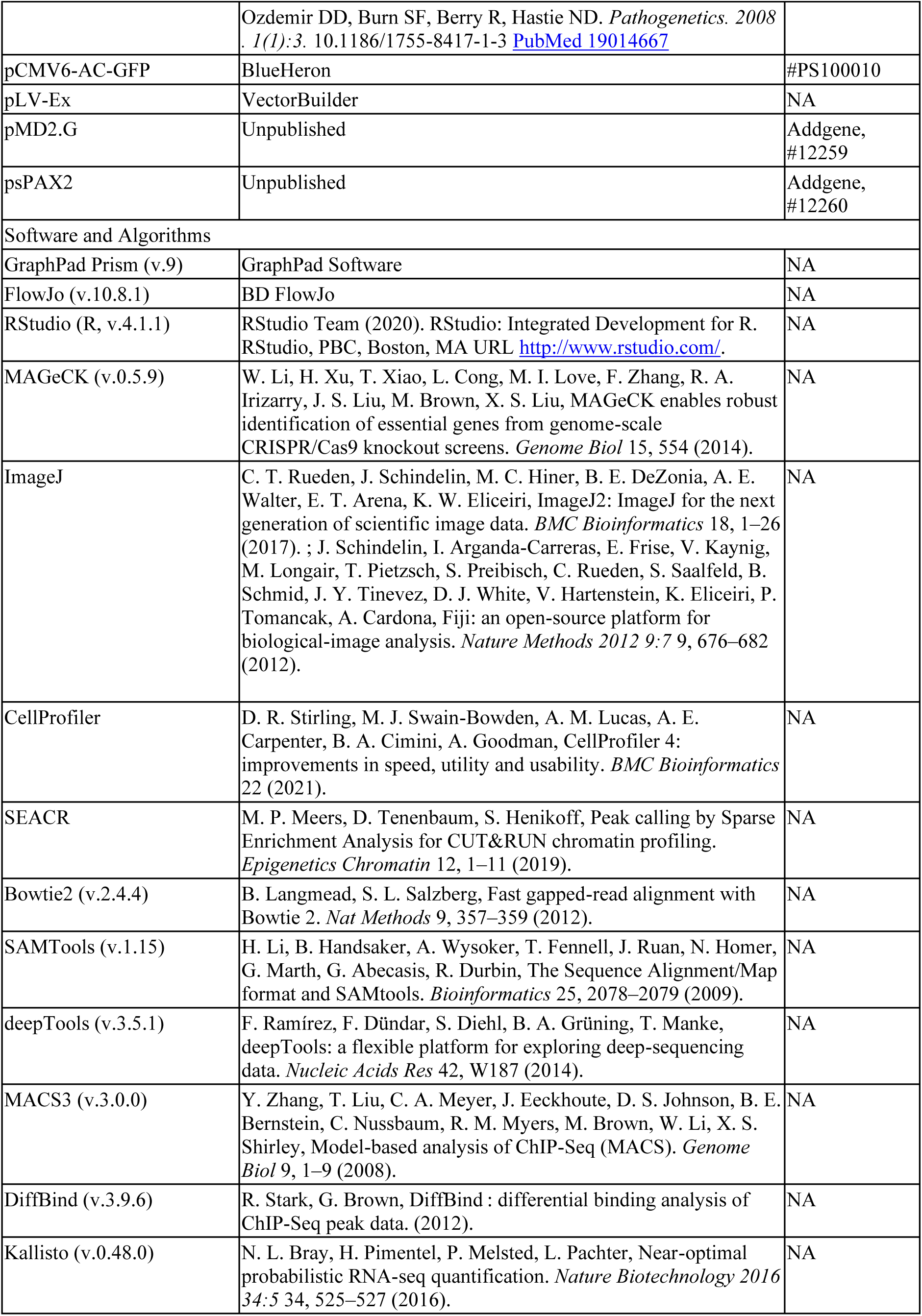

### Resource availability

#### Lead contact

Further information and requests for resources and reagents should be directed to and will be fulfilled by the lead contact: Hans Tomas Bjornsson

#### Materials availability

MHI are available upon request through MTA. All sequencing data are available through Gene Expression Omnibus (GSE234699). All other data are available in the main text or the supplementary materials.

#### Data and code availability

This paper does not report any original code.

### Experimental model and study participant details

We did not exclude mice from this study based on gender. We used mouse strains: C57BL6J WT, C57BL6/NTac WT as well as cell lines: HEK293, HEK293T, HCT-116, HeLa, Jurkat, K562, SK-N-SH, human iPS cells. For further information see methods details section and key resource table.

### Method details

#### Creation of indicators

We used the PGL4.10 plasmid (Promega, #9PIE665) as the backbone for our CIRBP- and SP1-MHI, where the *luc2* sequence was cut from the plasmid with NcoI (NEB) and XbaI (NEB). Promoter sequence for *CIRBP*^46^ was cloned from mouse DNA using *CIRBP* forward primer 5’-tcgataggtaccTGGCTTCACAAATGCGCCTCAGT-3’ and *CIRBP* reverse primer 3’-cctaaggcagatctGCGAGGGGGAGCGCAAGAGT-5’. Restriction enzymes KpnI (NEB) and BglII (NEB) were used to insert the promoter into the plasmid. Promoter sequence for *SP1* ^47^ was cloned from human DNA using the forward primer 5’-tcaagtcaggctagcGCAACTTAGTCTCACACGCCTTGG-3’ and reverse primer 3’-cagtgctgcctcgagGCTCAAGGGGGTCCTGTCCGG-5’. Restriction enzymes NheI (NEB) and XhoI (NEB) were used to insert the promoter into the plasmid. AcGFP1 was cloned from PT7XbG2-AcGFP1 (Novopro, #V002843) vector using the reverse primer 3’-cggcggagTCTAGAATTACTTGTACAGCTCGTCC-5’ and the forward primer 5’-taagccaccATGGTGAGCAAGGGCGAGGAGC-3’. Restriction enzymes NheI (NEB) and XhoI (NEB) were used to insert the acGFP1 into the plasmid. The neomycin selection cassette was cloned from pROSA26-dest plasmid (Addgene, #21189) using the forward primer 5’-CATTATCGTCGACTCTACCGGGTAGGGGAGGCGCTT-3’ and reverse primer 3’-CGCCGCCGACGATAGTCAAGCTTCTGATGGAATTAGAACTTGGC-5’. Restriction enzymes SalI (NEB) and PshAI (NEB) were used to insert the neomycin cassette into the plasmid. CIRBP- and SP1-MHI were made with and without MCRE^18^. The MCRE sequence was inserted in 5 repeats in front of the promoter sequences with a linker sequence between. Acc65I (NEB) restriction enzyme was used to insert the MCRE sequence into the CIRBP-MHI and Acc65I (NEB) and SacI (NEB) restriction enzymes were used to insert the MCRE into the SP1-MHI. Sanger Sequencing was used to validate insertion sites, size, and sequence of insertion. The *RBM3* promoter with MCRE enhancer (6 repeats) in front of the promoter was cloned into the multiple cloning site pCMV6-AC-GFP (BlueHeron, #PS100010). We also made lentiviral vectors for *CIRP*, *SP1* and *RBM3* on a pLV-Ex (VectorBuilder) lentiviral vector. The *CIRBP* promoter for the CIRBP-lenti-MHI was designed from the human promoter and the *SP1* promoter was moved upstream of the TSS for the SP1-lenti-MHI but *RBM3* promoter sequence remained unchanged. Included were MCRE (8 repeats for CIRBP-lenti-MHI, five repeats for other), promoter, neomycin selection cassette and a mCherry fluorescent protein behind each promoter sequence. The promoter region, size and backbone plasmid for each indicator is detailed in **Table S1** and sequence of each MHI in **File S1**. We also observed some variation in absolute levels among individual indicators/experiments, but the pattern was always consistent; given this, we only show data from individual experiments in any figure.

#### Culture of cell lines

HEK293 (gifted and DSMZ, CRL-1573, referred to as 293/293WT) cells were cultured in DMEM/F12 with GlutaMAX (Gibco, 10565018) with 10% filtered FBS (Gibco, 10270106) under 5% CO_2_. Cells were cultured at 37°C unless otherwise stated, then 37°C served as a control. For experiments at 32°C and 37°C, we split the cells into two plates, and cells were kept at 37°C until 16 h before flow cytometry then plates were kept at each temperature for comparison of fluorescence to account for any build-up of fluorescence over time. In addition to HEK293, we used the following cell types HEK293T (gifted, CRL-3216), HCT-116 (gifted, CCL-247), HeLa (gifted, CCL-2), Jurkat (gifted, TIB-152), K562 (gifted, CCL-243), SK-N-SH (gifted, HTB-11). Cells were sub-cultured when they had reached 70-90% confluence and media was changed every other day. Killing curves were used for all cell lines to determine the lowest selection concentration for all selection agents. We used culture recommendations from ATCC when possible. We also used human iPS cells (ATCC, ACS-1019) that were used to generate NPCs by neural induction of iPSCs using dual SMAD inhibition and embryoid body generation (StemCell Technologies; 08581, 05832, 05838), prior to experiments described here the NPCs were cultured in STEMdiff Neural Progenitor Medium (StemCell Technologies, 05833) on Matrigel (Corning, 354234). As well as murine primary NPCs (see section mNPC isolation for further information).

#### Transfection of MHI and SMYD5-Flag overexpression plasmid

HCT116, 293WT, HEK293T, HeLa and SK-N-SH cell lines were transfected with MHI when they had reached 70-90% confluency. We used either Lipofectamine^TM^ 2000 or 3000 (Thermo Fisher Scientific, 11668027, L3000015) according to the manufacturer’s protocol. For transfection of K562 and Jurkat cell lines we used electroporation for transfecting the MHIs into the cell lines. The fluorescence of the indicators was measured via flow cytometry up to 48 h after transfection. We analyzed flow cytometry data with the FlowJo Software, where we gated living cells, single cells, GFP positive cells and measured the fluorescence for that population. For further analysis, we used Excel from Microsoft, GraphPad Prism (v.9) and FlowJo (v.10.8.1). For transfection of SMYD5-Flag overexpression vector^30^ we used the same lipofectamine method as described above.

#### Construction of 293WT+Cas9 and 293WT+Cas9+SP1 cell lines

We made two cell lines from 293 cells, one that expressed Cas9 (lentiCas9-blast, Addgene #52962), referred to as 293WT+Cas9, and another that expressed both Cas9 and SP1-MHI, referred to as 293WT+Cas9+SP1. These cell lines were made by transfecting the plasmids with lipofectamine as described above and selecting stably transfected cells using Blasticidin (Thermo Fischer Scientific, A1113903) selection for Cas9 and Neomycin(G418/Geneticin) (Santa-Cruz, sc-29065A) selection for SP1-MHI.

#### Apoptosis assay

We based our apoptosis assay on a protocol published by Xiang et al.^48^. We exposed 293WT+Cas9+SP1 cell line either to 1-5 µM H_2_O_2_ or vehicle (H_2_O) for 4 h at 32°C or 37°C. 4 h after exposure gMFI of the cells were analyzed with flow cytometry as described above.

#### Genome Wide CRISPR-Cas9 screen on fluorescent HEK293WT cell lines that stably express SP1-MHI

Using the 293WT+Cas9+SP1 and 293WT+Cas9 (negative control) cell lines we performed a genome wide CRISPR-Cas9 knockout screen (GeCKO) screen, where we transduced the sgRNA pool made from library A and B (Addgene, #1000000049) in MOI 0.3^22–24^. We mostly followed the Joung et al. protocol ^22^, the exceptions are as follows. For making the lentivirus, a protocol described by Kutner et al.^49^ was used, using pMD2.G (Addgene, #12259) and psPAX2 (Addgene, #12260). Lentiviral sgRNA library concentration was done with Amicon ultracentrifugal filters (Millipore, UFC9003). 20 h after transduction with the sgRNA library selection was started with Puromycin and continued for 6 days and 8 h before the cells were moved to 32°C, where they were incubated for 16 h before fluorescent activating cell sorting (FACS) with Cell sorter SH800Z (Sony). The 5% highest fluorescent cells for 293WT+Cas9+SP1 and 5% lowest fluorescent cells for 293WT+Cas9+SP1 were sorted. Next, we isolated gDNA and performed a two-step PCR for all samples (sorted as well as negative control). According to cell number after puromycin selection, the sgRNA library coverage was >700 times coverage for HEK293WT+Cas9+SP1. Next generation sequencing (NGS) was performed with single read, 80 cycles and 8 indexing cycles with PhiX spike in of 20% on NovaSeq 6000 S4 at deCODE genetics for HEK293WT+Cas9+SP1 and HEK293WT+Cas9, 32-280 million aligned reads per sample. To analyze the screen data, single read fastq files for each replicate and condition were merged using cat command. Then, MAGeCK (v.0.5.9, ^29^) was used to identify enriched sgRNA’s in sorted samples and sgRNA counts were normalized to internal control for sgRNA’s (control sgRNAs). We used RStudio (R, v.4.1.1, ^50^) to visualize the results and to filter genes. We exported a list of genes that had both FDR value under 0.25 and LFC value over 3.5 for each condition in both screens. Then, we excluded genes, overrepresented in both the repressor and activators screen when compared to control as they most likely promote cell growth and therefore were present in both lists (*CEP250*, *FAM102B*, *hsa-mir-3944*, *HOXB4*, *hsa-mir-6729*, *TRMT6*, *METTL9*, *SH3BPI*, *VSIG10L*, *MALT1*, *NLRP12*, *hsa-mir-5699*, *LZTR1*, *SAG*, *RGS21*, *FUBP1*, *PTMA*, *hsa-mir-6818*, *CCL4*, *CSRP2BP*, *C14org178*). Two of these genes (*MALT1* and *RGS21*), were validated and found to not show any impact on indicators (data not shown).

#### Making of *SMYD5* knockout (*SMYD5*-KO) cells

Two ready-made sgRNA vectors (Genscript, *SMYD5* plasmid vector #2 and #6) were packed into lentivirus according to the protocol described by Kutner et al.^49^, using pMD2.G (Addgene, #12259) and psPAX2 (Addgene, #12260). For concentration we used the 4x Lentivirus concentrator solution protocol from the MD Anderson Cancer center at University of Texas (available online at https://www.mdanderson.org/). We selected stable *SMYD5*-KO cells with Puromycin selection for at least 7 days.

#### Making of *MALT1* and *RGS21* knockout cells

Ready-made sgRNA vectors (Genscript, plasmid vector *MALT1* #1 and #2, *RGS21* plasmid vector #2 and #3) were packed into lentivirus and selected as described above in the making of *SMYD5-KO* cells.

#### Making of gRNA resistant SMYD5-Flag plasmid (Flag-SMYD5 sgRNAres plasmid)

In an effort to make SMYD5-Flag resistant to CRISPR-Cas9 cutting from the two guides (Genscript, *SMYD5* plasmid vector #2 and #6), we used site-directed mutagenesis (Q5® Site-Directed Mutagenesis Kit from NEB, E0554S) to modify predicted PAM sites of the guides. We used the following primers to induce the mutations: SMYD5-2mut-F: 5’-gctctttacgAGGAAGCAGTCAGCCAGT-’3; SMYD5-2-mut-R: 5’-ttccgtaaagAGTCTCCGCAGAAGTTCC-’3; SMYD5-6-mut-F: 5’-accgatatcgAGCCTGTGACCACTGCCT-’3; SMYD5-6-mut-R: 5’-atagagcgtt-CCAGAGAAACTGTGCAGCC-’3. After site-directed mutagenesis, we isolated plasmid with Monarch Midiprep kit (NEB, T1010) and Sanger sequenced to verify changes in PAM-sites. This plasmid was then used for rescue experiments (**Fig. 3**).

#### SMYD5 siRNA knockdown for fluorescent analysis of SP1-MHI

We used the TriFECTa RNAi kit (IDT; TYE 563 Transfection Control DsiRNA, 51-01-20-19 (transfection control); Negative Control DsiRNA, 51-01-14-03 (negative control)) and used predesigned DsiRNA (IDT, hs.Ri.SMYD5.13.8) to knockdown *SMYD5* mRNA in three biological cell lines of HEK293+Cas9+SP1-MHI. For the reverse transfection we used Lipofectamine 2000 (Thermo Fisher Scientific, 11668027) where we seeded 1*10^6^ cells per well in a 6 well plate and exposed to siRNA. Around 24 h after the reverse transfection one batch of cells was exposed to 32°C and the other kept at 37°C for 6 hours before the cells underwent flow cytometry. Further statistical analysis of the flow cytometry data was similar to that described above.

#### Immunofluorescence of cell culture

HEK293 cells were seeded into two 8-well chamber slides (Falcon, 354118) and cultured for 24-48 h before one of the chamber slides was moved to 32°C. After 6 h cells from both slides at 32°C and 37°C were fixed and permeabilized with 4% PFA and 0.1% Triton-X 100 diluted in PBS for 10 minutes at room temperature. Then cells were washed twice with a blocking solution (PBS with 2.5% BSA and 10% normal goat serum) and followed by blocking for 30 minutes at room temperature in the blocking solution. Cells were incubated overnight with primary antibodies (rabbit anti-SMYD5, 1:1000, custom SMYD5 antibody has previously been validated by knocking out SMYD5 and observing the disappearance of the SMYD5 50 kd band^36^; anti-FLAG, 1:500, (Sigma-Aldrich, F1804-200UG)) at 4°C. The following day, cells were incubated at room temperature for 30 minutes and then washed three times every 10 minutes with a washing solution composed of PBS with 0.25% BSA. Cells were incubated with secondary antibodies (goat anti-mouse IgG (H+L) Alexa Fluor™ 488, 1:1000 (Invitrogen™, A-11001); goat anti-rabbit IgG (H+L) Alexa Fluor™ 647, 1:1000 (Invitrogen™, A21244)) for 1 h at room temperature in the dark. Cells were washed three times every 10 minutes with the washing solution and mounted with Fluoromount-G Mounting Medium with DAPI (Invitrogen, 00-4959-52). Cells were imaged using confocal microscopy (FV 1200, Olympus Fluoroview) using a 30x silicon objective (NA1.05). Images were processed by ImageJ ^51,52^ and quantified using CellProfiler (v.4.2.5.,^53^). Statistics were calculated in GraphPad Prism.

#### Real Time Quantitative Polymerase Chain Reaction (RT-qPCR)

Total RNA from cells incubated at 32 or 37°C was isolated, using Direct-zol™ RNA Microprep (Zymo Research, R2062) according to the manufacturer’s instructions. The concentration of RNA was measured by NanoDrop (Thermo Fisher Scientific) followed by cDNA synthesis with High-Capacity cDNA Reverse Transcription Kit (Applied Biosystems, 4368814) on MiniAmp™ Thermal Cycler (Applied Biosystems™). RT-qPCR was performed using Luna® Universal qPCR Master Mix (NEB, M3003) on CFX384™ Real-Time PCR Detection System (Bio-Rad). Each biological replicate of the RT-qPCR assay in this study was carried out in technical triplicates. Any technical replicate that deviated from other replicates by ≥ 0.4 cycle threshold (Ct) was removed from calculations of average Ct values. The primers used are as follows: SP1 (fwd): 5’-CACCCAATTCAAGGCCTGCCGT-3’; SP1 (rev): 5’-GGGTTGGGCATCTGGGCTGTTT-3’; RBM3 (fwd): 5’-GAGACTCAGCGGTCCAGGGGTT-3’; RBM3 (rev): 5’-CCTCTGGTTCCCCGAGCAGACT-3’; CIRBP (fwd): 5’-CCGAGTTGACCAGGCTGGCAAG-3’; CIRBP (rev): 5’-TCCATAGCCCCGGTCTCCTCCT-3’; GAPDH (fwd): 5’-TCAAGGCTGAGAACGGGAAG-3’; GAPDH (rev): 5’-CGCCCCACTTGATTTTGGAG-3’. SMYD5 1 (fwd): 5’-GCACTGTGCGCAAAGACCTCCA-3’, SMYD5 2 (fwd): 5’-GGAAACCAGGCCAGGTTCTGCC-3’, SMYD5 3 (fwd): 5’-CGTGGAAGTCCGTTTCGTGA-3’, SMYD5 1 (rev): 5’-CTGGGCACAGGACCTGGTGGTA-3’, SMYD5 2 (rev): 5’-GGCTGCCAACCGACATTCTGCA-3’, SMYD5 3 (rev): 5’-CCAGAGAAACTGTGCAGCCA-3’.

#### Western blot assay

Cells were washed with PBS and then lysed (whole cell lysate) for 30 minutes on ice in RIPA buffer (50mM Tris HCl ph8, 150mM NaCl, 1% NP-40, 1% Sodium deoxycholate, 0.1% SDS, 2mM EDTA, phosphatase inhibitor (either Cell signaling, 5870S or Thermo Fisher Scientific, 78437)). The lysate was centrifuged at 16,000 x g for 20 minutes at 4°C and the supernatant was collected. The supernatant was diluted with 4X loading buffer (LI-COR, 928-40004) and heated at 95°C for 5 minutes. The protein concentration was measured using Pierce™ BCA Assay Kit (Thermo Fisher Scientific, 23225). 20-30 µg of protein was loaded into each well and separated by SDS-PAGE. They were transferred to polyvinylidene fluoride (PVDF) membranes and blocked in 5% bovine serum albumin in Tris-buffered saline with Tween 20 for 1 h. Primary antibodies: rabbit anti-CIRBP, 1:2000 (Proteintech, 10209-2-AP); rabbit anti-RBM3, 1:1000 (Proteintech, 14363-1-AP); rabbit anti-SP1 1:1000 (Proteintech, 21962-1-AP); mouse anti-Lamin B1, 1:5000 (Proteintech 66095-1-Ig); rabbit anti-SMYD5, 1:1000, custom SMYD5 antibody has previously been validated by knocking out SMYD5 and observing the disappearance of the SMYD5 50 kd band^36^; mouse anti-GAPDH, 1:5000 (Abcam, ab8245) were applied to respective membranes after washing, and incubated at 4^°^C overnight. Membranes were incubated with IRDye® secondary antibodies; anti-rabbit (LI-COR,926-32213); anti-mouse (LI-COR, 926-68072), at room temperature for 90 minutes. After washing, the protein bands were visualized on the Odyssey® CLx Infrared Imaging System and protein band intensity quantified by Image J^51,52^ software.

#### Proteasome inhibition of SMYD5 using MG132

1,000,000 HEK293 cells were seeded in each well of two 6-well plates. When the cell confluency reached around 70%, 10 μM MG132 (Santa Cruz, SC-201270) was added to the cells, while the same amount (µL) of DMSO (Santa Cruz, SC-358801) was added to the control cells. One plate was then moved to a 32°C incubator, whereas the other plate was kept at 37°C for 6 hours. After the 6-hour incubation, cells were lysed following the procedures described above in the western blot assay chapter. The primary antibody rabbit anti-SMYD5 1:1000 (Abcam, ab137622) and rabbit anti-SMYD5, 1:1000^36^, was used for detection of SMYD5 in the same way as described above.

#### Mouse cooling followed by RNASeq and immunostaining on brain slices of cortex and hippocampus

C57BL6 mice (Jackson) were bred to create a litter (n=2). At P10, we split littermates into two groups. One group (n=6) was cooled to a core temperature of 32°C and other (n=6) were maintained at 37°C, for 6 hours respectively. These animals were then euthanized, and their brains were isolated immediately after the cooling period and flashed with PBS. One of the hemispheres was immersed in 4% PFA for 72-96 h for fixation and the other hemisphere was dissected for the cerebellum, thalamus/basal ganglia, hippocampus, anterior and posterior cortex. After fixation, samples were cryopreserved in a sucrose gradient and then flash-frozen. RNA was isolated using Qiagen’s RNeasy Mini Kit (Qiagen, 74104). Isolated RNA was constructed with directional mRNA library preparation (poly A enrichment) and then sequencing was performed by Novogene on a NovaSeq PE150 paired-end, generating over 80 million reads per sample. Analysis of the RNAseq was performed in the same manner as described in the “RNASeq mNPC” chapter below. Frozen hemispheres were cryosectioned with the help of a cryostat (Keldur, UI) and kept at - 20°C until stained. The sections were washed in TBS-T (TBS (LiCor, 927-60001) with 0.025% triton X-100 (Sigma-Aldrich, T8787-100ML)) 3x for 10 minutes. Antigen retrieval was performed using a sodium citrate buffer, pH 6, at 95°C for 20 minutes. Sections were washed in TBS-T 3x for 10 minutes and blocked in blocking buffer (TBS-T+ with 3% normal goat serum (NGS (Abcam, ab7481) and 0.1M Glycine (Sigma)) for 1 h at room temperature. The sections were stained with anti-rabbit-SMYD5^36^ primary antibody at 1:200 in TBS-T + 3% NGS for 24 h at 4°C. After washing 3x for 10 minutes in TBS-T the sections were incubated with goat anti-rabbit IgG (H+L) Alexa Fluor™ 647 at 1:1000 (Invitrogen™, A21244) for 24 h at 4°C in the dark. The sections were washed 3x for 10 minutes in TBS-T and stained with primary anti-rabbit-NeuN-488 (Abcam, ab190195) for 24h at 4°C in the dark. The liquid on the sections was briefly dried off and they mounted with Fluoromount-G Mounting Medium with DAPI (Invitrogen, 00-4959-52). The sections were incubated for 24 h at 4°C before being imaged using confocal microscopy (FV 1200, Olympus Fluoroview) using a 30x silicon objective (NA1.05). Images were pre-processed by ImageJ^51,52^ and quantified using CellProfiler (v.4.2.5.,^53^). Statistics were calculated in GraphPad Prism.

#### Mouse NPC isolation

Mouse NPCs were isolated from the cortex of E17.5 C57BL6/NTac embryos. The protocol was adapted from Bernas et al, 2017^54^. Embryo cortices were manually dissected from brains and the tissue was dissociated in 1X TrypLE™ Select Enzyme (Thermo Fisher Scientific, A1217701) for 10 minutes, with manual dissociation using a 1000 µL pipette. The cell suspension was washed in Neurobasal medium (Thermo Fisher Scientific, 21103049) and filtered through a 70 µm cell strainer (Miltenyi Biotech, 130-110-916). Cells were washed twice in 5 mL Neurobasal medium (Gibco, 21103049) by centrifugation at 200 x g for 10 minutes. Cells were resuspended in 1 mL Neurobasal growth medium containing 1X B27 supplement (Thermo Fisher Scientific, 17504044), 1X Penicillin/Streptomycin (Thermo Fisher Scientific, 15140122), 1X Glutamax (Thermo Fisher Scientific, 35050038), 20ng/mL FGF-2 (Peprotech, 100-18B), 20 ng/mL EFG (Peprotech, AF-100-15), and 2 µg/mL Heparin (MP Biomedicals, 210193125). Cells were seeded onto 12-well plates, previously coated for 2 h at 37°C in 1:100 dilution of Matrigel (Corning, 354234). After passaging, NPCs were cultured in the media described above on plates coated overnight in poly-D-lysine (Sigma, P7280), and then coated for 2 h in laminin (Sigma, L2020). Cells were either sub-cultured or media was changed every 2 days.

#### RNAseq on mNPC

Primary mouse NPC lines from 4 separate embryos were seeded 0.3 × 10^6^ cells per well, in two batches, in a poly-D-lysine/laminin coated 6-well plate after 9 days *in vitro*. Following 36 h of incubation at 37°C, one plate with 4 lines was incubated at 32°C for 6 h. Cells from both 37°C and 32°C conditions were harvested in tri-reagent and RNA was isolated using the Direct-zol RNA Microprep kit (Zymo Research, R2063). RNAseq library construction and Illumina next generation paired-end sequencing was performed by Novogene (NovaSeq PE150), generating 30 million paired-end reads per sample. For the RNAseq analysis, Fastq files were pseudo-aligned to the GRCm39 mouse transcriptome, downloaded from NCBI, using Kallisto (v.0.48.0,^55^) with paired-end mode on and 100 bootstraps. Kallisto output files were imported into R using the tximport package. Transcripts were assigned to genes using the TxDb.Mmusculus.UCSC.mm10.ensGene package (v.3.4.0) in R. Differential expression analysis and Z-scoring was performed using the DESeq2 package (v.3.16,^56^) using variance stabilization transformation for normalization, in R, after discarding transcripts with less than 10 counts. Differentially expressed genes were defined as those with an adjusted p-value less than 0.1. Overlaps of gene lists were calculated using the GeneOverlap package (v.1.34.0,^57^) in R, with overlaps tested using Fisher’s exact test.

#### CUT&RUN

HEK293WT and human NPCs (ATCC, ACS-1019) were harvested after 6 h at 32°C or kept at 37°C for the CUT&RUN. For making the *SMYD5*-KD we transfected the HEK293 cells with predesigned DsiRNA (IDT, hs.Ri.SMYD5.13.8) and Negative control using Lipofectamine 3000 (Thermo Fisher Scientific, L3000015) based on the manufactureŕs protocol. After 48 hours of transfection, the cells were placed in 32°C and 37°C for 6 hours post which they were harvested for CUT&RUN and RT-qPCR (for validation of the knockdown of *SMYD5*). CUT&RUN was performed according to Epicypher CUTANA protocol (v.1.7) on 300.000 cells per sample. We permeabilized the cells with 0.01% digitonin (Sigma, D141). For normalization we spiked in E. Coli DNA (Epicypher, 23618-1401) at the final concentration of 0.2 ng per sample. The library preparation was performed with TrueSeq-Chip Sample preparation Kit (Illumina, IP-202-9001). The following antibodies were used for the CUT&RUN: H3K36me3 (Thermo Fisher Scientific, MA5-24687), H4K20me3 (EpiCypher, 13-0054), H3K4me3 (EpiCypher, 13-0041 or Abcam, ab8580), H3K27me3 (Thermo Fisher Scientific, MA5-11198), H3K9me3 (Abcam, ab176916), rabbit anti-SMYD5 ^36^, and Rabbit IgG Antibody (Epicypher, 23613-0042). The sequencing was performed on a NovaSeq 6000 S4 at deCODE genetics with 150 bp paired-end sequencing, where we aimed for 10 million reads per sample. For our final batch, we sequenced on a NextSeq 550 using the NextSeq 500/550 Mid Output Kit v2.5 (20024904) with 75 bp paired-end sequencing. For the CUT&TAG re-analysis we followed published STAR protocols^58^ and used SEACR for peak calling^59^. For the CUT&RUN, briefly, we demultiplexed the files and then we used Bowtie2 (v.2.4.4,^60^) to align the reads to Hg38 and E. Coli reference genome, all downloaded from UCSC. Overall alignment to Hg38 were from 83.6-99.13% and to E. Coli spike in were from 0-4.18% for our data. The aligned Sam files were then converted to Bam files, indexed, and sorted via SAMtools (v.1.15,^61^). We used deepTools (v. 3.5.1,^62^) with RPGC normalization for making bigwig files for visualization in the UCSC Genome Browser as well as visualizing the Plot Profile of distinct modifications. We used MACS3 (v.3.0.0,^63^) for peak calling for the CUT&RUN, we used the broad mode for H3K4me3 and H3K36me3 with broad-cutoff set to 0.05 and --keep-dup all, we used --max-gap 300 for the H3K4me3 modification and --max-gap 20 for H3K36me3 modification. We called individual bam files for each replicate against merged IgG files for each condition. We used DiffBind (v.3.9.6,^64^) for the differential peak analysis, where we used the default normalization and analyzing mode. ChIPseeker (v.1.35.3, ^65^), TxDb.Hsapiens.UCSC.hg38.knownGene (v.3.4.6) and org.Hs.eg.db (v.3.8.2) were used for annotation, all in R^50^. For the comparison of *SMYD5*-KD at 37°C versus *SMYD5*-WT at 37°C and *SMYD5*-WT at 37°C versus 32°C we first excluded all genes that had p≥0.05 after DiffBind analysis. Next, we only kept genes that had sites that had a Log_2_FC ˂-1 for the *SMYD5*-WT at 37°C versus 32°C dataset. We then calculated the mean Log_2_FC of the promoter or distal intergenic sites for each gene for the two datasets and depicted those values. Genes that were both upregulated and downregulated in some of our datasets were excluded from the comparison.

### Quantification and statistical analysis

Statistical details can be found in the figure legends, and method details section of STAR methods. In this study we often calculated significance levels in GraphPad Prism, * = P<0.05, ** = P<0.01, *** = P<0.001, **** = P<0.0001. -Log10(RRA) score and LFC was calculated with MAGeCK^29^. Statistical analysis for Venn graphs were done with GeneOverlap^57^ package in R using Fisher’s Exact test. Significance levels from DESeq2^56^ analysis, were considered significant if p. adjusted < 0.1, calculated with DESeq2 in R. Significance levels calculated using DiffBind^64^ were considered significant if p value < 0.05 and/or Log2FC < −1.

### Additional resources

**Excel table: Table S2. Top candidate genes found enriched in CRISPR-Cas9 knock out mutation screening, related to Fig. 2**.

