## supplemental file for "SMYD5 is a regulator of the mild hypothermia response"

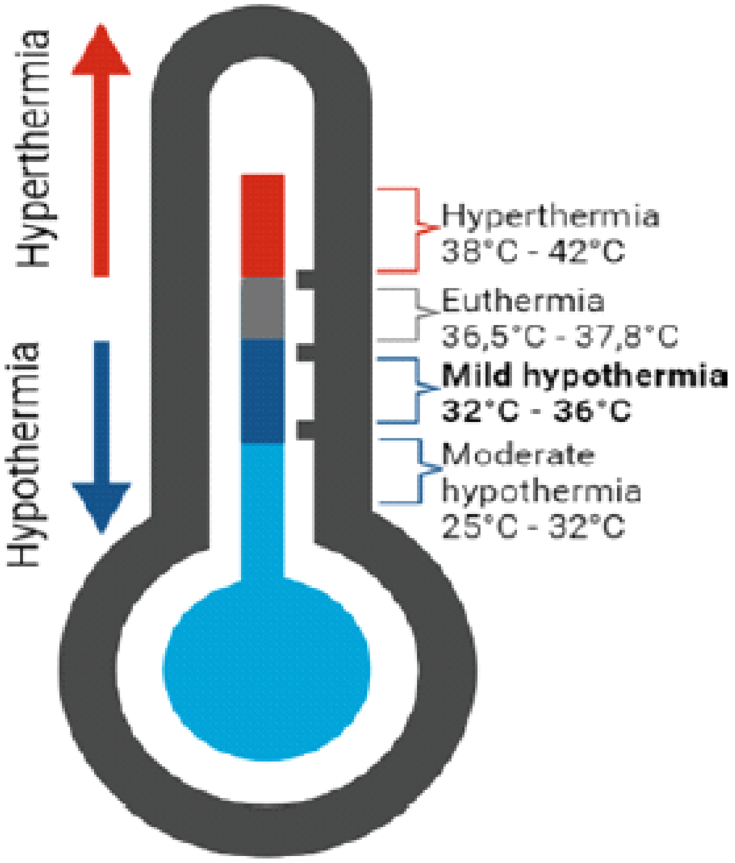


**Fig. S1. Different temperature ranges in humans, related to Fig. 1.** Temperatures known to induce distinct responses (moderate and mild hypothermia, euthermia and hyperthermia). Our range of interest (mild hypothermia) is bolded.

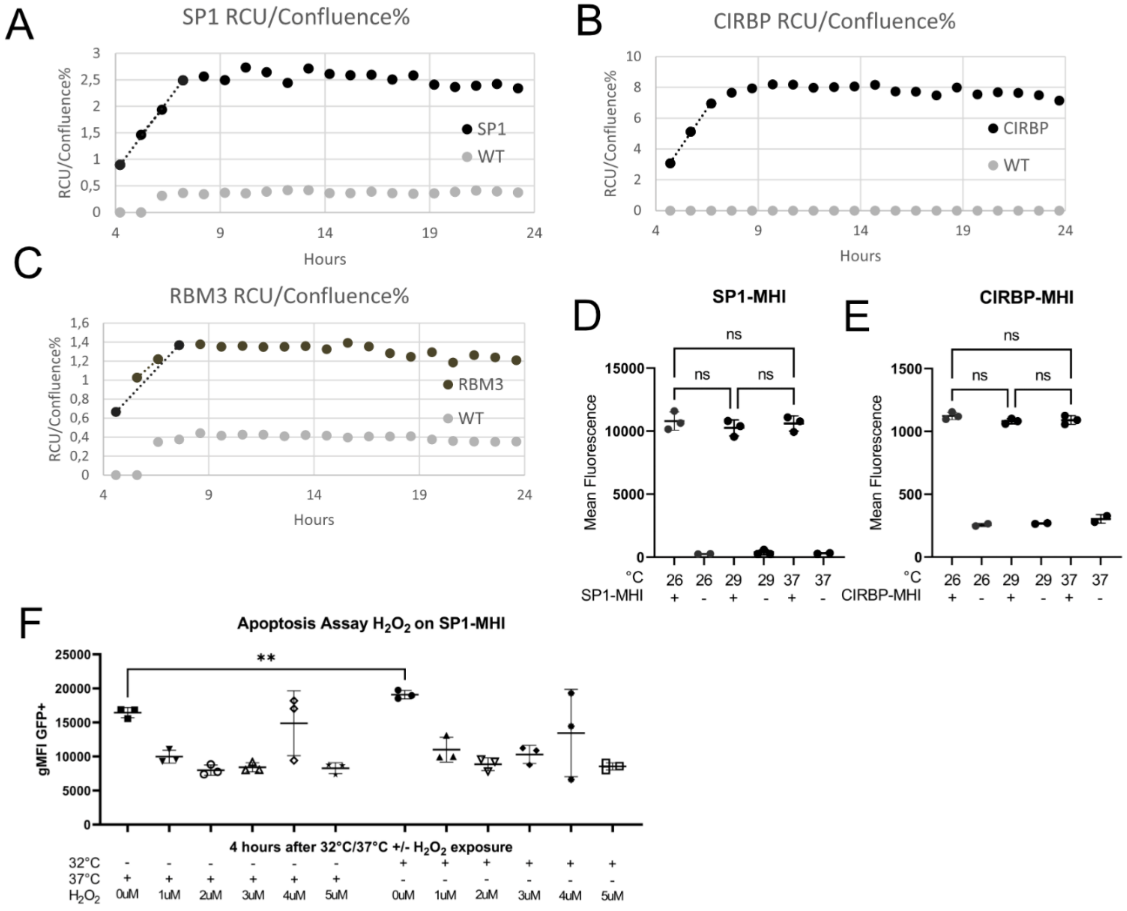


**Fig. S2. MHI activation peaks during mild hypothermia (32°C) and is limited at other temperatures, related to Fig. 1.** (A-C) Measured fluorescence hourly using an IncucyteS3 at 32°C using SP1-Lenti-MHI, CIRBP-Lenti-MHI, RBM3-Lenti-MHI respectively, in HEK293 cells (n=1). (D-E) No change in fluorescence from euthermia was observed at moderate hypothermia (26°C and 29°C) for either SP1-MHI or CIRBP-MHI in HEK293 cells. Each point is a technical replicate (n=3), with mean and SD where applicable. Significance levels were calculated with Šidák's multiple comparison test. (F) Apoptosis induction analysis done with $H_2O_2$ exposure on HEK293+Cas9+SP1 cell line. Shown are data points, each point a technical replicate (n=3), mean and SD where applicable. Significance levels calculated with an unpaired two-tailed t-test calculated in GraphPad Prism. All significance levels in this figure were calculated in GraphPad Prism, * = P<0.05, ** = P<0.01, *** = P<0.001, **** = P<0.0001.

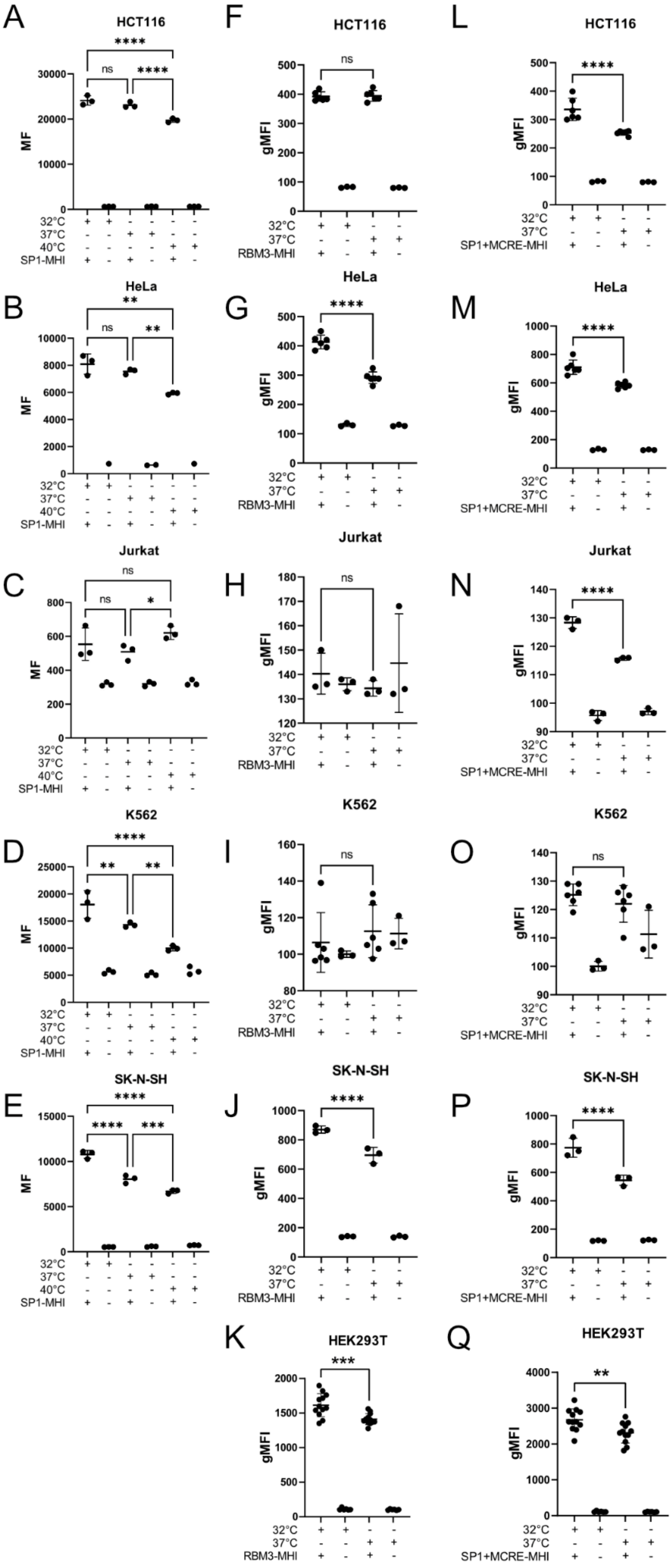

A
HCT116
MF
32°C
37°C
40°C
SP1-MHI
B
HeLa
C
Jurkat
D
K562
E
SK-N-SH
F
gMFI
RBM3-MHI
G
H
I
J
K
HEK293T
L
SP1+MCRE-MHI
M
N
O
P
Q
ns

**Fig. S3. MHI demonstrates consistent activation at 32°C in diverse cellular systems, related to Fig. 1.** (A-E) Left column shows mean fluorescence (MF, flow cytometry) after transient transfection of SP1-MHI into HCT116, HeLa, Jurkat, K562 and SK-N-SH (16 h at 32°C, 37°C or 40°C). Shown are data points, each data point is a technical replicate (n=3), mean and SD where applicable, Šidák‘s multiple comparison test calculated in GraphPad Prism. (F-K) Middle column shows gMFI (flow cytometry) after transient transfection of RBM3-MHI into HCT116, HeLa, Jurkat, K562, SK-N-SH, and HEK293T, 16 h at 32°C or 37°C. Shown are data points, each data point is a technical replicate (n=3-11), mean and SD where applicable, Tukey‘s multiple comparison test calculated in GraphPad Prism. (L-Q) Right column shows gMFI (flow cytometry) after transient transfection of SP1+MCRE-MHI into HCT116, HeLa, Jurkat, K562, SK-N-SH, and HEK293T, 16 h at 32°C or 37°C, (n=3-11). Data shown as in (F-K). * = P<0.05, ** = P<0.01, *** = P<0.001, **** = P<0.0001.

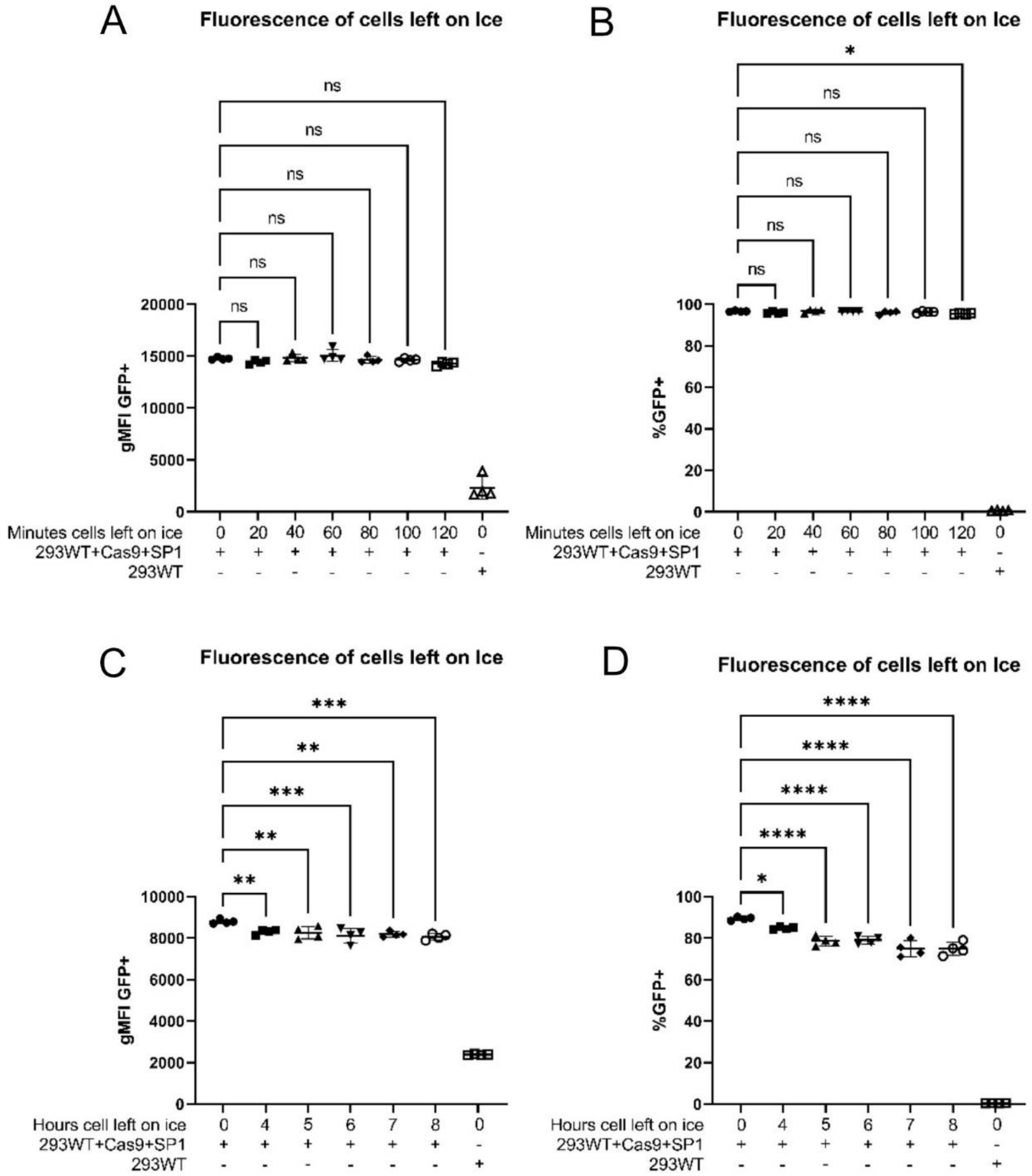


**Fig. S4. Storage of samples on ice does not lead to activation of MHI, related to Fig. 2.** (A-D) gMFI or percentage of fluorescent cells of HEK293WT+Cas9+SP1 left on ice for different periods of time before flow cytometry analysis. Shown are data points, each data point is a technical replicate, mean and SD where applicable. Dunnett's multiple comparison test calculated in GraphPad Prism. * = P<0.05, ** = P<0.01, *** = P<0.001, **** = P<0.0001.

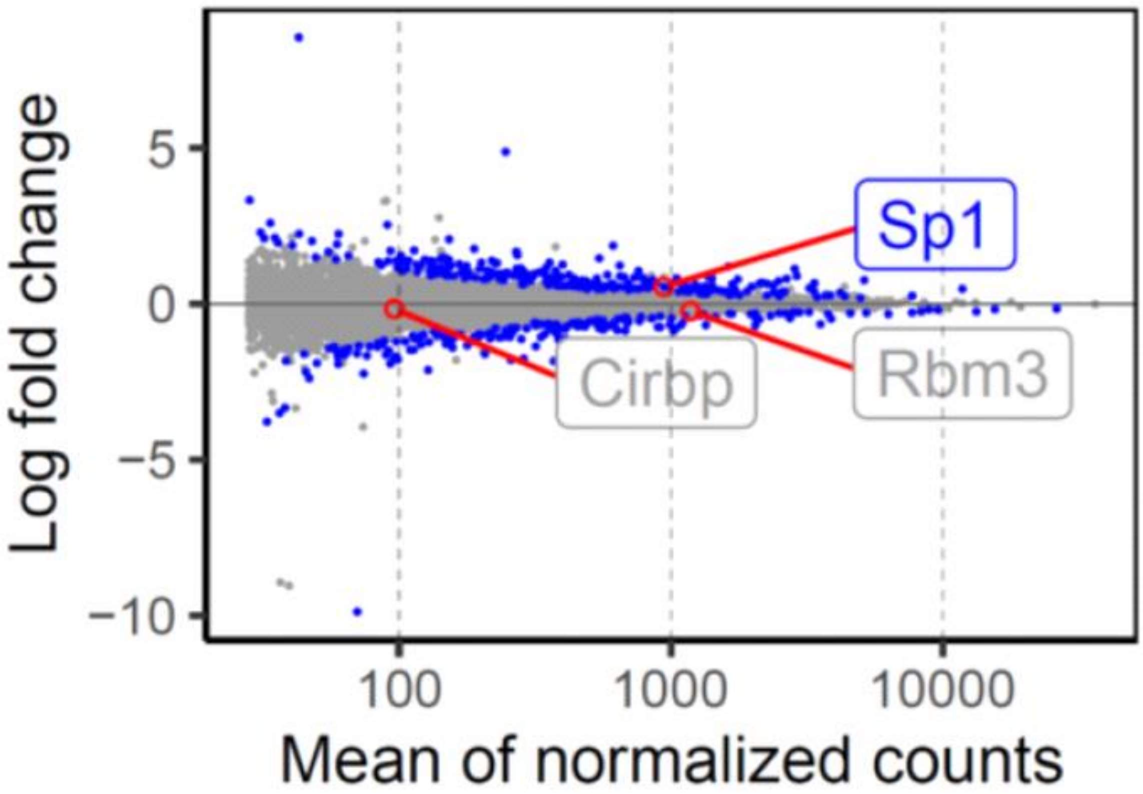


**Fig. S5. *Cirbp* has fewer normalized counts compared to *Sp1* and *Rbm3,* related to Fig. 3.** Normalized counts from RNAseq in mESc (referring to data from **Fig. 3**).

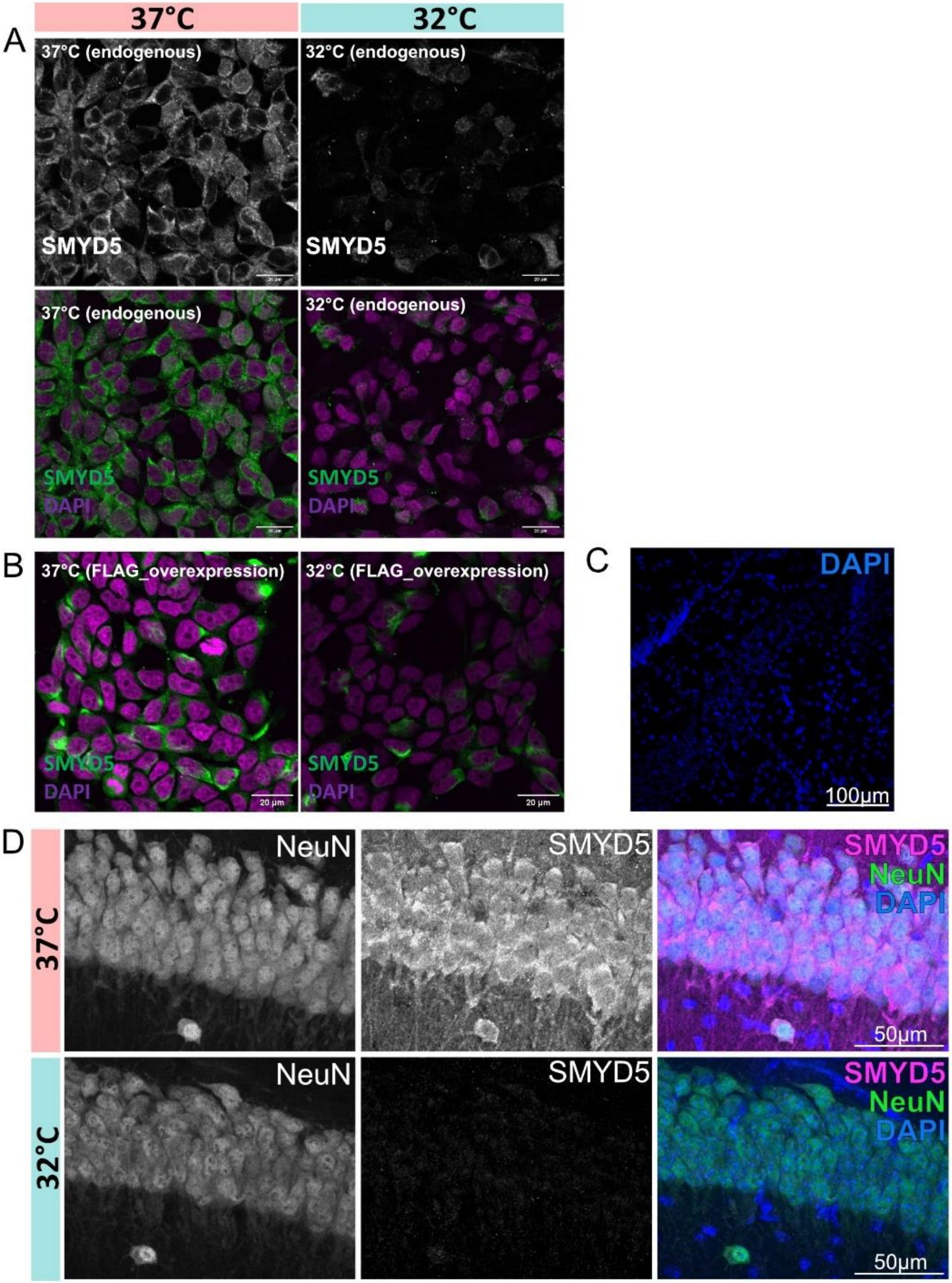

37°C
32°C
A
37°C (endogenous)
SMYD5
32°C (endogenous)
SMYD5
37°C (endogenous)
SMYD5
DAPI
32°C (endogenous)
SMYD5
DAPI
B
37°C (FLAG_overexpression)
SMYD5
DAPI
20 µm
32°C (FLAG_overexpression)
SMYD5
DAPI
20 µm
C
DAPI
100µm
D
37°C
NeuN
SMYD5
SMYD5
NeuN
DAPI
50µm
32°C
NeuN
SMYD5
SMYD5
NeuN
DAPI
50µm

**Fig. S6. Validation of temperature dependence of SMYD5, related to Fig. 4.** (A) Another independent biological replicate with anti-SMYD5 antibody at 32°C and 37°C. Green or white (SMYD5), purple (DAPI). (B) A validation with anti-FLAG antibody, showing the same difference in SMYD5 amount as in (A) at the two temperatures. Green (SMYD5), purple (DAPI). (C) An image showing secondary antibody only staining using anti-rabbit alexa-647 and DAPI. (D) SMYD5, DAPI and NeuN staining in brain sections of mice kept at euthermia (37°C; upper) after neonatal hypoxic-ischemic injury compared to those treated with cooling at 32°C, for 6 h (lower). Sagittal brain sections are from the CA1 region of the hippocampus.

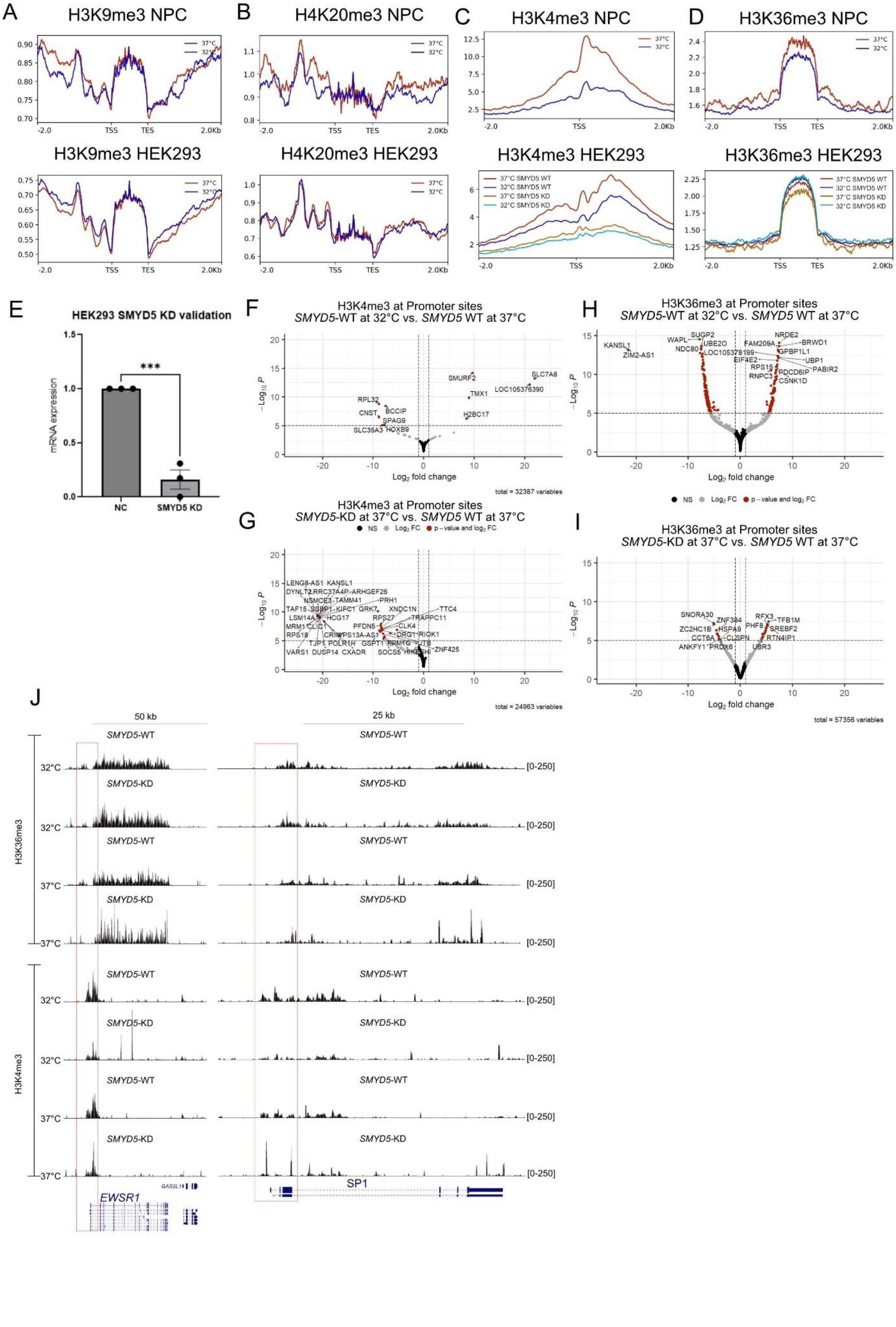

A
H3K9me3 NPC
H3K9me3 HEK293
B
H4K20me3 NPC
H4K20me3 HEK293
C
H3K4me3 NPC
H3K4me3 HEK293
D
H3K36me3 NPC
H3K36me3 HEK293
37°C
32°C
37°C SMYD5 WT
32°C SMYD5 WT
37°C SMYD5 KD
32°C SMYD5 KD
TSS
TES
-2.0
2.0Kb
E
HEK293 SMYD5 KD validation
mRNA expression
***
NC
SMYD5 KD
F
H3K4me3 at Promoter sites
SMYD5-WT at 32°C vs. SMYD5 WT at 37°C
G
H3K4me3 at Promoter sites
SMYD5-KD at 37°C vs. SMYD5 WT at 37°C
H
H3K36me3 at Promoter sites
SMYD5-WT at 32°C vs. SMYD5 WT at 37°C
I
H3K36me3 at Promoter sites
SMYD5-KD at 37°C vs. SMYD5 WT at 37°C
$-Log_{10}$ P
$Log_2$ fold change
NS
$Log_2$ FC
p - value and $log_2$ FC
total = 32387 variables
total = 24963 variables
total = 57356 variables
J
50 kb
25 kb
H3K36me3
H3K4me3
SMYD5-WT
SMYD5-KD
32°C
37°C
[0-250]
EWSR1
GAS2L1
SP1

**Fig. S7. SMYD5 is a temperature dependent repressor via H3K36me3 at 37°C, related to Fig. 5.** (A) There is not a global change in H3K9me3 at 32°C compared to 37°C in either NPC (top, n=3) nor HEK293 (bottom, n=3) cell lines. (**B**) There is not a global change in H4K20me3 at 32°C compared to 37°C in either NPC (top, n=3) nor HEK293 (bottom, n=3) cell lines. (**C**) We observe a decrease in H3K4me3 at 32°C compared to 37° in both NPC (top, n=3) and HEK293 (bottom, n=5) cell lines. We see a further decrease in H3K4me3 with *SMYD5*-KD in HEK293 cell lines (n=3). (**D**) We observe a decrease in H3K36me3 at 32°C compared to 37° in NPC (top, n=3) cell line and the opposite in HEK293 (bottom, n=6) cell line. We see a further decrease in H3K36me3 at 37°C with *SMYD5*-KD (n=3) but we do not observe a change in H3K36me3 at 32°C with *SMYD5*-KD (n=3). (**E**) Validation of *SMYD5*-KD in HEK293 cells via *SMYD5* relative expression compared to *GAPDH* quantified with RT-qPCR. Significance level calculated with an unpaired two-tailed t-test in GraphPad Prism. (**F**) Volcano plot depicting Log2 Fold Change of called H3K4me3 modification peaks at 32°C (n=5) compared to 37°C (n=5) in SMYD5-WT HEK293 cell line. Each data point is a called peak. (**G**) Volcano plot depicting $Log_2$ Fold Change of called H3K4me3 modification peaks at promoter site at 37°C SMYD5-KD (n=3) compared to 37°C (n=5) in SMYD5-WT HEK293 cell line. Data shown as in (F). (**H**) Volcano plot depicting Log2 Fold Change of called H3K36me3 modification peaks at 32°C (n=6) compared to 37°C (n=6) in SMYD5-WT HEK293 cell line. Data shown as in (F). (**I**) Volcano plot depicting $Log_2$ Fold Change of called H3K36me3 modification peaks at promoter site at 37°C SMYD5-KD (n=3) compared to 37°C (n=6) in SMYD5-WT HEK293 cell line. Data shown as in (F). (**J**) Two examples, *EWSR1* and *SP1*, of modification peaks for H3K36me3 and H3K4me3. Red box surrounds the promoter region of the genes. * = P<0.05, ** = P<0.01, *** = P<0.001, **** = P<0.0001.

**Table S1. Specifics of our MHI, including sequence material used, vector, selection markers, plasmid size and individual fluorophores, related to Fig. 1.**

| | **Promoter sequence, location in bp from TSS** | **Animal** | **Location** | **Repeats of MCRE** | **Plasmid size without MCRE** | **Plasmid size with MCRE** | **Selection marker** | **Fluorescent protein** | **Backbone plasmid** |
|---|---|---|---|---|---|---|---|---|---|
| **CIRP Indicator** | -452 to +72 | Mouse | Chr. 10: 80167244-80168059 | 5 | 5571 | 5657 | Neomycin | AcGFP | pGL4.10 |
| **RBM3 Indicator** | -667 to +229 | Human | Chr. X: 48573772-48574678 | 6 | NA | 7557 | Neomycin | TurboGFP | pCMV6-AC-GFP |
| **SP1 Indicator** | -281 to -20 | Human | Chr. 12: 53379972-53380272 | 5 | 5313 | 5392 | Neomycin | AcGFP | pGL4.10 |
| **CIRP lentiviral Indicator** | -448 to +153 | Human | Chr. 19: 1268820-1269420 | 8 | NA | 8959 | Neomycin | mCherry | pLV-Ex |
| **RBM3 lentiviral Indicator** | -667 to +229 | Human | Chr. X: 48573772-48574678 | 5 | NA | 9235 | Neomycin | mCherry | pLV-Ex |
| **SP1 lentiviral Indicator** | -358 to -96 | Human | Chr. 12: 53379895-53380156 | 5 | NA | 8590 | Neomycin | mCherry | pLV-Ex |

**Table S2. Top candidate genes found enriched in CRISPR-Cas9 knock out mutation screening, related to Fig. 2.**

**SP1 candidate repressors**

| **Gene Symbol** | **Score** | **FDR** | **GeCKO_number** |
|---|---|---|---|
| *STXBP1* | 5.704806884 | 0.034653 | 1 |
| *LRRC26* | 5.193650413 | 0.081683 | 3 |
| *MUC15* | 5.009257235 | 0.081683 | 4 |
| *GATAD2A* | 4.994647864 | 0.081683 | 5 |
| *MUT* | 4.717423199 | 0.082744 | 7 |
| *SLC18A3* | 4.627216832 | 0.10396 | 8 |
| *SCGB3A2* | 4.564474149 | 0.10396 | 9 |
| *hsa-mir-4314* | 4.555549219 | 0.10396 | 10 |
| *GFOD2* | 4.525419548 | 0.10396 | 11 |
| *LILRB5* | 4.49629101 | 0.10396 | 12 |
| *CCDC73* | 4.452816239 | 0.10396 | 13 |
| *PTPN11* | 4.434104978 | 0.10396 | 15 |
| *hsa-mir-6832* | 4.364385582 | 0.116773 | 16 |
| *FAM169B* | 4.340359479 | 0.116773 | 17 |
| *hsa-let-7i* | 4.32670256 | 0.117633 | 18 |
| *GPR18* | 4.31068226 | 0.117633 | 19 |
| *ATXN10* | 4.250936512 | 0.117633 | 21 |
| *ZNF813* | 4.246057093 | 0.119018 | 22 |
| *KAZALD1* | 4.197000637 | 0.119018 | 23 |
| *RAB11FIP5* | 4.192877815 | 0.119018 | 24 |
| *KAT5* | 4.156898091 | 0.127327 | 25 |
| *ZNF260* | 4.128327842 | 0.131096 | 26 |
| *CGB1* | 4.122202632 | 0.133663 | 27 |
| *HRC* | 4.091718772 | 0.131096 | 28 |
| *hsa-mir-137* | 4.066325469 | 0.133663 | 29 |
| *hsa-mir-578* | 4.044788648 | 0.133663 | 30 |
| *DDA1* | 4.043658655 | 0.133663 | 31 |
| *hsa-mir-548a-1* | 4.011548069 | 0.133663 | 32 |
| *ANKZF1* | 4.011222613 | 0.133663 | 33 |
| *UVRAG* | 3.996281036 | 0.133663 | 34 |
| *PTGS2* | 3.975391204 | 0.133663 | 35 |
| *hsa-mir-558* | 3.944621669 | 0.15186 | 36 |
| *ZDHHC21* | 3.932111193 | 0.142052 | 37 |
| *MNS1* | 3.907700523 | 0.15568 | 38 |
| *hsa-mir-588* | 3.835379511 | 0.168097 | 39 |
| *CCDC59* | 3.830795963 | 0.167955 | 41 |
| *ZFHX2* | 3.830237423 | 0.168097 | 42 |
| *EIF5B* | 3.801755414 | 0.168097 | 43 |
| *DUS4L* | 3.790860464 | 0.167955 | 44 |

| | | | |
|---|---|---|---|
| *ULBP1* | 3.784415306 | 0.167955 | 45 |
| *LACTB* | 3.777647761 | 0.173869 | 46 |
| *ZNF829* | 3.732922183 | 0.173869 | 47 |
| *hsa-mir-3667* | 3.724251116 | 0.173869 | 48 |
| *hsa-mir-548l* | 3.71942163 | 0.173869 | 49 |
| *USP28* | 3.719285081 | 0.173869 | 50 |
| *KISS1R* | 3.71701556 | 0.173869 | 51 |
| *SGCB* | 3.716653535 | 0.173869 | 52 |
| *SMYD5* | 3.709609191 | 0.173869 | 53 |
| *BMPR1A* | 3.698514234 | 0.173869 | 54 |
| *CXorf58* | 3.681769661 | 0.173869 | 55 |
| *RNF175* | 3.67997841 | 0.173869 | 56 |
| *DMRT1* | 3.668917036 | 0.173869 | 57 |
| *DMRTA2* | 3.667622144 | 0.173869 | 58 |
| *KRT77* | 3.656926366 | 0.173869 | 59 |
| *MROH9* | 3.639690656 | 0.173869 | 60 |
| *TRMT44* | 3.637555226 | 0.173869 | 61 |
| *TYR* | 3.634212605 | 0.173869 | 62 |
| *KIAA1199* | 3.631434215 | 0.173869 | 63 |
| *hsa-mir-507* | 3.628821268 | 0.173869 | 64 |
| *PAX3* | 3.625930196 | 0.173869 | 65 |
| *KRT86* | 3.624482699 | 0.173869 | 66 |
| *RAD54L2* | 3.620277615 | 0.173869 | 67 |
| *FBXW9* | 3.611898798 | 0.173869 | 68 |
| *SERPINB4* | 3.609329787 | 0.173869 | 69 |
| *hsa-mir-6877* | 3.606354613 | 0.173869 | 70 |
| *STRA13* | 3.606126596 | 0.173869 | 71 |
| *OC90* | 3.605495787 | 0.173869 | 72 |
| *PRY* | 3.600603208 | 0.173869 | 73 |
| *ZNF579* | 3.592643093 | 0.173869 | 74 |
| *TTC37* | 3.591065041 | 0.174323 | 75 |
| *SAR1B* | 3.584375698 | 0.178034 | 76 |
| *RIF1* | 3.581682133 | 0.175026 | 77 |
| *SFTPA2* | 3.567030709 | 0.178034 | 78 |
| *SLC22A9* | 3.54945838 | 0.178025 | 79 |
| *RSPRY1* | 3.549442991 | 0.178755 | 80 |
| *PDK1* | 3.548781741 | 0.178034 | 81 |
| *CTDSPL* | 3.532939732 | 0.178755 | 82 |
| *MMP17* | 3.531948209 | 0.178034 | 83 |

**SP1 candidate activators**

| **Gene symbol** | **Score** | **FDR** | **GeCKO_number** |
|---|---|---|---|
| *PDS5B* | 4.567190995 | 0.183168 | 4 |

| | | | |
|---|---|---|---|
| *SYT1* | 4.508008336 | 0.183168 | 5 |
| *DDX18* | 4.454779504 | 0.183168 | 7 |
| *POU4F3* | 4.442324406 | 0.183168 | 8 |
| *ZNF71* | 4.430567564 | 0.183168 | 9 |
| *hsa-mir-6739* | 4.419177626 | 0.183168 | 10 |
| *PDE1B* | 4.334400249 | 0.183168 | 12 |
| *RHOT2* | 4.322758154 | 0.183168 | 14 |
| *CACNB1* | 4.315100804 | 0.183168 | 15 |
| *TXLNB* | 4.294777944 | 0.183168 | 17 |
| *GRB14* | 4.258147475 | 0.183168 | 19 |
| *AIFM1* | 4.249545606 | 0.183168 | 20 |
| *ARHGAP29* | 4.209911967 | 0.183168 | 21 |
| *hsa-mir-596* | 4.124921365 | 0.191368 | 23 |
| *hsa-mir-1323* | 4.093078034 | 0.191368 | 25 |
| *OR2M5* | 4.090080428 | 0.191368 | 26 |
| *TBC1D22B* | 4.065491449 | 0.191368 | 27 |
| *TOR2A* | 4.063642109 | 0.191368 | 28 |
| *CRTAC1* | 4.052497659 | 0.19757 | 29 |
| *FAM32A* | 4.036316038 | 0.191368 | 31 |
| *hsa-mir-1910* | 4.034633646 | 0.191368 | 32 |
| *PRAMEF22* | 3.98021927 | 0.191368 | 33 |
| *KLK2* | 3.949349619 | 0.210917 | 34 |
| *PSMA5* | 3.948886083 | 0.210917 | 35 |
| *HNRNPUL1* | 3.947421475 | 0.210917 | 36 |
| *SNX22* | 3.916784138 | 0.210917 | 37 |
| *PRSS45* | 3.902638555 | 0.21545 | 39 |
| *SIGLEC8* | 3.880579137 | 0.21545 | 40 |
| *C3orf80* | 3.873251858 | 0.21545 | 41 |
| *TRIM36* | 3.838003475 | 0.21545 | 43 |
| *PRKCE* | 3.826784585 | 0.21545 | 45 |
| *TLL2* | 3.8012055 | 0.21545 | 48 |
| *LCK* | 3.784521064 | 0.218102 | 49 |
| *HNRNPDL* | 3.773864278 | 0.218102 | 50 |
| *PGAP1* | 3.769117001 | 0.218102 | 51 |
| *LEO1* | 3.760900818 | 0.218102 | 52 |
| *hsa-mir-5091* | 3.752788029 | 0.2157 | 53 |
| *SPTAN1* | 3.730393669 | 0.225915 | 54 |
| *hsa-mir-4505* | 3.697387715 | 0.225915 | 55 |
| *KIF13B* | 3.682249971 | 0.225915 | 57 |
| *AMBN* | 3.66756154 | 0.225915 | 59 |
| *hsa-mir-6823* | 3.66524476 | 0.225915 | 60 |
| *hsa-mir-3176* | 3.660508808 | 0.225915 | 61 |
| *FKBP8* | 3.651714612 | 0.225915 | 62 |

| | | | |
|---|---|---|---|
| *RNASE2* | 3.625086763 | 0.228211 | 63 |
| *TRIL* | 3.618578042 | 0.228211 | 64 |
| *TAF13* | 3.617892864 | 0.228211 | 65 |
| *DLG3* | 3.616939023 | 0.228211 | 66 |
| *PTAFR* | 3.611348283 | 0.228211 | 67 |
| *hsa-mir-3925* | 3.588767628 | 0.225915 | 69 |
| *SCAF11* | 3.58384231 | 0.228211 | 70 |
| *NLGN3* | 3.577278874 | 0.233188 | 71 |
| *MS4A3* | 3.573439944 | 0.233188 | 72 |
| *hsa-mir-548aq* | 3.567239092 | 0.228211 | 73 |
| *ARRDC3* | 3.564076042 | 0.233188 | 74 |
| *MICALCL* | 3.554753965 | 0.233188 | 75 |
| *THUMPD1* | 3.549427601 | 0.233188 | 76 |
| *NKX6-2* | 3.537212809 | 0.238057 | 77 |
| *HS6ST2* | 3.532480686 | 0.233188 | 78 |
| *BANF1* | 3.518815445 | 0.238057 | 80 |

**Table S3. List of repressed SMYD5-regulated genes at 37°C, related to Fig. 5**

| Genes |
|---|
| *Mybl2* |
| *Hspg2* |
| *Xrn2* |
| *Fabp3* |
| *Ywhab* |
| *Phf20* |
| *Neo1* |
| *Tulp4* |
| *Trip12* |
| *Nt5e* |
| *Thrap3* |
| *Neat1* |
| *Mier1* |
| *Rell1* |
| *Eif4g3* |
| *Ift52* |
| *Tfcp2l1* |
| *Fam21* |
| *Ctc1* |
| *Lamc1* |
| *Amotl2* |
| *Gabbr1* |
| *Hnrnpul1* |
| *Asxl1* |
| *Atp2b1* |
| *Atp2a3* |
| *Klf15* |
| *Cd24a* |
| *Tex15* |
| *Slc25a12* |
| *Slc35g1* |
| *Stag1* |
| *Pkn1* |
| *Vps41* |
| *Ewsr1* |
| *Acsf2* |
| *Ass1* |

**Table S4. Related to STAR Methods Key Resource Table (Oligonucleotides section): Full list of oligonucleotides used in this study, both primers and DsiRNA.**

| Oligonucleotide name | Source or Sequence |
|---|---|
| *CIRBP* mouse forward primer | 5'-tcgataggtaccTGGCTTCACAAATGCGCCTCAGT-3' |
| *CIRBP* mouse reverse primer | 3'-cctaaggcagatctGCGAGGGGGAGCGCAAGAGT-5' |
| *SP1* human forward primer | 5'-tcaagtcaggctagcGCAACTTAGTCTCACACGCCTTGG-3' |
| *SP1* human reverse primer | 3'-cagtgctgcctcgagGCTCAAGGGGGTCCTGTCCGG-5' |
| AcGFP1 forward primer | 5'-taagccaccATGGTGAGCAAGGGCGAGGAGC-3' |
| AcGFP1 reverse primer | 3'-cggcggagTCTAGAATTACTTGTACAGCTCGTCC-5' |
| Neomycin forward primer | 5'-CATTATCGTCGACTCTACCGGGTAGGGGAGGCGCTT-3' |
| Neomycin reverse primer | 3'-CGCCGCCGACGATAGTCAAGCTTCTGATGGAATTAGAACTTGGC-5' |
| SMYD5-2mut-F: | 5'-gctctttacgAGGAAGCAGTCAGCCAGT-'3 |
| SMYD5-2-mut-R: | 5'-ttccgtaaagAGTCTCCGCAGAAGTTCC-'3; |
| SMYD5-6-mut-F: | 5'-accgatatcgAGCCTGTGACCACTGCCT-'3; |
| SMYD5-6-mut-R: | 5'-atagagcgtt-CCAGAGAAACTGTGCAGCC-'3 |
| DsiRNA hs.Ri.SMYD5.13.8 | IDT |
| RT-qPCR SP1 (fwd) | 5'-CACCCAATTCAAGGCCTGCCGT-3' |
| RT-qPCR SP1 (rev) | 5'-GGGTTGGGCATCTGGGCTGTTT-3' |
| RT-qPCR RBM3 (fwd) | 5'-GAGACTCAGCGGTCCAGGGGTT-3' |
| RT-qPCR RBM3 (rev) | 5'-CCTCTGGTTCCCCGAGCAGACT-3' |
| RT-qPCR CIRBP (fwd) | 5'-CCGAGTTGACCAGGCTGGCAAG-3' |
| RT-qPCR CIRBP (rev) | 5'-TCCATAGCCCCGGTCTCCTCCT-3' |
| RT-qPCR GAPDH (fwd) | 5'-TCAAGGCTGAGAACGGGAAG-3' |
| RT-qPCR GAPDH (rev) | 5'-CGCCCCACTTGATTTTGGAG-3' |
| RT-qPCR SMYD5 1 (fwd) | 5'-GCACTGTGCGCAAAGACCTCCA-3' |
| RT-qPCR SMYD5 2 (fwd) | 5'-GGAAACCAGGCCAGGTTCTGCC-3' |
| RT-qPCR SMYD5 3 (fwd) | 5'-CGTGGAAGTCCGTTTCGTGA-3' |

| | |
|---|---|
| RT-qPCR SMYD5 1 (rev) | 5’-CTGGGCACAGGACCTGGTGGTA-3’ |
| RT-qPCR SMYD5 2 (rev) | 5’-GGCTGCCAACCGACATTCTGCA-3’ |
| RT-qPCR SMYD5 3 (rev) | 5’-CCAGAGAAACTGTGCAGCCA-3’. |

**File S1.**

Sequences of individual MHIs.

CIRBP-MHI (CIRP+GFP+Neo-pGL4.10)
*CIRBP* promoter
5′GGCCTAACTGGCCGGTACCCATCTGTCTTCTGGCTTCACAAATGCGCCTCAGTTAAATAACTGTAATT
AAATATTAAAAGAAAAAAAAGAACATAAAAGTGTTGAGTCGATGAGTCAAGGTTGGAGCCATATTTG
GGAGTCTCATTGAGGTAGCAAAAGCTCTTTGTTCCCCGCCGCAGGACAAACGGGTCGGGAACGCGGA
AGTGGGTCAGCTCCGCCTCCAAATCAGCTGTCCCCGCCTCCTAGTCGCTAGCGGGCGGGGGGTGGGGG
GGAAAAATGCGGCCAGGAAAGATTCCAGCCAATCAAAACACTCAGCGCCTCCGCGCGGACTAGGGCA
AAGCCAACGGAGTCCCGCCCCTCCACCGAGGCGCAGATGCCGAATCTAAGGCGTCGGATTGGTCAGC
GCGGCGTGGTGGGCGGGTTAAGGGGCGTGGCTCCAGGAGTGGAGTATATCAGGCGGGACTCTTGCGC
TCCCCCTCGCACTCGCGCCTTAGGAAGCTTGGGTGTGTGTGGCGCGCTGTCTTCCCGCTCGCGTCAGGG
ACCTGCCCGACTCAGCGGTGAGGGAGATCTGGCCTCGGCGGCCAAGCTTGGCAATCCGGTACTGTTGG
TAAAGCCACCATGGATGGTGAGCAAGGGCGCCGAGCTGTTCACCGGCATCGTGCCCATCCTGATCGAG
CTGAATGGCGATGTGAATGGCCACAAGTTCAGCGTGAGCGGCGAGGGCGAGGGCGATGCCACCTACG
GCAAGCTGACCCTGAAGTTCATCTGCACCACCGGCAAGCTGCCTGTGCCCTGGCCCACCCTGGTGACC
ACCCTGAGCTACGGCGTGCAGTGCTTCTCACGCTACCCCGATCACATGAAGCAGCACGACTTCTTCAA
GAGCGCCATGCCTGAGGGCTACATCCAGGAGCGCACCATCTTCTTCGAGGATGACGGCAACTACAAGT
CGCGCGCCGAGGTGAAGTTCGAGGGCGATACCCTGGTGAATCGCATCGAGCTGACCGGCACCGATTTC
AAGGAGGATGGCAACATCCTGGGCAATAAGATGGAGTACAACTACAACGCCCACAATGTGTACATCA
TGACCGACAAGGCCAAGAATGGCATCAAGGTGAACTTCAAGATCCGCCACAACATCGAGGATGGCAG
CGTGCAGCTGGCCGACCACTACCAGCAGAATACCCCCATCGGCGATGGCCCTGTGCTGCTGCCCGATA
ACCACTACCTGTCCACCCAGAGCGCCCTGTCCAAGGACCCCAACGAGAAGCGCGATCACATGATCTAC
TTCGGCTTCGTGACCGCCGCCGCCATCACCCACGGCATGGATGAGCTGTACAAGTGATCTAGAGTCGG
GGCGGCCGGCCGCTTCGAGCAGACATGATAAGATACATTGATGAGTTTGGACAAACCACAACTAGAA
TGCAGTGAAAAAAATGCTTTATTTGTGAAATTTGTGATGCTATTGCTTTATTTGTAACCATTATAAGCT
GCAATAAACAAGTTAACAACAACAATTGCATTCATTTTATGTTTCAGGTTCAGGGGGAGGTGTGGGAG
GTTTTTTAAAGCAAGTAAAACCTCTACAAATGTGGTAAAATCGATAAGGATCCGTCGACtctaccgggtaggg
gaggcgcttttcccaaggcagtctggagcatgcgctttagcagccccgctgggcacttggcgctacacaagtggcctctggcctcgcacacattccacatccaccggta
ggcgccaaccggctccgttctttggtggccccttcgcgccaccttctactcctcccctagtcaggaagttcccccccgccccgcagctcgcgtcgtgcaggacgtgacaa
atggaagtagcacgtctcactagtctcgtgcagatggacagcaccgctgagcaatggaagcgggtaggcctttggggcagcggccaatagcagctttgctccttcgcttt
ctgggctcagaggctgggaaggggtgggtccgggggcgggctcaggggcgggctcaggggcggggcgggcgcccgaaggtcctccggaggcccggcattctgc
acgcttcaaaagcgcacgtctgccgcgctgttctcctcttcctcatctccgggcctttcgacctgcagccaatatgggatcggccattgaacaagatggattgcacgcaggt
tctccggccgcttgggtggagaggctattcggctatgactgggcacaacagacaatcggctgctctgatgccgccgtgttccggctgtcagcgcaggggcgcccggttc
tttttgtcaagaccgacctgtccggtgccctgaatgaactgcaggacgaggcagcgcggctatcgtggctggccacgacgggcgttccttgcgcagctgtgctcgacgtt
gtcactgaagcgggaagggactggctgctattgggcgaagtgccggggcaggatctcctgtcatctcaccttgctcctgccgagaaagtatccatcatggctgatgcaat
gcggcggctgcatacgcttgatccggctacctgcccattcgaccaccaagcgaaacatcgcatcgagcgagcacgtactcggatggaagccggtcttgtcgatcaggat
gatctggacgaagagcatcaggggctcgcgccagccgaactgttcgccaggctcaaggcgcgcatgcccgacggcgatgatctcgtcgtgacccatggcgatgcctg
cttgccgaatatcatggtggaaaatggccgcttttctggattcatcgactgtggccggctgggtgtggcggaccgctatcaggacatagcgttggctacccgtgatattgct
gaagagcttggcggcgaatgggctgaccgcttcctcgtgctttacggtatcgccgctcccgattcgcagcgcatcgccttctatcgccttcttgacgagttcttctgagggg
atccgctgtaagtctgcagaaattgatgatctattaaacaataaagatgtccactaaaatggaagtttttcctgtcatactttgttaagaagggtgagaacagagtacctacattt
tgaatggaaggattggagctacgggggtgggggtggggtgggattagataaatgcctgctctttactgaaggctctttactattgctttatgataatgtttcatagttggatatc
ataatttaaacaagcaaaaccaaattaagggccagctcattcctcccactcatgatctatagatctatagatctctcgtgggatcattgtttttctcttgattcccactttgtggttct
aagtactgtggtttccaaatgtgtcagtttcatagcctgaagaacgagatcagcagcctctgttccacatacacttcattctcagtattgttttgccaagttctaattccatcagaa
gcttGACTATCGTCGCCGCACTTATGACTGTCTTCTTTATCATGCAACTCGTAGGACAGGTGCCGGCAGC
GCTCTTCCGCTTCCTCGCTCACTGACTCGCTGCGCTCGGTCGTTCGGCTGCGGCGAGCGGTATCAGCTC

ACTCAAAGGCGGTAATACGGTTATCCACAGAATCAGGGGATAACGCAGGAAAGAACATGTGAGCAAA
AGGCCAGCAAAAGGCCAGGAACCGTAAAAAGGCCGCGTTGCTGGCGTTTTTCCATAGGCTCCGCCCCC
CTGACGAGCATCACAAAAATCGACGCTCAAGTCAGAGGTGGCGAAACCCGACAGGACTATAAAGATA
CCAGGCGTTTCCCCCTGGAAGCTCCCTCGTGCGCTCTCCTGTTCCGACCCTGCCGCTTACCGGATACCT
GTCCGCCTTTCTCCCTTCGGGAAGCGTGGCGCTTTCTCATAGCTCACGCTGTAGGTATCTCAGTTCGGT
GTAGGTCGTTCGCTCCAAGCTGGGCTGTGTGCACGAACCCCCCGTTCAGCCCGACCGCTGCGCCTTATC
CGGTAACTATCGTCTTGAGTCCAACCCGGTAAGACACGACTTATCGCCACTGGCAGCAGCCACTGGTA
ACAGGATTAGCAGAGCGAGGTATGTAGGCGGTGCTACAGAGTTCTTGAAGTGGTGGCCTAACTACGGC
TACACTAGAAGAACAGTATTTGGTATCTGCGCTCTGCTGAAGCCAGTTACCTTCGGAAAAAGAGTTGG
TAGCTCTTGATCCGGCAAACAAACCACCGCTGGTAGCGGTGGTTTTTTTGTTTGCAAGCAGCAGATTAC
GCGCAGAAAAAAAGGATCTCAAGAAGATCCTTTGATCTTTTCTACGGGGTCTGACGCTCAGTGGAACG
AAAACTCACGTTAAGGGATTTTGGTCATGAGATTATCAAAAAGGATCTTCACCTAGATCCTTTTAAATT
AAAAATGAAGTTTTAAATCAATCTAAAGTATATATGAGTAAACTTGGTCTGACAGCGGCCGCAAATGC
TAAACCACTGCAGTGGTTACCAGTGCTTGATCAGTGAGGCACCGATCTCAGCGATCTGCCTATTTCGTT
CGTCCATAGTGGCCTGACTCCCCGTCGTGTAGATCACTACGATTCGTGAGGGCTTACCATCAGGCCCCA
GCGCAGCAATGATGCCGCGAGAGCCGCGTTCACCGGCCCCCGATTTGTCAGCAATGAACCAGCCAGCA
GGGAGGGCCGAGCGAAGAAGTGGTCCTGCTACTTTGTCCGCCTCCATCCAGTCTATGAGCTGCTGTCG
TGATGCTAGAGTAAGAAGTTCGCCAGTGAGTAGTTTCCGAAGAGTTGTGGCCATTGCTACTGGCATCG
TGGTATCACGCTCGTCGTTCGGTATGGCTTCGTTCAACTCTGGTTCCCAGCGGTCAAGCCGGGTCACAT
GATCACCCATATTATGAAGAAATGCAGTCAGCTCCTTAGGGCCTCCGATCGTTGTCAGAAGTAAGTTG
GCCGCGGTGTTGTCGCTCATGGTAATGGCAGCACTACACAATTCTCTTACCGTCATGCCATCCGTAAGA
TGCTTTTCCGTGACCGGCGAGTACTCAACCAAGTCGTTTTGTGAGTAGTGTATACGGCGACCAAGCTGC
TCTTGCCCGGCGTCTATACGGGACAACACCGCGCCACATAGCAGTACTTTGAAAGTGCTCATCATCGG
GAATCGTTCTTCGGGGCGGAAAGACTCAAGGATCTTGCCGCTATTGAGATCCAGTTCGATATAGCCCA
CTCTTGCACCCAGTTGATCTTCAGCATCTTTTACTTTCACCAGCGTTTCGGGGTGTGCAAAAACAGGCA
AGCAAAATGCCGCAAAGAAGGGAATGAGTGCGACACGAAAATGTTGGATGCTCATACTCGTCCTTTTT
CAATATTATTGAAGCATTTATCAGGGTTACTAGTACGTCTCTCAAGGATAAGTAAGTAATATTAAGGT
ACGGGAGGTATTGGACAGGCCGCAATAAAATATCTTTATTTTCATTACATCTGTGTGTTGGTTTTTTGT
GTGAATCGATAGTACTAACATACGCTCTCCATCAAAACAAAACGAAACAAAACAAACTAGCAAAATA
GGCTGTCCCCAGTGCAAGTGCAGGTGCCAGAACATTTCTCT 3′

CIRBP+MCRE-MHI (MCRE+CIRP+GFP+Neo-pGL4.10)
MCRE enhancer
CIRBP promoter
GGCCTAACTGGCCGGTACCttccccgccgttccccgccgttccccgccgttccccgccgttccccgccgTCTGGCTAACTAGAGAAC
CCACTGCGCCACAGTACCCATCTGTCTTCTGGCTTCACAAATGCGCCTCAGTTAAATAACTGTAATTAA
ATATTAAAAGAAAAAAAAGAACATAAAAGTGTTGAGTCGATGAGTCAAGGTTGGAGCCATATTTGGG
AGTCTCATTGAGGTAGCAAAAGCTCTTTGTTCCCCGCCGCAGGACAAACGGGTCGGGAACGCGGAAGT
GGGTCAGCTCCGCCTCCAAATCAGCTGTCCCCGCCTCCTAGTCGCTAGCGGGCGGGGGGTGGGGGGA
AAAATGCGGCCAGGAAAGATTCCAGCCAATCAAAACACTCAGCGCCTCCGCGCGGACTAGGGCAAAG
CCAACGGAGTCCCGCCCCTCCACCGAGGCGCAGATGCCGAATCTAAGGCGTCGGATTGGTCAGCGCGG
CGTGGTGGGCGGGTTAAGGGGCGTGGCTCCAGGAGTGGAGTATATCAGGCGGGACTCTTGCGCTCCCC
CTCGCACTCGCGCCTTAGGAAGCTTGGGTGTGTGTGGCGCGCTGTCTTCCCGCTCGCGTCAGGGACCTG
CCCGACTCAGCGGTGAGGGAGATCTGGCCTCGGCGGCCAAGCTTGGCAATCCGGTACTGTTGGTAAAG
CCACCATGGATGGTGAGCAAGGGCGCCGAGCTGTTCACCGGCATCGTGCCCATCCTGATCGAGCTGAA
TGGCGATGTGAATGGCCACAAGTTCAGCGTGAGCGGCGAGGGCGAGGGCGATGCCACCTACGGCAAG
CTGACCCTGAAGTTCATCTGCACCACCGGCAAGCTGCCTGTGCCCTGGCCCACCCTGGTGACCACCCTG
AGCTACGGCGTGCAGTGCTTCTCACGCTACCCCGATCACATGAAGCAGCACGACTTCTTCAAGAGCGC
CATGCCTGAGGGCTACATCCAGGAGCGCACCATCTTCTTCGAGGATGACGGCAACTACAAGTCGCGCG
CCGAGGTGAAGTTCGAGGGCGATACCCTGGTGAATCGCATCGAGCTGACCGGCACCGATTTCAAGGA
GGATGGCAACATCCTGGGCAATAAGATGGAGTACAACTACAACGCCCACAATGTGTACATCATGACC
GACAAGGCCAAGAATGGCATCAAGGTGAACTTCAAGATCCGCCACAACATCGAGGATGGCAGCGTGC
AGCTGGCCGACCACTACCAGCAGAATACCCCCATCGGCGATGGCCCTGTGCTGCTGCCCGATAACCAC
TACCTGTCCACCCAGAGCGCCCTGTCCAAGGACCCCAACGAGAAGCGCGATCACATGATCTACTTCGG
CTTCGTGACCGCCGCCGCCATCACCCACGGCATGGATGAGCTGTACAAGTGATCTAGAGTCGGGGCGG
CCGGCCGCTTCGAGCAGACATGATAAGATACATTGATGAGTTTGGACAAACCACAACTAGAATGCAGT

GAAAAAAATGCTTTATTTGTGAAATTTGTGATGCTATTGCTTTATTTGTAACCATTATAAGCTGCAATA
AACAAGTTAACAACAACAATTGCATTCATTTTATGTTTCAGGTTCAGGGGGAGGTGTGGGAGGTTTTTT
AAAGCAAGTAAAACCTCTACAAATGTGGTAAAATCGATAAGGATCCGTCGACtctaccgggtaggggaggcgctttt
cccaaggcagtctggagcatgcgctttagcagccccgctgggcacttggcgctacacaagtggcctctggcctcgcacacattccacatccaccggtaggcgccaacc
ggctccgttctttggtggccccttcgcgccaccttctactcctcccctagtcaggaagttcccccccgccccgcagctcgcgtcgtgcaggacgtgacaaatggaagtag
cacgtctcactagtctcgtgcagatggacagcaccgctgagcaatggaagcgggtaggcctttggggcagcggccaatagcagctttgctccttcgctttctgggctcag
aggctgggaaggggtgggtccgggggcgggctcaggggcgggctcaggggcggggcgggcgcccgaaggtcctccggaggcccggcattctgcacgcttcaaa
agcgcacgtctgccgcgctgttctcctcttcctcatctccgggcctttcgacctgcagccaatatgggatcggccattgaacaagatggattgcacgcaggttctccggcc
gcttgggtggagaggctattcggctatgactgggcacaacagacaatcggctgctctgatgccgccgtgttccggctgtcagcgcaggggcgcccggttctttttgtcaa
gaccgacctgtccggtgccctgaatgaactgcaggacgaggcagcgcggctatcgtggctggccacgacgggcgttccttgcgcagctgtgctcgacgttgtcactga
agcgggaagggactggctgctattgggcgaagtgccggggcaggatctcctgtcatctcaccttgctcctgccgagaaagtatccatcatggctgatgcaatgcggcgg
ctgcatacgcttgatccggctacctgcccattcgaccaccaagcgaaacatcgcatcgagcgagcacgtactcggatggaagccggtcttgtcgatcaggatgatctgga
cgaagagcatcaggggctcgcgccagccgaactgttcgccaggctcaaggcgcgcatgcccgacggcgatgatctcgtcgtgacccatggcgatgcctgcttgccga
atatcatggtggaaaatggccgcttttctggattcatcgactgtggccggctgggtgtggcggaccgctatcaggacatagcgttggctacccgtgatattgctgaagagct
tggcggcgaatgggctgaccgcttcctcgtgctttacggtatcgccgctcccgattcgcagcgcatcgccttctatcgccttcttgacgagttcttctgaggggatccgctgt
aagtctgcagaaattgatgatctattaaacaataaagatgtccactaaaatggaagtttttcctgtcatactttgttaagaagggtgagaacagagtacctacattttgaatgga
aggattggagctacgggggtgggggtggggtgggattagataaatgcctgctctttactgaaggctctttactattgctttatgataatgtttcatagttggatatcataatttaa
acaagcaaaaccaaattaagggccagctcattcctcccactcatgatctatagatctatagatctctcgtgggatcattgtttttctcttgattcccactttgtggttctaagtact
gtggtttccaaatgtgtcagtttcatagcctgaagaacgagatcagcagcctctgttccacatacacttcattctcagtattgttttgccaagttctaattccatcagaagcttG
ACTATCGTCGCCGCACTTATGACTGTCTTCTTTATCATGCAACTCGTAGGACAGGTGCCGGCAGCGCTC
TTCCGCTTCCTCGCTCACTGACTCGCTGCGCTCGGTCGTTCGGCTGCGGCGAGCGGTATCAGCTCACTC
AAAGGCGGTAATACGGTTATCCACAGAATCAGGGGATAACGCAGGAAAGAACATGTGAGCAAAAGGC
CAGCAAAAGGCCAGGAACCGTAAAAAGGCCGCGTTGCTGGCGTTTTTCCATAGGCTCCGCCCCCCTGA
CGAGCATCACAAAAATCGACGCTCAAGTCAGAGGTGGCGAAACCCGACAGGACTATAAAGATACCAG
GCGTTTCCCCCTGGAAGCTCCCTCGTGCGCTCTCCTGTTCCGACCCTGCCGCTTACCGGATACCTGTCC
GCCTTTCTCCCTTCGGGAAGCGTGGCGCTTTCTCATAGCTCACGCTGTAGGTATCTCAGTTCGGTGTAG
GTCGTTCGCTCCAAGCTGGGCTGTGTGCACGAACCCCCCGTTCAGCCCGACCGCTGCGCCTTATCCGGT
AACTATCGTCTTGAGTCCAACCCGGTAAGACACGACTTATCGCCACTGGCAGCAGCCACTGGTAACAG
GATTAGCAGAGCGAGGTATGTAGGCGGTGCTACAGAGTTCTTGAAGTGGTGGCCTAACTACGGCTACA
CTAGAAGAACAGTATTTGGTATCTGCGCTCTGCTGAAGCCAGTTACCTTCGGAAAAAGAGTTGGTAGC
TCTTGATCCGGCAAACAAACCACCGCTGGTAGCGGTGGTTTTTTTGTTTGCAAGCAGCAGATTACGCGC
AGAAAAAAAGGATCTCAAGAAGATCCTTTGATCTTTTCTACGGGGTCTGACGCTCAGTGGAACGAAAA
CTCACGTTAAGGGATTTTGGTCATGAGATTATCAAAAAGGATCTTCACCTAGATCCTTTTAAATTAAAA
ATGAAGTTTTAAATCAATCTAAAGTATATATGAGTAAACTTGGTCTGACAGCGGCCGCAAATGCTAAA
CCACTGCAGTGGTTACCAGTGCTTGATCAGTGAGGCACCGATCTCAGCGATCTGCCTATTTCGTTCGTC
CATAGTGGCCTGACTCCCCGTCGTGTAGATCACTACGATTCGTGAGGGCTTACCATCAGGCCCCAGCG
CAGCAATGATGCCGCGAGAGCCGCGTTCACCGGCCCCCGATTTGTCAGCAATGAACCAGCCAGCAGG
GAGGGCCGAGCGAAGAAGTGGTCCTGCTACTTTGTCCGCCTCCATCCAGTCTATGAGCTGCTGTCGTG
ATGCTAGAGTAAGAAGTTCGCCAGTGAGTAGTTTCCGAAGAGTTGTGGCCATTGCTACTGGCATCGTG
GTATCACGCTCGTCGTTCGGTATGGCTTCGTTCAACTCTGGTTCCCAGCGGTCAAGCCGGGTCACATGA
TCACCCATATTATGAAGAAATGCAGTCAGCTCCTTAGGGCCTCCGATCGTTGTCAGAAGTAAGTTGGC
CGCGGTGTTGTCGCTCATGGTAATGGCAGCACTACACAATTCTCTTACCGTCATGCCATCCGTAAGATG
CTTTTCCGTGACCGGCGAGTACTCAACCAAGTCGTTTTGTGAGTAGTGTATACGGCGACCAAGCTGCTC
TTGCCCGGCGTCTATACGGGACAACACCGCGCCACATAGCAGTACTTTGAAAGTGCTCATCATCGGGA
ATCGTTCTTCGGGGCGGAAAGACTCAAGGATCTTGCCGCTATTGAGATCCAGTTCGATATAGCCCACT
CTTGCACCCAGTTGATCTTCAGCATCTTTTACTTTCACCAGCGTTTCGGGGTGTGCAAAAACAGGCAAG
CAAAATGCCGCAAAGAAGGGAATGAGTGCGACACGAAAATGTTGGATGCTCATACTCGTCCTTTTTCA
ATATTATTGAAGCATTTATCAGGGTTACTAGTACGTCTCTCAAGGATAAGTAAGTAATATTAAGGTAC
GGGAGGTATTGGACAGGCCGCAATAAAATATCTTTATTTTCATTACATCTGTGTGTTGGTTTTTTGTGT
GAATCGATAGTACTAACATACGCTCTCCATCAAAACAAAACGAAACAAAACAAACTAGCAAAATAGG
CTGTCCCCAGTGCAAGTGCAGGTGCCAGAACATTTCTCT

RBM3-MHI (MCRE+RBM3+TurboGFP+Neo-pCMV6)
MCRE enhancer
*RBM3* promoter

5′AACAAAATATTAACGCTTACAATTTCCATTCGCCATTCAGGCTGCGCAACTGTTGGGAAGGGCGATC
GGTGCGGGCCTCTTCGCTATTACGCCAGCTGGCGAAAGGGGGATGTGCTGCAAGGCGATTAAGTTGGG
TAACGCCAGGGTTTTCCCAGTCACGACGTTGTAAAACGACGGCCAGTGCCAAGCTGATCTATACATTG
AATCAATATTGGCAATTAGCCATATTAGTCATTGGTTATATAGCATAAATCAATATTGGCTATTGGCCA
TTGCATACGTTGTATCTATATCATAATATGTACATTTATATTGGCTCATGTCCAATATGACCGCCATGTT
GACATTGATTATTGACTAGTTATTAATAGTAATCAATTACGGGGTCATTAGTTCATAGCCCATATATGG
AGTTCCGCGTTACATAACTTACGGTAAATGGCCCGCCTGGCTGACCGCCCAACGACCCCCGCCCATTG
ACGTCAATAATGACGTATGTTCCCATAGTAACGCCAATAGGGACTTTCCATTGACGTCAATGGGTGGA
GTATTTACGGTAAACTGCCCACTTGGCAGTACATCAAGTGTATCATATGCCAAGTCCGCCCCCTATTGA
CGTCAATGACGGTAAATGGCCCGCCTGGCATTATGCCCAGTACATGACCTTACGGGACTTTCCTACTTG
GCAGTACATCTACGTATTAGTCATCGCTATTACCATGGTGATGCGGTTTTGGCAGTACACCAATGGGCG
TGGATAGCGGTTTGACTCACGGGGATTTCCAAGTCTCCACCCCATTGACGTCAATGGGAGTTTGTTTTG
GCACCAAAATCAACGGGACTTTCCAAAATGTCGTAATAACCCCGCCCCGTTGACGCAAATGGGCGGTA
GGCGTGTACGGTGGGAGGTCTATATAAGCAGAGCTCGTTTAGTGAACCGTCAGAATTTTGTAATACGA
CTCACTATAGGGCGGCCGGGAATTCGTCGACTGGATCCGGTACCGAGGAGATCTGCCGCCGCGATCGC
CTTCCCCGCCGTTCCCCGCCGTTCCCCGCCGTTCCCCGCCGTTCCCCGCCGTTCCCCGCCGCTAACTTCG
ACCGTCAAGCCCCCTCGGTTCCAGCATCCTGTTTTGTAAAAGCCATCTCAGGCAGTCTATTGGTGAGAA
AATTAAGGGGCTGGCTTCACGCCAGCAGTCAGACAAGGGTGAGGGCTGAAGTCAGAGATGAGATGGA
ATGACGCCAGGGAAAAGGGTCCCATGACAGGCGGCTAGATGTTGAGCACAGAGGAGAAAGAGTAGG
AGTCAGGGTCAAACCTTAAAACGTTGGGTCTGACATCATGGACTGTGAAGGGTGATGACATCAGTGAC
GGTGGGGGATAAAATGTCAGGCCTTTAAAGGGTATGAGGTTAAAGAACTAGAAGGTGAAGGAGGTTA
ACATCAGAGAATGGCAGGGACCTAGAGGAAGACGACTACGATGGTGAGGAAGACATCCGATCCAGCG
GGGGAAATGATGGCAAGGACGGGAGGATGATGTCAGGGACAGCAAGTAGTATGAGATTGAGAAGGTT
CCTTTGTGGAAGGGATAAAGGCAGCTCAGCCGGTTGTGGGGGGGATGCTGACGTTTAGGGTGCACCAG
CTGTTCAAAAGTCGACCCCATTTCCGGAAACACCCCATTTCCGGAAACACCCCATTTCGCGGAGAACA
CTGCTCCCTTCGCTTTGCTGTCCCTTCCTCTCCCCACCACTCCGCTCCTGCTGGCCTCAGCCAATCAGCA
CGCACGCCGGGACGCGCAAGGGGAACGTTCCGGGACGTTCTCGCTACGTACTCTTTATCAATCGTCTT
CCGGCGCAGCCCCGTCCCTGTTTTTTGTGCTCCTCCGAGCTCGCTGTTCGTCCGGGTTTTTTACGTTTTA
ATTTCCAGGTAAGTTGCATATGTCCTGCGAAACGGCGGCTCCTCGCCACACACGCGCCGCGCCGGCCG
TGACCGGGGAATAGCGAGTGGGGAGAGTAATGGCCACTGACACGTACGCGTACGCGGCCGCTCGAGA
TGGAGAGCGACGAGAGCGGCCTGCCCGCCATGGAGATCGAGTGCCGCATCACCGGCACCCTGAACGG
CGTGGAGTTCGAGCTGGTGGGCGGCGGAGAGGGCACCCCCGAGCAGGGCCGCATGACCAACAAGATG
AAGAGCACCAAAGGCGCCCTGACCTTCAGCCCCTACCTGCTGAGCCACGTGATGGGCTACGGCTTCTA
CCACTTCGGCACCTACCCCAGCGGCTACGAGAACCCCTTCCTGCACGCCATCAACAACGGCGGCTACA
CCAACACCCGCATCGAGAAGTACGAGGACGGCGGCGTGCTGCACGTGAGCTTCAGCTACCGCTACGA
GGCCGGCCGCGTGATCGGCGACTTCAAGGTGATGGGCACCGGCTTCCCCGAGGACAGCGTGATCTTCA
CCGACAAGATCATCCGCAGCAACGCCACCGTGGAGCACCTGCACCCCATGGGCGATAACGATCTGGAT
GGCAGCTTCACCCGCACCTTCAGCCTGCGCGACGGCGGCTACTACAGCTCCGTGGTGGACAGCCACAT
GCACTTCAAGAGCGCCATCCACCCCAGCATCCTGCAGAACGGGGGCCCCATGTTCGCCTTCCGCCGCG
TGGAGGAGGATCACAGCAACACCGAGCTGGGCATCGTGGAGTACCAGCACGCCTTCAAGACCCCGGA
TGCAGATGCCGGTGAAGAAAGAGTTTAAACGGCCGGCCGCGGTCATAGCTGTTTCCTGAACAGATCCC
GGGTGGCATCCCTGTGACCCCTCCCCAGTGCCTCTCCTGGCCCTGGAAGTTGCCACTCCAGTGCCCACC
AGCCTTGTCCTAATAAAATTAAGTTGCATCATTTTGTCTGACTAGGTGTCCTTCTATAATATTATGGGG
TGGAGGGGGGTGGTATGGAGCAAGGGGCAAGTTGGGAAGACAACCTGTAGGGCCTGCGGGGTCTATT
GGGAACCAAGCTGGAGTGCAGTGGCACAATCTTGGCTCACTGCAATCTCCGCCTCCTGGGTTCAAGCG
ATTCTCCTGCCTCAGCCTCCCGAGTTGTTGGGATTCCAGGCATGCATGACCAGGCTCAGCTAATTTTTG
TTTTTTTGGTAGAGACGGGGTTTCACCATATTGGCCAGGCTGGTCTCCAACTCCTAATCTCAGGTGATC
TACCCACCTTGGCCTCCCAAATTGCTGGGATTACAGGCGTGAACCACTGCTCCCTTCCCTGTCCTTCTG
ATTTTAAAATAACTATACCAGCAGGAGGACGTCCAGACACAGCATAGGCTACCTGGCCATGCCCAACC
GGTGGGACATTTGAGTTGCTTGCTTGGCACTGTCCTCTCATGCGTTGGGTCCACTCAGTAGATGCCTGT
TGAATTGGGTACGCGGCCAGCTTGGCTGTGGAATGTGTGTCAGTTAGGGTGTGGAAAGTCCCCAGGCT
CCCCAGCAGGCAGAAGTATGCAAAGCATGCATCTCAATTAGTCAGCAACCAGGTGTGGAAAGTCCCC
AGGCTCCCCAGCAGGCAGAAGTATGCAAAGCATGCATCTCAATTAGTCAGCAACCATAGTCCCGCCCC
TAACTCCGCCCATCCCGCCCCTAACTCCGCCCAGTTCCGCCCATTCTCCGCCCCATGGCTGACTAATTT
TTTTTATTTATGCAGAGGCCGAGGCCGCCTCGGCCTCTGAGCTATTCCAGAAGTAGTGAGGAGGCTTTT
TTGGAGGCCTAGGCTTTTGCAAAAAGCTCCCGGGAGCTTGTATATCCATTTTCGGATCTGATCAAGAG

ACAGGATGAGGATCGTTTCGCATGATTGAACAAGATGGATTGCACGCAGGTTCTCCGGCCGCTTGGGT
GGAGAGGCTATTCGGCTATGACTGGGCACAACAGACAATCGGCTGCTCTGATGCCGCCGTGTTCCGGC
TGTCAGCGCAGGGGCGCCCGGTTCTTTTTGTCAAGACCGACCTGTCCGGTGCCCTGAATGAACTGCAG
GACGAGGCAGCGCGGCTATCGTGGCTGGCCACGACGGGCGTTCCTTGCGCAGCTGTGCTCGACGTTGT
CACTGAAGCGGGAAGGGACTGGCTGCTATTGGGCGAAGTGCCGGGGCAGGATCTCCTGTCATCTCACC
TTGCTCCTGCCGAGAAAGTATCCATCATGGCTGATGCAATGCGGCGGCTGCATACGCTTGATCCGGCT
ACCTGCCCATTCGACCACCAAGCGAAACATCGCATCGAGCGAGCACGTACTCGGATGGAAGCCGGTCT
TGTCGATCAGGATGATCTGGACGAAGAGCATCAGGGGCTCGCGCCAGCCGAACTGTTCGCCAGGCTCA
AGGCGCGCATGCCCGACGGCGAGGATCTCGTCGTGACCCATGGCGATGCCTGCTTGCCGAATATCATG
GTGGAAAATGGCCGCTTTTCTGGATTCATCGACTGTGGCCGGCTGGGTGTGGCCGACCGCTATCAGGA
CATAGCGTTGGCTACCCGTGATATTGCTGAAGAGCTTGGCGGCGAATGGGCTGACCGCTTCCTCGTGC
TTTACGGTATCGCCGCTCCCGATTCGCAGCGCATCGCCTTCTATCGCCTTCTTGACGAGTTCTTCTGAGC
GGGACTCTGGGGTTCGAAATGACCGACCAAGCGACGCCCAACCTGCCATCACGAGATTTCGATTCCAC
CGCCGCCTTCTATGAAAGGTTGGGCTTCGGAATCGTTTTCCGGGACGCCGGCTGGATGATCCTCCAGC
GCGGGGATCTCATGCTGGAGTTCTTCGCCCACCCCAACTTGTTTATTGCAGCTTATAATGGTTACAAAT
AAAGCAATAGCATCACAAATTTCACAAATAAAGCATTTTTTTCACTGCATTCTAGTTGTGGTTTGTCCA
AACTCATCAATGTATCTTATCATGTCTGTATACCGTCGACCTCTAGCTAGAGCTTGGCGTAATCATGGT
CATAGCTGTTTCCTGTGTGAAATTGTTATCCGCTCACAATTCCACACAACATACGAGCCGGAAGCATA
AAGTGTAAAGCCTGGGGTGCCTAATGAGTGAGCTAACTCACATTAATTGCGTTGCGCTCACTGCCCGC
TTTCCAGTCGGGAAACCTGTCGTGCCAGCTGCATTAATGAATCGGCCAACGCGCGGGGAGAGGCGGTT
TGCGTATTGGGCGCTCTTCCGCTTCCTCGCTCACTGACTCGCTGCGCTCGGTCGTTCGGCTGCGGCGAG
CGGTATCAGCTCACTCAAAGGCGGTAATACGGTTATCCACAGAATCAGGGGATAACGCAGGAAAGAA
CATGTGAGCAAAAGGCCAGCAAAAGGCCAGGAACCGTAAAAAGGCCGCGTTGCTGGCGTTTTTCCAT
AGGCTCCGCCCCCCTGACGAGCATCACAAAAATCGACGCTCAAGTCAGAGGTGGCGAAACCCGACAG
GACTATAAAGATACCAGGCGTTTCCCCCTGGAAGCTCCCTCGTGCGCTCTCCTGTTCCGACCCTGCCGC
TTACCGGATACCTGTCCGCCTTTCTCCCTTCGGGAAGCGTGGCGCTTTCTCATAGCTCACGCTGTAGGT
ATCTCAGTTCGGTGTAGGTCGTTCGCTCCAAGCTGGGCTGTGTGCACGAACCCCCCGTTCAGCCCGACC
GCTGCGCCTTATCCGGTAACTATCGTCTTGAGTCCAACCCGGTAAGACACGACTTATCGCCACTGGCA
GCAGCCACTGGTAACAGGATTAGCAGAGCGAGGTATGTAGGCGGTGCTACAGAGTTCTTGAAGTGGT
GGCCTAACTACGGCTACACTAGAAGAACAGTATTTGGTATCTGCGCTCTGCTGAAGCCAGTTACCTTC
GGAAAAAGAGTTGGTAGCTCTTGATCCGGCAAACAAACCACCGCTGGTAGCGGTTTTTTTGTTTGCAA
GCAGCAGATTACGCGCAGAAAAAAAGGATCTCAAGAAGATCCTTTGATCTTTTCTACGGGGTCTGACG
CTCAGTGGAACGAAAACTCACGTTAAGGGATTTTGGTCATGAGATTATCAAAAAGGATCTTCACCTAG
ATCCTTTTAAATTAAAAATGAAGTTTTAAATCAATCTAAAGTATATATGAGTAAACTTGGTCTGACAGT
TACCAATGCTTAATCAGTGAGGCACCTATCTCAGCGATCTGTCTATTTCGTTCATCCATAGTTGCCTGA
CTCCCCGTCGTGTAGATAACTACGATACGGGAGGGCTTACCATCTGGCCCCAGTGCTGCAATGATACC
GCGAGACCCACGCTCACCGGCTCCAGATTTATCAGCAATAAACCAGCCAGCCGGAAGGGCCGAGCGC
AGAAGTGGTCCTGCAACTTTATCCGCCTCCATCCAGTCTATTAATTGTTGCCGGGAAGCTAGAGTAAGT
AGTTCGCCAGTTAATAGTTTGCGCAACGTTGTTGCCATTGCTACAGGCATCGTGGTGTCACGCTCGTCG
TTTGGTATGGCTTCATTCAGCTCCGGTTCCCAACGATCAAGGCGAGTTACATGATCCCCCATGTTGTGC
AAAAAAGCGGTTAGCTCCTTCGGTCCTCCGATCGTTGTCAGAAGTAAGTTGGCCGCAGTGTTATCACTC
ATGGTTATGGCAGCACTGCATAATTCTCTTACTGTCATGCCATCCGTAAGATGCTTTTCTGTGACTGGT
GAGTACTCAACCAAGTCATTCTGAGAATAGTGTATGCGGCGACCGAGTTGCTCTTGCCCGGCGTCAAT
ACGGGATAATACCGCGCCACATAGCAGAACTTTAAAAGTGCTCATCATTGGAAAACGTTCTTCGGGGC
GAAAACTCTCAAGGATCTTACCGCTGTTGAGATCCAGTTCGATGTAACCCACTCGTGCACCCAACTGA
TCTTCAGCATCTTTTACTTTCACCAGCGTTTCTGGGTGAGCAAAAACAGGAAGGCAAAATGCCGCAAA
AAAGGGAATAAGGGCGACACGGAAATGTTGAATACTCATACTCTTCCTTTTTCAATATTATTGAAGCA
TTTATCAGGGTTATTGTCTCATGAGCGGATACATATTTGAATGTATTTAGAAAAATAAACAAATAGGG
GTTCCGCGCACATTTCCCCGAAAAGTGCCACCTGACGCGCCCTGTAGCGGCGCATTAAGCGCGGCGGG
TGTGGTGGTTACGCGCAGCGTGACCGCTACACTTGCCAGCGCCCTAGCGCCCGCTCCTTTCGCTTTCTT
CCCTTCCTTTCTCGCCACGTTCGCCGGCTTTCCCCGTCAAGCTCTAAATCGGGGGCTCCCTTTAGGGTTC
CGATTTAGTGCTTTACGGCACCTCGACCCCAAAAAACTTGATTAGGGTGATGGTTCACGTAGTGGGCC
ATCGCCCTGATAGACGGTTTTTCGCCCTTTGACGTTGGAGTCCACGTTCTTTAATAGTGGACTCTTGTTC
CAAACTGGAACAACACTCAACCCTATCTCGGTCTATTCTTTTGATTTATAAGGGATTTTGCCGATTTCG
GCCTATTGGTTAAAAAATGAGCTGATTTAACAAAAATTTAACGCGAATTTT 3′

SP1-MHI (SP1+GFP+Neo-pGL4.10)

*SP1* promoter

5′GGCCTAACTGGCCGGTACCTGAGCTCGCTAGCGCAACTTAGTCTCACACGCCTTGGAGAGCAAGCGA
GTCTTGCCATTGGATAATTCCACCGTCTTTCTTCTGCAAGTCCCTCCTTTCCCCCTCCCTCATTGGGCGG
GGCAGCAGAGAAGGGGCGGGGCCTAGGTTGGGCTTGTGGCGCGCTGCTCCCTCCTCCTTACCCCCCCC
TCCCTGTCCGGTCCGGGTTCGCTTGCCTCGTCAGCGTCCGCGTTTTTCCCGGCCCCCCCCAACCCCCCC
GGACAGGACCCCCTTGAGCCTCGAGGATATCAAGATCTGGCCTCGGCGGCCAAGCTTGGCAATCCGGT
ACTGTTGGTAAAGCCACCATGGATGGTGAGCAAGGGCGCCGAGCTGTTCACCGGCATCGTGCCCATCC
TGATCGAGCTGAATGGCGATGTGAATGGCCACAAGTTCAGCGTGAGCGGCGAGGGCGAGGGCGATGC
CACCTACGGCAAGCTGACCCTGAAGTTCATCTGCACCACCGGCAAGCTGCCTGTGCCCTGGCCCACCC
TGGTGACCACCCTGAGCTACGGCGTGCAGTGCTTCTCACGCTACCCCGATCACATGAAGCAGCACGAC
TTCTTCAAGAGCGCCATGCCTGAGGGCTACATCCAGGAGCGCACCATCTTCTTCGAGGATGACGGCAA
CTACAAGTCGCGCGCCGAGGTGAAGTTCGAGGGCGATACCCTGGTGAATCGCATCGAGCTGACCGGC
ACCGATTTCAAGGAGGATGGCAACATCCTGGGCAATAAGATGGAGTACAACTACAACGCCCACAATG
TGTACATCATGACCGACAAGGCCAAGAATGGCATCAAGGTGAACTTCAAGATCCGCCACAACATCGA
GGATGGCAGCGTGCAGCTGGCCGACCACTACCAGCAGAATACCCCCATCGGCGATGGCCCTGTGCTGC
TGCCCGATAACCACTACCTGTCCACCCAGAGCGCCCTGTCCAAGGACCCCAACGAGAAGCGCGATCAC
ATGATCTACTTCGGCTTCGTGACCGCCGCCGCCATCACCCACGGCATGGATGAGCTGTACAAGTGATC
TAGAGTCGGGGCGGCCGGCCGCTTCGAGCAGACATGATAAGATACATTGATGAGTTTGGACAAACCA
CAACTAGAATGCAGTGAAAAAAATGCTTTATTTGTGAAATTTGTGATGCTATTGCTTTATTTGTAACCA
TTATAAGCTGCAATAAACAAGTTAACAACAACAATTGCATTCATTTTATGTTTCAGGTTCAGGGGGAG
GTGTGGGAGGTTTTTTAAAGCAAGTAAAACCTCTACAAATGTGGTAAAATCGATAAGGATCCGTCGAC
tctaccgggtaggggaggcgcttttcccaaggcagtctggagcatgcgctttagcagccccgctgggcacttggcgctacacaagtggcctctggcctcgcacacattc
cacatccaccggtaggcgccaaccggctccgttctttggtggccccttcgcgccaccttctactcctcccctagtcaggaagttcccccccgccccgcagctcgcgtcgt
gcaggacgtgacaaatggaagtagcacgtctcactagtctcgtgcagatggacagcaccgctgagcaatggaagcgggtaggcctttggggcagcggccaatagcag
ctttgctccttcgctttctgggctcagaggctgggaaggggtgggtccgggggcgggctcaggggcgggctcaggggcggggcgggcgcccgaaggtcctccgga
ggcccggcattctgcacgcttcaaaagcgcacgtctgccgcgctgttctcctcttcctcatctccgggcctttcgacctgcagccaatatgggatcggccattgaacaagat
ggattgcacgcaggttctccggccgcttgggtggagaggctattcggctatgactgggcacaacagacaatcggctgctctgatgccgccgtgttccggctgtcagcgc
aggggcgcccggttctttttgtcaagaccgacctgtccggtgccctgaatgaactgcaggacgaggcagcgcggctatcgtggctggccacgacgggcgttccttgcg
cagctgtgctcgacgttgtcactgaagcgggaagggactggctgctattgggcgaagtgccggggcaggatctcctgtcatctcaccttgctcctgccgagaaagtatcc
atcatggctgatgcaatgcggcggctgcatacgcttgatccggctacctgcccattcgaccaccaagcgaaacatcgcatcgagcgagcacgtactcggatggaagcc
ggtcttgtcgatcaggatgatctggacgaagagcatcaggggctcgcgccagccgaactgttcgccaggctcaaggcgcgcatgcccgacggcgatgatctcgtcgtg
acccatggcgatgcctgcttgccgaatatcatggtggaaaatggccgcttttctggattcatcgactgtggccggctgggtgtggcggaccgctatcaggacatagcgttg
gctacccgtgatattgctgaagagcttggcggcgaatgggctgaccgcttcctcgtgctttacggtatcgccgctcccgattcgcagcgcatcgccttctatcgccttcttga
cgagttcttctgaggggatccgctgtaagtctgcagaaattgatgatctattaaacaataaagatgtccactaaaatggaagtttttcctgtcatactttgttaagaagggtgag
aacagagtacctacattttgaatggaaggattggagctacgggggtgggggtggggtgggattagataaatgcctgctctttactgaaggctctttactattgctttatgataa
tgtttcatagttggatatcataatttaaacaagcaaaaccaaattaagggccagctcattcctcccactcatgatctatagatctatagatctctcgtgggatcattgtttttctctt
gattcccactttgtggttctaagtactgtggtttccaaatgtgtcagtttcatagcctgaagaacgagatcagcagcctctgttccacatacacttcattctcagtattgttttgcc
aagttctaattccatcagaagcttGACTATCGTCGCCGCACTTATGACTGTCTTCTTTATCATGCAACTCGTAGGACAG
GTGCCGGCAGCGCTCTTCCGCTTCCTCGCTCACTGACTCGCTGCGCTCGGTCGTTCGGCTGCGGCGAGC
GGTATCAGCTCACTCAAAGGCGGTAATACGGTTATCCACAGAATCAGGGGATAACGCAGGAAAGAAC
ATGTGAGCAAAAGGCCAGCAAAAGGCCAGGAACCGTAAAAAGGCCGCGTTGCTGGCGTTTTTCCATA
GGCTCCGCCCCCCTGACGAGCATCACAAAAATCGACGCTCAAGTCAGAGGTGGCGAAACCCGACAGG
ACTATAAAGATACCAGGCGTTTCCCCCTGGAAGCTCCCTCGTGCGCTCTCCTGTTCCGACCCTGCCGCT
TACCGGATACCTGTCCGCCTTTCTCCCTTCGGGAAGCGTGGCGCTTTCTCATAGCTCACGCTGTAGGTA
TCTCAGTTCGGTGTAGGTCGTTCGCTCCAAGCTGGGCTGTGTGCACGAACCCCCCGTTCAGCCCGACCG
CTGCGCCTTATCCGGTAACTATCGTCTTGAGTCCAACCCGGTAAGACACGACTTATCGCCACTGGCAGC
AGCCACTGGTAACAGGATTAGCAGAGCGAGGTATGTAGGCGGTGCTACAGAGTTCTTGAAGTGGTGG
CCTAACTACGGCTACACTAGAAGAACAGTATTTGGTATCTGCGCTCTGCTGAAGCCAGTTACCTTCGG
AAAAAGAGTTGGTAGCTCTTGATCCGGCAAACAAACCACCGCTGGTAGCGGTGGTTTTTTTGTTTGCA
AGCAGCAGATTACGCGCAGAAAAAAAGGATCTCAAGAAGATCCTTTGATCTTTTCTACGGGGTCTGAC
GCTCAGTGGAACGAAAACTCACGTTAAGGGATTTTGGTCATGAGATTATCAAAAAGGATCTTCACCTA
GATCCTTTTAAATTAAAAATGAAGTTTTAAATCAATCTAAAGTATATATGAGTAAACTTGGTCTGACAG
CGGCCGCAAATGCTAAACCACTGCAGTGGTTACCAGTGCTTGATCAGTGAGGCACCGATCTCAGCGAT
CTGCCTATTTCGTTCGTCCATAGTGGCCTGACTCCCCGTCGTGTAGATCACTACGATTCGTGAGGGCTT
ACCATCAGGCCCCAGCGCAGCAATGATGCCGCGAGAGCCGCGTTCACCGGCCCCCGATTTGTCAGCAA

TGAACCAGCCAGCAGGGAGGGCCGAGCGAAGAAGTGGTCCTGCTACTTTGTCCGCCTCCATCCAGTCT
ATGAGCTGCTGTCGTGATGCTAGAGTAAGAAGTTCGCCAGTGAGTAGTTTCCGAAGAGTTGTGGCCAT
TGCTACTGGCATCGTGGTATCACGCTCGTCGTTCGGTATGGCTTCGTTCAACTCTGGTTCCCAGCGGTC
AAGCCGGGTCACATGATCACCCATATTATGAAGAAATGCAGTCAGCTCCTTAGGGCCTCCGATCGTTG
TCAGAAGTAAGTTGGCCGCGGTGTTGTCGCTCATGGTAATGGCAGCACTACACAATTCTCTTACCGTCA
TGCCATCCGTAAGATGCTTTTCCGTGACCGGCGAGTACTCAACCAAGTCGTTTTGTGAGTAGTGTATAC
GGCGACCAAGCTGCTCTTGCCCGGCGTCTATACGGGACAACACCGCGCCACATAGCAGTACTTTGAAA
GTGCTCATCATCGGGAATCGTTCTTCGGGGCGGAAAGACTCAAGGATCTTGCCGCTATTGAGATCCAG
TTCGATATAGCCCACTCTTGCACCCAGTTGATCTTCAGCATCTTTTACTTTCACCAGCGTTTCGGGGTGT
GCAAAAACAGGCAAGCAAAATGCCGCAAAGAAGGGAATGAGTGCGACACGAAAATGTTGGATGCTC
ATACTCGTCCTTTTTCAATATTATTGAAGCATTTATCAGGGTTACTAGTACGTCTCTCAAGGATAAGTA
AGTAATATTAAGGTACGGGAGGTATTGGACAGGCCGCAATAAAATATCTTTATTTTCATTACATCTGTG
TGTTGGTTTTTTGTGTGAATCGATAGTACTAACATACGCTCTCCATCAAAACAAAACGAAACAAAACA
AACTAGCAAAATAGGCTGTCCCCAGTGCAAGTGCAGGTGCCAGAACATTTCTCT 3′

SP1+MCRE-MHI (MCRE+SP1+GFP+Neo-PGL4.10)
MCRE enhancer
*SP1* promoter
GGCCTAACTGGCCGGTACCttccccgccgttccccgccgttccccgccgttccccgccgttccccgccgTCTGGCTAACTAGAGAAC
CCACTGCGCCACGAGCTCGCTAGCGCAACTTAGTCTCACACGCCTTGGAGAGCAAGCGAGTCTTGCCA
TTGGATAATTCCACCGTCTTTCTTCTGCAAGTCCCTCCTTTCCCCCTCCCTCATTGGGCGGGGCAGCAG
AGAAGGGGCGGGGCCTAGGTTGGGCTTGTGGCGCGCTGCTCCCTCCTCCTTACCCCCCCCTCCCTGTCC
GGTCCGGGTTCGCTTGCCTCGTCAGCGTCCGCGTTTTTCCCGGCCCCCCCCAACCCCCCCGGACAGGAC
CCCCTTGAGCCTCGAGGATATCAAGATCTGGCCTCGGCGGCCAAGCTTGGCAATCCGGTACTGTTGGT
AAAGCCACCATGGATGGTGAGCAAGGGCGCCGAGCTGTTCACCGGCATCGTGCCCATCCTGATCGAGC
TGAATGGCGATGTGAATGGCCACAAGTTCAGCGTGAGCGGCGAGGGCGAGGGCGATGCCACCTACGG
CAAGCTGACCCTGAAGTTCATCTGCACCACCGGCAAGCTGCCTGTGCCCTGGCCCACCCTGGTGACCA
CCCTGAGCTACGGCGTGCAGTGCTTCTCACGCTACCCCGATCACATGAAGCAGCACGACTTCTTCAAG
AGCGCCATGCCTGAGGGCTACATCCAGGAGCGCACCATCTTCTTCGAGGATGACGGCAACTACAAGTC
GCGCGCCGAGGTGAAGTTCGAGGGCGATACCCTGGTGAATCGCATCGAGCTGACCGGCACCGATTTCA
AGGAGGATGGCAACATCCTGGGCAATAAGATGGAGTACAACTACAACGCCCACAATGTGTACATCAT
GACCGACAAGGCCAAGAATGGCATCAAGGTGAACTTCAAGATCCGCCACAACATCGAGGATGGCAGC
GTGCAGCTGGCCGACCACTACCAGCAGAATACCCCCATCGGCGATGGCCCTGTGCTGCTGCCCGATAA
CCACTACCTGTCCACCCAGAGCGCCCTGTCCAAGGACCCCAACGAGAAGCGCGATCACATGATCTACT
TCGGCTTCGTGACCGCCGCCGCCATCACCCACGGCATGGATGAGCTGTACAAGTGATCTAGAGTCGGG
GCGGCCGGCCGCTTCGAGCAGACATGATAAGATACATTGATGAGTTTGGACAAACCACAACTAGAAT
GCAGTGAAAAAAATGCTTTATTTGTGAAATTTGTGATGCTATTGCTTTATTTGTAACCATTATAAGCTG
CAATAAACAAGTTAACAACAACAATTGCATTCATTTTATGTTTCAGGTTCAGGGGGAGGTGTGGGAGG
TTTTTTAAAGCAAGTAAAACCTCTACAAATGTGGTAAAATCGATAAGGATCCGTCGACtctaccgggtaggga
ggcgcttttcccaaggcagtctggagcatgcgctttagcagccccgctgggcacttggcgctacacaagtggcctctggcctcgcacacattccacatccaccggtagg
cgccaaccggctccgttctttggtggccccttcgcgccaccttctactcctcccctagtcaggaagttcccccccgccccgcagctcgcgtcgtgcaggacgtgacaaat
ggaagtagcacgtctcactagtctcgtgcagatggacagcaccgctgagcaatggaagcgggtaggcctttggggcagcggccaatagcagctttgctccttcgctttct
gggctcagaggctgggaaggggtgggtccgggggcgggctcaggggcgggctcaggggcggggcgggcgcccgaaggtcctccggaggcccggcattctgca
cgcttcaaaagcgcacgtctgccgcgctgttctcctcttcctcatctccgggcctttcgacctgcagccaatatgggatcggccattgaacaagatggattgcacgcaggtt
ctccggccgcttgggtggagaggctattcggctatgactgggcacaacagacaatcggctgctctgatgccgccgtgttccggctgtcagcgcaggggcgcccggttct
ttttgtcaagaccgacctgtccggtgccctgaatgaactgcaggacgaggcagcgcggctatcgtggctggccacgacgggcgttccttgcgcagctgtgctcgacgtt
gtcactgaagcgggaagggactggctgctattgggcgaagtgccggggcaggatctcctgtcatctcaccttgctcctgccgagaaagtatccatcatggctgatgcaat
gcggcggctgcatacgcttgatccggctacctgcccattcgaccaccaagcgaaacatcgcatcgagcgagcacgtactcggatggaagccggtcttgtcgatcaggat
gatctggacgaagagcatcaggggctcgcgccagccgaactgttcgccaggctcaaggcgcgcatgcccgacggcgatgatctcgtcgtgacccatggcgatgcctg
cttgccgaatatcatggtggaaaatggccgcttttctggattcatcgactgtggccggctgggtgtggcggaccgctatcaggacatagcgttggctacccgtgatattgct
gaagagcttggcggcgaatgggctgaccgcttcctcgtgctttacggtatcgccgctcccgattcgcagcgcatcgccttctatcgccttcttgacgagttcttctgagggg
atccgctgtaagtctgcagaaattgatgatctattaaacaataaagatgtccactaaaatggaagtttttcctgtcatactttgttaagaagggtgagaacagagtacctacattt
tgaatggaaggattggagctacgggggtgggggtggggtgggattagataaatgcctgctctttactgaaggctctttactattgctttatgataatgtttcatagttggatatc
ataatttaaacaagcaaaaccaaattaagggccagctcattcctcccactcatgatctatagatctatagatctctcgtgggatcattgtttttctcttgattcccactttgtggttct
aagtactgtggtttccaaatgtgtcagtttcatagcctgaagaacgagatcagcagcctctgttccacatacacttcattctcagtattgttttgccaagttctaattccatcagaa
gcttGACTATCGTCGCCGCACTTATGACTGTCTTCTTTATCATGCAACTCGTAGGACAGGTGCCGGCAGC

GCTCTTCCGCTTCCTCGCTCACTGACTCGCTGCGCTCGGTCGTTCGGCTGCGGCGAGCGGTATCAGCTC
ACTCAAAGGCGGTAATACGGTTATCCACAGAATCAGGGGATAACGCAGGAAAGAACATGTGAGCAAA
AGGCCAGCAAAAGGCCAGGAACCGTAAAAAGGCCGCGTTGCTGGCGTTTTTCCATAGGCTCCGCCCCC
CTGACGAGCATCACAAAAATCGACGCTCAAGTCAGAGGTGGCGAAACCCGACAGGACTATAAAGATA
CCAGGCGTTTCCCCCTGGAAGCTCCCTCGTGCGCTCTCCTGTTCCGACCCTGCCGCTTACCGGATACCT
GTCCGCCTTTCTCCCTTCGGGAAGCGTGGCGCTTTCTCATAGCTCACGCTGTAGGTATCTCAGTTCGGT
GTAGGTCGTTCGCTCCAAGCTGGGCTGTGTGCACGAACCCCCCGTTCAGCCCGACCGCTGCGCCTTATC
CGGTAACTATCGTCTTGAGTCCAACCCGGTAAGACACGACTTATCGCCACTGGCAGCAGCCACTGGTA
ACAGGATTAGCAGAGCGAGGTATGTAGGCGGTGCTACAGAGTTCTTGAAGTGGTGGCCTAACTACGGC
TACACTAGAAGAACAGTATTTGGTATCTGCGCTCTGCTGAAGCCAGTTACCTTCGGAAAAAGAGTTGG
TAGCTCTTGATCCGGCAAACAAACCACCGCTGGTAGCGGTGGTTTTTTTGTTTGCAAGCAGCAGATTAC
GCGCAGAAAAAAAGGATCTCAAGAAGATCCTTTGATCTTTTCTACGGGGTCTGACGCTCAGTGGAACG
AAAACTCACGTTAAGGGATTTTGGTCATGAGATTATCAAAAAGGATCTTCACCTAGATCCTTTTAAATT
AAAAATGAAGTTTTAAATCAATCTAAAGTATATATGAGTAAACTTGGTCTGACAGCGGCCGCAAATGC
TAAACCACTGCAGTGGTTACCAGTGCTTGATCAGTGAGGCACCGATCTCAGCGATCTGCCTATTTCGTT
CGTCCATAGTGGCCTGACTCCCCGTCGTGTAGATCACTACGATTCGTGAGGGCTTACCATCAGGCCCCA
GCGCAGCAATGATGCCGCGAGAGCCGCGTTCACCGGCCCCCGATTTGTCAGCAATGAACCAGCCAGCA
GGGAGGGCCGAGCGAAGAAGTGGTCCTGCTACTTTGTCCGCCTCCATCCAGTCTATGAGCTGCTGTCG
TGATGCTAGAGTAAGAAGTTCGCCAGTGAGTAGTTTCCGAAGAGTTGTGGCCATTGCTACTGGCATCG
TGGTATCACGCTCGTCGTTCGGTATGGCTTCGTTCAACTCTGGTTCCCAGCGGTCAAGCCGGGTCACAT
GATCACCCATATTATGAAGAAATGCAGTCAGCTCCTTAGGGCCTCCGATCGTTGTCAGAAGTAAGTTG
GCCGCGGTGTTGTCGCTCATGGTAATGGCAGCACTACACAATTCTCTTACCGTCATGCCATCCGTAAGA
TGCTTTTCCGTGACCGGCGAGTACTCAACCAAGTCGTTTTGTGAGTAGTGTATACGGCGACCAAGCTGC
TCTTGCCCGGCGTCTATACGGGACAACACCGCGCCACATAGCAGTACTTTGAAAGTGCTCATCATCGG
GAATCGTTCTTCGGGGCGGAAAGACTCAAGGATCTTGCCGCTATTGAGATCCAGTTCGATATAGCCCA
CTCTTGCACCCAGTTGATCTTCAGCATCTTTTACTTTCACCAGCGTTTCGGGGTGTGCAAAAACAGGCA
AGCAAAATGCCGCAAAGAAGGGAATGAGTGCGACACGAAAATGTTGGATGCTCATACTCGTCCTTTTT
CAATATTATTGAAGCATTTATCAGGGTTACTAGTACGTCTCTCAAGGATAAGTAAGTAATATTAAGGT
ACGGGAGGTATTGGACAGGCCGCAATAAAATATCTTTATTTTCATTACATCTGTGTGTTGGTTTTTTGT
GTGAATCGATAGTACTAACATACGCTCTCCATCAAAACAAAACGAAACAAAACAAACTAGCAAAATA
GGCTGTCCCCAGTGCAAGTGCAGGTGCCAGAACATTTCTCT

CIRBP-Lenti-MHI (pLV[Exp]-Neo-{MCRE_spacer_CIRP_promorer}>mCherry):
MCRE enhancer
*CIRBP* promoter
aatgtagtcttatgcaatactcttgtagtcttgcaacatggtaacgatgagttagcaacatgccttacaaggagagaaaaagcaccgtgcatgccgattggtggaagtaagg
tggtacgatcgtgccttattaggaaggcaacagacgggtctgacatggattggacgaaccactgaattgccgcattgcagagatattgtatttaagtgcctagctcgataca
taaacgggtctctctggttagaccagatctgagcctgggagctctctggctaactagggaacccactgcttaagcctcaataaagcttgccttgagtgcttcaagtagtgtgt
gcccgtctgttgtgtgactctggtaactagagatccctcagacccttttagtcagtgtggaaaatctctagcagtggcgcccgaacagggacttgaaagcgaaagggaaa
ccagaggagctctctcgacgcaggactcggcttgctgaagcgcgcacggcaagaggcgaggggcggcgactggtgagtacgccaaaaattttgactagcggaggct
agaaggagagagatgggtgcgagagcgtcagtattaagcgggggagaattagatcgcgatgggaaaaaattcggttaaggccagggggaaagaaaaaatataaatta
aaacatatagtatgggcaagcagggagctagaacgattcgcagttaatcctggcctgttagaaacatcagaaggctgtagacaaatactgggacagctacaaccatccct
tcagacaggatcagaagaacttagatcattatataatacagtagcaaccctctattgtgtgcatcaaaggatagagataaaagacaccaaggaagctttagacaagataga
ggaagagcaaaacaaaagtaagaccaccgcacagcaagcggccgctgatcttcagacctggaggaggagatatgagggacaattggagaagtgaattatataaatata
aagtagtaaaaattgaaccattaggagtagcacccaccaaggcaaagagaagagtggtgcagagagaaaaaagagcagtgggaataggagctttgttccttgggttctt
gggagcagcaggaagcactatgggcgcagcgtcaatgacgctgacggtacaggccagacaattattgtctggtatagtgcagcagcagaacaatttgctgagggctatt
gaggcgcaacagcatctgttgcaactcacagtctggggcatcaagcagctccaggcaagaatcctggctgtggaaagatacctaaaggatcaacagctcctggggattt
ggggttgctctggaaaactcatttgcaccactgctgtgccttggaatgctagttggagtaataaatctctggaacagatttggaatcacacgacctggatggagtgggacag
agaaattaacaattacacaagcttaatacactccttaattgaagaatcgcaaaaccagcaagaaaagaatgaacaagaattattggaattagataaatgggcaagtttgtgg
aattggtttaacataacaaattggctgtggtatataaaattattcataatgatagtaggaggcttggtaggtttaagaatagtttttgctgtactttctatagtgaatagagttaggc
agggatattcaccattatcgtttcagacccacctcccaaccccgaggggacccgacaggcccgaaggaatagaagaagaaggtggagagagagacagagacagatc
cattcgattagtgaacggatctcgacggtatcgctagcttttaaaagaaaaggggggattggggggtacagtgcaggggaaagaatagtagacataatagcaacagaca
tacaaactaaagaattacaaaaacaaattacaaaaattcaaaattttactagtgattatcggatcaactttgtatagaaaagttgttccccgccgttccccgccgttccccgccg
ttccccgccgttccccgccgttccccgccgttccccgccgttccccgccgtctggtggcggaggctcgggcggaggtgggtcgggtggcggcggatcagctgggaaa
actgggacgtcggttgtattggcaagaatatcacgacaggtgtcggggttcagatgaagttgggatgtctgacagccgggtcttggttgggtaggggatctgggctaggtt
gttatgtcgggactgctggagcacaggcaaagtcggggtcccagtgacacccagaagtccccatgaggcctgaggcctgggaatcggggctagaggcccatttgttcc

ccgccgcaggccagggagcaggcacgcgtaactgggtcagccccgccccgccccgccgcgcgctggtcacgccccctcgagcgcgggtggaggactgcggact
gcaaacgtctcggccaatagtcgagcccgacaccagtccgcgggagggggcggaaccgagatagactcggcccccagctcatgaatataagacgcaggcgcgccc
tctgattggcccaggatgtgtagggggcggggcccggcggaagcgtatataaggccgggctcggggacgcccccccctcactcgcgcgttaggaggctcgggtcgtt
gtggtgcgctgtcttcccgcttgcgtcagggacctgcccgactcagtggtgagggccccaagtttgtacaaaaaagcaggctgccaccatggtgagcaagggcgagga
ggataacatggccatcatcaaggagttcatgcgcttcaaggtgcacatggagggctccgtgaacggccacgagttcgagatcgagggcgagggcgagggccgcccc
tacgagggcacccagaccgccaagctgaaggtgaccaagggtggccccctgcccttcgcctgggacatcctgtcccctcagttcatgtacggctccaaggcctacgtg
aagcaccccgccgacatccccgactacttgaagctgtccttccccgagggcttcaagtgggagcgcgtgatgaacttcgaggacggcggcgtggtgaccgtgaccca
ggactcctccctgcaggacggcgagttcatctacaaggtgaagctgcgcggcaccaacttcccctccgacggccccgtaatgcagaagaagaccatgggctgggagg
cctcctccgagcggatgtaccccgaggacggcgccctgaagggcgagatcaagcagaggctgaagctgaaggacggcggccactacgacgctgaggtcaagacca
cctacaaggccaagaagcccgtgcagctgcccggcgcctacaacgtcaacatcaagttggacatcacctcccacaacgaggactacaccatcgtggaacagtacgaa
cgcgccgagggccgccactccaccggcggcatggacgagctgtacaagtaaacccagctttcttgtacaaagtggtgataatcgaattccgataatcaacctctggatta
caaaatttgtgaaagattgactggtattcttaactatgttgctccttttacgctatgtggatacgctgctttaatgcctttgtatcatgctattgcttcccgtatggctttcattttctcct
ccttgtataaatcctggttgctgtctctttatgaggagttgtggcccgttgtcaggcaacgtggcgtggtgtgcactgtgtttgctgacgcaacccccactggttggggcattg
ccaccacctgtcagctcctttccgggactttcgctttccccctccctattgccacggcggaactcatcgccgcctgccttgcccgctgctggacaggggctcggctgttgg
gcactgacaattccgtggtgttgtcggggaagctgacgtcctttccatggctgctcgcctgtgttgccacctggattctgcgcgggacgtccttctgctacgtcccttcggcc
ctcaatccagcggaccttccttcccgcggcctgctgccggctctgcggcctcttccgcgtcttcgccttcgccctcagacgagtcggatctccctttgggccgcctccccg
catcgggaattcccgcggttcgaattctaccgggtaggggaggcgcttttcccaaggcagtctggagcatgcgctttagcagccccgctgggcacttggcgctacacaa
gtggcctctggcctcgcacacattccacatccaccggtaggcgccaaccggctccgttctttggtggccccttcgcgccaccttctactcctcccctagtcaggaagttccc
ccccgccccgcagctcgcgtcgtgcaggacgtgacaaatggaagtagcacgtctcactagtctcgtgcagatggacagcaccgctgagcaatggaagcgggtaggcc
tttggggcagcggccaatagcagctttgctccttcgctttctgggctcagaggctgggaaggggtgggtccgggggcgggctcaggggcgggctcaggggcggggc
gggcgcccgaaggtcctccggaggcccggcattctgcacgcttcaaaagcgcacgtctgccgcgctgttctcctcttcctcatctccgggcctttcgacctcacgtgcgc
atgattgaacaagatggattgcacgcaggttctccggccgcttgggtggagaggctattcggctatgactgggcacaacagacaatcggctgctctgatgccgccgtgtt
ccggctgtcagcgcaggggcgcccggttctttttgtcaagaccgacctgtccggtgccctgaatgaactgcaagacgaggcagcgcggctatcgtggctggccacgac
gggcgttccttgcgcagctgtgctcgacgttgtcactgaagcgggaagggactggctgctattgggcgaagtgccggggcaggatctcctgtcatctcaccttgctcctg
ccgagaaagtatccatcatggctgatgcaatgcggcggctgcatacgcttgatccggctacctgcccattcgaccaccaagcgaaacatcgcatcgagcgagcacgtac
tcggatggaagccggtcttgtcgatcaggatgatctggacgaagagcatcaggggctcgcgccagccgaactgttcgccaggctcaaggcgagcatgcccgacggc
gaggatctcgtcgtgacccatggcgatgcctgcttgccgaatatcatggtggaaaatggccgcttttctggattcatcgactgtggccggctgggtgtggcggaccgctat
caggacatagcgttggctacccgtgatattgctgaagagcttggcggcgaatgggctgaccgcttcctcgtgctttacggtatcgccgctcccgattcgcagcgcatcgc
cttctatcgccttcttgacgagttcttctgagcgggactctgggtacctttaagaccaatgacttacaaggcagctgtagatcttagccactttttaaaagaaaaggggggact
ggaagggctaattcactcccaacgaagacaagatctgctttttgcttgtactgggtctctctggttagaccagatctgagcctgggagctctctggctaactagggaaccca
ctgcttaagcctcaataaagcttgccttgagtgcttcaagtagtgtgtgcccgtctgttgtgtgactctggtaactagagatccctcagacccttttagtcagtgtggaaaatct
ctagcagtagtagttcatgtcatcttattattcagtatttataacttgcaaagaaatgaatatcagagagtgagaggaacttgtttattgcagcttataatggttacaaataaagca
atagcatcacaaatttcacaaataaagcatttttttcactgcattctagttgtggtttgtccaaactcatcaatgtatcttatcatgtctggctctagctatcccgcccctaactccg
cccatcccgcccctaactccgcccagttccgcccattctccgccccatggctgactaattttttttatttatgcagaggccgaggccgcctcggcctctgagctattccagaa
gtagtgaggaggcttttttggaggcctagggacgtacccaattcgccctatagtgagtcgtattacgcgcgctcactggccgtcgttttacaacgtcgtgactgggaaaacc
ctggcgttacccaacttaatcgccttgcagcacatccccctttcgccagctggcgtaatagcgaagaggcccgcaccgatcgcccttcccaacagttgcgcagcctgaat
ggcgaatgggacgcgccctgtagcggcgcattaagcgcggcgggtgtggtggttacgcgcagcgtgaccgctacacttgccagcgccctagcgcccgctcctttcgct
ttcttcccttcctttctcgccacgttcgccggctttccccgtcaagctctaaatcgggggctccctttagggttccgatttagtgctttacggcacctcgaccccaaaaaacttg
attagggtgatggttcacgtagtgggccatcgccctgatagacggtttttcgccctttgacgttggagtccacgttctttaatagtggactcttgttccaaactggaacaacact
caaccctatctcggtctattcttttgatttataagggattttgccgatttcggcctattggttaaaaaatgagctgatttaacaaaaatttaacgcgaattttaacaaaatattaacg
cttacaatttaggtggcacttttcggggaaatgtgcgcggaacccctatttgtttatttttctaaatacattcaaatatgtatccgctcatgagacaataaccctgataaatgcttc
aataatattgaaaaaggaagagtatgagtattcaacatttccgtgtcgcccttattcccttttttgcggcattttgccttcctgtttttgctcacccagaaacgctggtgaaagtaa
aagatgctgaagatcagttgggtgcacgagtgggttacatcgaactggatctcaacagcggtaagatccttgagagttttcgccccgaagaacgttttccaatgatgagca
cttttaaagttctgctatgtggcgcggtattatcccgtattgacgccgggcaagagcaactcggtcgccgcatacactattctcagaatgacttggttgagtactcaccagtc
acagaaaagcatcttacggatggcatgacagtaagagaattatgcagtgctgccataaccatgagtgataacactgcggccaacttacttctgacaacgatcggaggacc
gaaggagctaaccgcttttttgcacaacatgggggatcatgtaactcgccttgatcgttgggaaccggagctgaatgaagccataccaaacgacgagcgtgacaccacg
atgcctgtagcaatggcaacaacgttgcgcaaactattaactggcgaactacttactctagcttcccggcaacaattaatagactggatggaggcggataaagttgcagga
ccacttctgcgctcggcccttccggctggctggtttattgctgataaatctggagccggtgagcgtgggtctcgcggtatcattgcagcactggggccagatggtaagccc
tcccgtatcgtagttatctacacgacggggagtcaggcaactatggatgaacgaaatagacagatcgctgagataggtgcctcactgattaagcattggtaactgtcagac
caagtttactcatatatactttagattgatttaaaacttcatttttaatttaaaaggatctaggtgaagatcctttttgataatctcatgaccaaaatcccttaacgtgagttttcgttcc
actgagcgtcagaccccgtagaaaagatcaaaggatcttcttgagatcctttttttctgcgcgtaatctgctgcttgcaaacaaaaaaaccaccgctaccagcggtggtttgtt
tgccggatcaagagctaccaactctttttccgaaggtaactggcttcagcagagcgcagataccaaatactgttcttctagtgtagccgtagttaggccaccacttcaagaa
ctctgtagcaccgcctacatacctcgctctgctaatcctgttaccagtggctgctgccagtggcgataagtcgtgtcttaccgggttggactcaagacgatagttaccggata
aggcgcagcggtcgggctgaacggggggttcgtgcacacagcccagcttggagcgaacgacctacaccgaactgagatacctacagcgtgagctatgagaaagcgc
cacgcttcccgaagagagaaaggcggacaggtatccggtaagcggcagggtcggaacaggagagcgcacgagggagcttccagggggaaacgcctggtatctttat
agtcctgtcgggtttcgccacctctgacttgagcgtcgatttttgtgatgctcgtcaggggggcggagcctatggaaaaacgccagcaacgcggcctttttacggttcctgg

cctttgctggccttttgctcacatgttctttcctgcgttatcccctgattctgtggataaccgtattaccgcctttgagtgagctgataccgctcgccgcagccgaacgaccga
gcgcagcgagtcagtgagcgaggaagcggaagagcgcccaatacgcaaaccgcctctccccgcgcgttggccgattcattaatgcagctggcacgacaggtttcccg
actggaaagcgggcagtgagcgcaacgcaattaatgtgagttagctcactcattaggcaccccaggctttacactttatgcttccggctcgtatgttgtgtggaattgtgagc
ggataacaatttcacacaggaaacagctatgaccatgattacgccaagcgcgcaattaaccctcactaaagggaacaaaagctggagctgcaagctt

RBM3-Lenti-MHI (pLV[Exp]-Neo-{MCRE_spacer_RBM3_promorer}>mCherry):
MCRE enhancer
*RBM3* promoter
aatgtagtcttatgcaatactcttgtagtcttgcaacatggtaacgatgagttagcaacatgccttacaaggagagaaaaagcaccgtgcatgccgattggtggaagtaagg
tggtacgatcgtgccttattaggaaggcaacagacgggtctgacatggattggacgaaccactgaattgccgcattgcagagatattgtatttaagtgcctagctcgataca
taaacgggtctctctggttagaccagatctgagcctgggagctctctggctaactagggaacccactgcttaagcctcaataaagcttgccttgagtgcttcaagtagtgtgt
gcccgtctgttgtgtgactctggtaactagagatccctcagacccttttagtcagtgtggaaaatctctagcagtggcgcccgaacagggacttgaaagcgaaagggaaa
ccagaggagctctctcgacgcaggactcggcttgctgaagcgcgcacggcaagaggcgaggggcggcgactggtgagtacgccaaaaattttgactagcggaggct
agaaggagagagatgggtgcgagagcgtcagtattaagcgggggagaattagatcgcgatgggaaaaaattcggttaaggccagggggaaagaaaaaatataaatta
aaacatatagtatgggcaagcagggagctagaacgattcgcagttaatcctggcctgttagaaacatcagaaggctgtagacaaatactgggacagctacaaccatccct
tcagacaggatcagaagaacttagatcattatataatacagtagcaaccctctattgtgtgcatcaaaggatagagataaaagacaccaaggaagctttagacaagataga
ggaagagcaaaacaaaagtaagaccaccgcacagcaagcggccgctgatcttcagacctggaggaggagatatgagggacaattggagaagtgaattatataaatata
aagtagtaaaaattgaaccattaggagtagcacccaccaaggcaaagagaagagtggtgcagagagaaaaaagagcagtgggaataggagctttgttccttgggttctt
gggagcagcaggaagcactatgggcgcagcgtcaatgacgctgacggtacaggccagacaattattgtctggtatagtgcagcagcagaacaatttgctgagggctatt
gaggcgcaacagcatctgttgcaactcacagtctggggcatcaagcagctccaggcaagaatcctggctgtggaaagatacctaaaggatcaacagctcctggggattt
ggggttgctctggaaaactcatttgcaccactgctgtgccttggaatgctagttggagtaataaatctctggaacagatttggaatcacacgacctggatggagtgggacag
agaaattaacaattacacaagcttaatacactccttaattgaagaatcgcaaaaccagcaagaaaagaatgaacaagaattattggaattagataaatgggcaagtttgtgg
aattggtttaacataacaaattggctgtggtatataaaattattcataatgatagtaggaggcttggtaggtttaagaatagtttttgctgtactttctatagtgaatagagttaggc
agggatattcaccattatcgtttcagacccacctcccaaccccgaggggacccgacaggcccgaaggaatagaagaagaaggtggagagagagacagagacagatc
cattcgattagtgaacggatctcgacggtatcgctagcttttaaaagaaaaggggggattggggggtacagtgcaggggaaagaatagtagacataatagcaacagaca
tacaaactaaagaattacaaaaacaaattacaaaaattcaaaattttactagtgattatcggatcaactttgtatagaaaagttgttccccgccgttccccgccgttccccgccg
ttccccgccgttccccgccgtctggtggcggaggctcgggcggaggtgggtcgggtggcggcggatcacagcatcctgttttgtaaaagccatctcaggcagtctattg
gtgagaaaattaaggggctggcttcacgccagcagtcagacaagggtgagggctgaagtcagagatgagatggaatgacgccagggaaaagggtcccatgacaggc
ggctagatgttgagcacagaggagaaagagtaggagtcagggtcaaaccttaaaacgttgggtctgacatcatggactgtgaagggtgatgacatcagtgacggtggg
ggataaaatgtcaggcctttaaagggtatgaggttaaagaactagaaggtgaaggaggttaacatcagagaatggcagggacctagaggaagacgactacgatggtga
ggaagacatccgatccagcgggggaaatgatggcaaggacgggaggatgatgtcagggacagcaagtagtatgagattgagaaggttcctttgtggaagggataaag
gcagctcagccggttgtggggggggatgctgacgtttagggtgcaccagctgttcaaaagtcgaccccatttccggaaacaccccatttccggaaacaccccatttcgcgg
agaacactgctcccttcgctttgctgtcccttcctctccccaccactccgctcctgctggcctcagccaatcagcacgcacgccgggacgcgcaaggggaacgttccggg
acgttctcgctacgtactctttatcaatcgtcttccggcgcagccccgtccctgtttttgtgctcctccgagctcgctgttcgtccgggtttttacgttttaatttccaggtaagtt
gcatatgtcctgcgaaacggcggctcctcgccacacacgcgccgcgccggccgtgaccggggaatagcgagtggggagagtaatggccactgacacgtcaagtttgt
acaaaaaagcaggctgccaccatggtgagcaagggcgaggaggataacatggccatcatcaaggagttcatgcgcttcaaggtgcacatggagggctccgtgaacg
gccacgagttcgagatcgagggcgagggcgagggccgcccctacgagggcacccagaccgccaagctgaaggtgaccaagggtggccccctgcccttcgcctgg
gacatcctgtcccctcagttcatgtacggctccaaggcctacgtgaagcaccccgccgacatccccgactacttgaagctgtccttccccgagggcttcaagtgggagcg
cgtgatgaacttcgaggacggcggcgtggtgaccgtgacccaggactcctccctgcaggacggcgagttcatctacaaggtgaagctgcgcggcaccaacttcccctc
cgacggccccgtaatgcagaagaagaccatgggctgggaggcctcctccgagcggatgtaccccgaggacggcgccctgaagggcgagatcaagcagaggctga
agctgaaggacggcggccactacgacgctgaggtcaagaccacctacaaggccaagaagcccgtgcagctgcccggcgcctacaacgtcaacatcaagttggacat
cacctcccacaacgaggactacaccatcgtggaacagtacgaacgcgccgagggccgccactccaccggcggcatggacgagctgtacaagtaaacccagctttctt
gtacaaagtggtgataatcgaattccgataatcaacctctggattacaaaatttgtgaaagattgactggtattcttaactatgttgctccttttacgctatgtggatacgctgcttt
aatgcctttgtatcatgctattgcttcccgtatggctttcattttctcctccttgtataaatcctggttgctgtctctttatgaggagttgtggcccgttgtcaggcaacgtggcgtg
gtgtgcactgtgtttgctgacgcaacccccactggttggggcattgccaccacctgtcagctcctttccgggactttcgctttccccctccctattgccacggcggaactcat
cgccgcctgccttgcccgctgctggacaggggctcggctgttgggcactgacaattccgtggtgttgtcggggaagctgacgtcctttccatggctgctcgcctgtgttgc
cacctggattctgcgcgggacgtccttctgctacgtcccttcggccctcaatccagcggaccttccttcccgcggcctgctgccggctctgcggcctcttccgcgtcttcgc
cttcgccctcagacgagtcggatctccctttgggccgcctccccgcatcgggaattcccgcggttcgaattctaccgggtaggggaggcgcttttcccaaggcagtctgg
agcatgcgctttagcagccccgctgggcacttggcgctacacaagtggcctctggcctcgcacacattccacatccaccggtaggcgccaaccggctccgttctttggtg
gccccttcgcgccaccttctactcctcccctagtcaggaagttcccccccgccccgcagctcgcgtcgtgcaggacgtgacaaatggaagtagcacgtctcactagtctc
gtgcagatggacagcaccgctgagcaatggaagcgggtaggcctttggggcagcggccaatagcagctttgctccttcgctttctgggctcagaggctgggaaggggt
gggtccgggggcgggctcaggggcgggctcaggggcggggcgggcgcccgaaggtcctccggaggcccggcattctgcacgcttcaaaagcgcacgtctgccgc
gctgttctcctcttcctcatctccgggcctttcgacctcacgtgcgcatgattgaacaagatggattgcacgcaggttctccggccgcttgggtggagaggctattcggctat
gactgggcacaacagacaatcggctgctctgatgccgccgtgttccggctgtcagcgcaggggcgcccggttctttttgtcaagaccgacctgtccggtgccctgaatga
actgcaagacgaggcagcgcggctatcgtggctggccacgacgggcgttccttgcgcagctgtgctcgacgttgtcactgaagcgggaagggactggctgctattggg
cgaagtgccggggcaggatctcctgtcatctcaccttgctcctgccgagaaagtatccatcatggctgatgcaatgcggcggctgcatacgcttgatccggctacctgcc

cattcgaccaccaagcgaaacatcgcatcgagcgagcacgtactcggatggaagccggtcttgtcgatcaggatgatctggacgaagagcatcaggggctcgcgcca
gccgaactgttcgccaggctcaaggcgagcatgcccgacggcgaggatctcgtcgtgacccatggcgatgcctgcttgccgaatatcatggtggaaaatggccgctttt
ctggattcatcgactgtggccggctgggtgtggcggaccgctatcaggacatagcgttggctacccgtgatattgctgaagagcttggcggcgaatgggctgaccgcttc
ctcgtgctttacggtatcgccgctcccgattcgcagcgcatcgccttctatcgccttcttgacgagttcttctgagcgggactctgggtacctttaagaccaatgacttacaag
gcagctgtagatcttagccactttttaaaagaaaaggggggactggaagggctaattcactcccaacgaagacaagatctgctttttgcttgtactgggtctctctggttaga
ccagatctgagcctgggagctctctggctaactagggaacccactgcttaagcctcaataaagcttgccttgagtgcttcaagtagtgtgtgcccgtctgttgtgtgactctg
gtaactagagatccctcagacccttttagtcagtgtggaaaatctctagcagtagtagttcatgtcatcttattattcagtatttataacttgcaaagaaatgaatatcagagagt
gagaggaacttgtttattgcagcttataatggttacaaataaagcaatagcatcacaaatttcacaaataaagcatttttttcactgcattctagttgtggtttgtccaaactcatca
atgtatcttatcatgtctggctctagctatcccgcccctaactccgcccatcccgcccctaactccgcccagttccgcccattctccgccccatggctgactaattttttttattta
tgcagaggccgaggccgcctcggcctctgagctattccagaagtagtgaggaggcttttttggaggcctagggacgtacccaattcgccctatagtgagtcgtattacgc
gcgctcactggccgtcgttttacaacgtcgtgactgggaaaaccctggcgttacccaacttaatcgccttgcagcacatccccctttcgccagctggcgtaatagcgaaga
ggcccgcaccgatcgcccttcccaacagttgcgcagcctgaatggcgaatgggacgcgccctgtagcggcgcattaagcgcggcgggtgtggtggttacgcgcagc
gtgaccgctacacttgccagcgccctagcgcccgctcctttcgctttcttcccttcctttctcgccacgttcgccggctttccccgtcaagctctaaatcgggggctccctttta
gggttccgatttagtgctttacggcacctcgaccccaaaaaacttgattagggtgatggttcacgtagtgggccatcgccctgatagacggtttttcgccctttgacgttgga
gtccacgttctttaatagtggactcttgttccaaactggaacaacactcaaccctatctcggtctattcttttgatttataagggattttgccgatttcggcctattggttaaaaaat
gagctgatttaacaaaaatttaacgcgaattttaacaaaatattaacgcttacaatttaggtggcacttttcggggaaatgtgcgcggaacccctatttgtttatttttctaaatac
attcaaatatgtatccgctcatgagacaataaccctgataaatgcttcaataatattgaaaaaggaagagtatgagtattcaacatttccgtgtcgcccttattcccttttttgcgg
cattttgccttcctgtttttgctcacccagaaacgctggtgaaagtaaaagatgctgaagatcagttgggtgcacgagtgggttacatcgaactggatctcaacagcggtaa
gatccttgagagttttcgccccgaagaacgttttccaatgatgagcacttttaaagttctgctatgtggcgcggtattatcccgtattgacgccgggcaagagcaactcggtc
gccgcatacactattctcagaatgacttggttgagtactcaccagtcacagaaaagcatcttacggatggcatgacagtaagagaattatgcagtgctgccataaccatgag
tgataacactgcggccaacttacttctgacaacgatcggaggaccgaaggagctaaccgcttttttgcacaacatgggggatcatgtaactcgccttgatcgttgggaacc
ggagctgaatgaagccataccaaacgacgagcgtgacaccacgatgcctgtagcaatggcaacaacgttgcgcaaactattaactggcgaactacttactctagcttccc
ggcaacaattaatagactggatggaggcggataaagttgcaggaccacttctgcgctcggcccttccggctggctggtttattgctgataaatctggagccggtgagcgt
gggtctcgcggtatcattgcagcactggggccagatggtaagccctcccgtatcgtagttatctacacgacggggagtcaggcaactatggatgaacgaaatagacaga
tcgctgagataggtgcctcactgattaagcattggtaactgtcagaccaagtttactcatatatactttagattgatttaaaacttcatttttaatttaaaaggatctaggtgaagat
cctttttgataatctcatgaccaaaatcccttaacgtgagttttcgttccactgagcgtcagaccccgtagaaaagatcaaaggatcttcttgagatcctttttttctgcgcgtaat
ctgctgcttgcaaacaaaaaaaccaccgctaccagcggtggtttgtttgccggatcaagagctaccaactctttttccgaaggtaactggcttcagcagagcgcagatacc
aaatactgttcttctagtgtagccgtagttaggccaccacttcaagaactctgtagcaccgcctacatacctcgctctgctaatcctgttaccagtggctgctgccagtggcga
taagtcgtgtcttaccgggttggactcaagacgatagttaccggataaggcgcagcggtcgggctgaacggggggttcgtgcacacagcccagcttggagcgaacgac
ctacaccgaactgagatacctacagcgtgagctatgagaaagcgccacgcttcccgaagagagaaaggcggacaggtatccggtaagcggcagggtcggaacagga
gagcgcacgagggagcttccagggggaaacgcctggtatctttatagtcctgtcgggtttcgccacctctgacttgagcgtcgatttttgtgatgctcgtcaggggggcgg
agcctatggaaaaacgccagcaacgcggcctttttacggttcctggccttttgctggccttttgctcacatgttctttcctgcgttatcccctgattctgtggataaccgtattac
cgcctttgagtgagctgataccgctcgccgcagccgaacgaccgagcgcagcgagtcagtgagcgaggaagcggaagagcgcccaatacgcaaaccgcctctccc
cgcgcgttggccgattcattaatgcagctggcacgacaggtttcccgactggaaagcgggcagtgagcgcaacgcaattaatgtgagttagctcactcattaggcacccc
aggctttacactttatgcttccggctcgtatgttgtgtggaattgtgagcggataacaatttcacacaggaaacagctatgaccatgattacgccaagcgcgcaattaaccct
cactaaagggaacaaaagctggagctgcaagctt

SP1-Lenti-MHI (pLV[Exp]-Neo-{MCRE_spacer_SP1_promorer}>mCherry):
MCRE enhancer
*SP1* promoter
5′AATGTAGTCTTATGCAATACTCTTGTAGTCTTGCAACATGGTAACGATGAGTTAGCAACATGCCTTAC
AAGGAGAGAAAAAGCACCGTGCATGCCGATTGGTGGAAGTAAGGTGGTACGATCGTGCCTTATTAGG
AAGGCAACAGACGGGTCTGACATGGATTGGACGAACCACTGAATTGCCGCATTGCAGAGATATTGTAT
TTAAGTGCCTAGCTCGATACATAAACGGGTCTCTCTGGTTAGACCAGATCTGAGCCTGGGAGCTCTCTG
GCTAACTAGGGAACCCACTGCTTAAGCCTCAATAAAGCTTGCCTTGAGTGCTTCAAGTAGTGTGTGCC
CGTCTGTTGTGTGACTCTGGTAACTAGAGATCCCTCAGACCCTTTTAGTCAGTGTGGAAAATCTCTAGC
AGTGGCGCCCGAACAGGGACTTGAAAGCGAAAGGGAAACCAGAGGAGCTCTCTCGACGCAGGACTCG
GCTTGCTGAAGCGCGCACGGCAAGAGGCGAGGGGCGGCGACTGGTGAGTACGCCAAAAATTTTGACT
AGCGGAGGCTAGAAGGAGAGAGATGGGTGCGAGAGCGTCAGTATTAAGCGGGGGAGAATTAGATCG
CGATGGGAAAAAATTCGGTTAAGGCCAGGGGGAAAGAAAAAATATAAATTAAAACATATAGTATGGG
CAAGCAGGGAGCTAGAACGATTCGCAGTTAATCCTGGCCTGTTAGAAACATCAGAAGGCTGTAGACA
AATACTGGGACAGCTACAACCATCCCTTCAGACAGGATCAGAAGAACTTAGATCATTATATAATACAG
TAGCAACCCTCTATTGTGTGCATCAAAGGATAGAGATAAAAGACACCAAGGAAGCTTTAGACAAGAT
AGAGGAAGAGCAAAACAAAAGTAAGACCACCGCACAGCAAGCGGCCGCTGATCTTCAGACCTGGAGG
AGGAGATATGAGGGACAATTGGAGAAGTGAATTATATAAATATAAAGTAGTAAAAATTGAACCATTA
GGAGTAGCACCCACCAAGGCAAAGAGAAGAGTGGTGCAGAGAGAAAAAAGAGCAGTGGGAATAGGA

GCTTTGTTCCTTGGGTTCTTGGGAGCAGCAGGAAGCACTATGGGCGCAGCGTCAATGACGCTGACGGT
ACAGGCCAGACAATTATTGTCTGGTATAGTGCAGCAGCAGAACAATTTGCTGAGGGCTATTGAGGCGC
AACAGCATCTGTTGCAACTCACAGTCTGGGGCATCAAGCAGCTCCAGGCAAGAATCCTGGCTGTGGAA
AGATACCTAAAGGATCAACAGCTCCTGGGGATTTGGGGTTGCTCTGGAAAACTCATTTGCACCACTGC
TGTGCCTTGGAATGCTAGTTGGAGTAATAAATCTCTGGAACAGATTTGGAATCACACGACCTGGATGG
AGTGGGACAGAGAAATTAACAATTACACAAGCTTAATACACTCCTTAATTGAAGAATCGCAAAACCA
GCAAGAAAAGAATGAACAAGAATTATTGGAATTAGATAAATGGGCAAGTTTGTGGAATTGGTTTAAC
ATAACAAATTGGCTGTGGTATATAAAATTATTCATAATGATAGTAGGAGGCTTGGTAGGTTTAAGAAT
AGTTTTTGCTGTACTTTCTATAGTGAATAGAGTTAGGCAGGGATATTCACCATTATCGTTTCAGACCCA
CCTCCCAACCCCGAGGGGACCCGACAGGCCCGAAGGAATAGAAGAAGAAGGTGGAGAGAGAGACAG
AGACAGATCCATTCGATTAGTGAACGGATCTCGACGGTATCGCTAGCTTTTAAAAGAAAAGGGGGGAT
TGGGGGGTACAGTGCAGGGGAAAGAATAGTAGACATAATAGCAACAGACATACAAACTAAAGAATTA
CAAAAACAAATTACAAAAATTCAAAATTTTACTAGTGATTATCGGATCAACTTTGTATAGAAAAGTTG
TTCCCCGCCGTTCCCCGCCGTTCCCCGCCGTTCCCCGCCGTTCCCCGCCGTCTGGTGGCGGAGGCTCGG
GCGGAGGTGGGTCGGGTGGCGGCGGATCAGGTTGACTGGTTTCCTTCCAAGCCAATCATCTCCAGCTC
CCGCCCATCTTCACTTCCTGCATCCTTCATTGGCTTTTAACACTGAGAGGGCGGTCTTTTTAGGCGGAC
ACCAGGCACGCAACTTAGTCTCACACGCCTTGGAGAGCAAGCGAGTCTTGCCATTGGATAATTCCACC
GTCTTTCTTCTGCAAGTCCCTCCTTTCCCCCTCCCTCATTGGGCGGGGCAGCAGAGAAGGGGCGGGGCC
TAGGTTGGGCTTGTGGCCAAGTTTGTACAAAAAAGCAGGCTGCCACCATGGTGAGCAAGGGCGAGGA
GGATAACATGGCCATCATCAAGGAGTTCATGCGCTTCAAGGTGCACATGGAGGGCTCCGTGAACGGCC
ACGAGTTCGAGATCGAGGGCGAGGGCGAGGGCCGCCCCTACGAGGGCACCCAGACCGCCAAGCTGAA
GGTGACCAAGGGTGGCCCCCTGCCCTTCGCCTGGGACATCCTGTCCCCTCAGTTCATGTACGGCTCCAA
GGCCTACGTGAAGCACCCCGCCGACATCCCCGACTACTTGAAGCTGTCCTTCCCCGAGGGCTTCAAGT
GGGAGCGCGTGATGAACTTCGAGGACGGCGGCGTGGTGACCGTGACCCAGGACTCCTCCCTGCAGGA
CGGCGAGTTCATCTACAAGGTGAAGCTGCGCGGCACCAACTTCCCCTCCGACGGCCCCGTAATGCAGA
AGAAGACCATGGGCTGGGAGGCCTCCTCCGAGCGGATGTACCCCGAGGACGGCGCCCTGAAGGGCGA
GATCAAGCAGAGGCTGAAGCTGAAGGACGGCGGCCACTACGACGCTGAGGTCAAGACCACCTACAAG
GCCAAGAAGCCCGTGCAGCTGCCCGGCGCCTACAACGTCAACATCAAGTTGGACATCACCTCCCACAA
CGAGGACTACACCATCGTGGAACAGTACGAACGCGCCGAGGGCCGCCACTCCACCGGCGGCATGGAC
GAGCTGTACAAGTAAACCCAGCTTTCTTGTACAAAGTGGTGATAATCGAATTCCGATAATCAACCTCT
GGATTACAAAATTTGTGAAAGATTGACTGGTATTCTTAACTATGTTGCTCCTTTTACGCTATGTGGATA
CGCTGCTTTAATGCCTTTGTATCATGCTATTGCTTCCCGTATGGCTTTCATTTTCTCCTCCTTGTATAAAT
CCTGGTTGCTGTCTCTTTATGAGGAGTTGTGGCCCGTTGTCAGGCAACGTGGCGTGGTGTGCACTGTGT
TTGCTGACGCAACCCCCACTGGTTGGGGCATTGCCACCACCTGTCAGCTCCTTTCCGGGACTTTCGCTT
TCCCCCTCCCTATTGCCACGGCGGAACTCATCGCCGCCTGCCTTGCCCGCTGCTGGACAGGGGCTCGGC
TGTTGGGCACTGACAATTCCGTGGTGTTGTCGGGGAAGCTGACGTCCTTTCCATGGCTGCTCGCCTGTG
TTGCCACCTGGATTCTGCGCGGGACGTCCTTCTGCTACGTCCCTTCGGCCCTCAATCCAGCGGACCTTC
CTTCCCGCGGCCTGCTGCCGGCTCTGCGGCCTCTTCCGCGTCTTCGCCTTCGCCCTCAGACGAGTCGGA
TCTCCCTTTGGGCCGCCTCCCCGCATCGGGAATTCCCGCGGTTCGAATTCTACCGGGTAGGGGAGGCG
CTTTTCCCAAGGCAGTCTGGAGCATGCGCTTTAGCAGCCCCGCTGGGCACTTGGCGCTACACAAGTGG
CCTCTGGCCTCGCACACATTCCACATCCACCGGTAGGCGCCAACCGGCTCCGTTCTTTGGTGGCCCCTT
CGCGCCACCTTCTACTCCTCCCCTAGTCAGGAAGTTCCCCCCCGCCCCGCAGCTCGCGTCGTGCAGGAC
GTGACAAATGGAAGTAGCACGTCTCACTAGTCTCGTGCAGATGGACAGCACCGCTGAGCAATGGAAG
CGGGTAGGCCTTTGGGGCAGCGGCCAATAGCAGCTTTGCTCCTTCGCTTTCTGGGCTCAGAGGCTGGG
AAGGGGTGGGTCCGGGGGCGGGCTCAGGGGCGGGCTCAGGGGCGGGGCGGGCGCCCGAAGGTCCTCC
GGAGGCCCGGCATTCTGCACGCTTCAAAAGCGCACGTCTGCCGCGCTGTTCTCCTCTTCCTCATCTCCG
GGCCTTTCGACCTCACGTGCGCATGATTGAACAAGATGGATTGCACGCAGGTTCTCCGGCCGCTTGGG
TGGAGAGGCTATTCGGCTATGACTGGGCACAACAGACAATCGGCTGCTCTGATGCCGCCGTGTTCCGG
CTGTCAGCGCAGGGGCGCCCGGTTCTTTTTGTCAAGACCGACCTGTCCGGTGCCCTGAATGAACTGCA
AGACGAGGCAGCGCGGCTATCGTGGCTGGCCACGACGGGCGTTCCTTGCGCAGCTGTGCTCGACGTTG
TCACTGAAGCGGGAAGGGACTGGCTGCTATTGGGCGAAGTGCCGGGGCAGGATCTCCTGTCATCTCAC
CTTGCTCCTGCCGAGAAAGTATCCATCATGGCTGATGCAATGCGGCGGCTGCATACGCTTGATCCGGC
TACCTGCCCATTCGACCACCAAGCGAAACATCGCATCGAGCGAGCACGTACTCGGATGGAAGCCGGTC
TTGTCGATCAGGATGATCTGGACGAAGAGCATCAGGGGCTCGCGCCAGCCGAACTGTTCGCCAGGCTC
AAGGCGAGCATGCCCGACGGCGAGGATCTCGTCGTGACCCATGGCGATGCCTGCTTGCCGAATATCAT
GGTGGAAAATGGCCGCTTTTCTGGATTCATCGACTGTGGCCGGCTGGGTGTGGCGGACCGCTATCAGG

ACATAGCGTTGGCTACCCGTGATATTGCTGAAGAGCTTGGCGGCGAATGGGCTGACCGCTTCCTCGTG
CTTTACGGTATCGCCGCTCCCGATTCGCAGCGCATCGCCTTCTATCGCCTTCTTGACGAGTTCTTCTGAG
CGGGACTCTGGGTACCTTTAAGACCAATGACTTACAAGGCAGCTGTAGATCTTAGCCACTTTTTAAAA
GAAAAGGGGGGACTGGAAGGGCTAATTCACTCCCAACGAAGACAAGATCTGCTTTTTGCTTGTACTGG
GTCTCTCTGGTTAGACCAGATCTGAGCCTGGGAGCTCTCTGGCTAACTAGGGAACCCACTGCTTAAGC
CTCAATAAAGCTTGCCTTGAGTGCTTCAAGTAGTGTGTGCCCGTCTGTTGTGTGACTCTGGTAACTAGA
GATCCCTCAGACCCTTTTAGTCAGTGTGGAAAATCTCTAGCAGTAGTAGTTCATGTCATCTTATTATTC
AGTATTTATAACTTGCAAAGAAATGAATATCAGAGAGTGAGAGGAACTTGTTTATTGCAGCTTATAAT
GGTTACAAATAAAGCAATAGCATCACAAATTTCACAAATAAAGCATTTTTTTCACTGCATTCTAGTTGT
GGTTTGTCCAAACTCATCAATGTATCTTATCATGTCTGGCTCTAGCTATCCCGCCCCTAACTCCGCCCAT
CCCGCCCCTAACTCCGCCCAGTTCCGCCCATTCTCCGCCCCATGGCTGACTAATTTTTTTTATTTATGCA
GAGGCCGAGGCCGCCTCGGCCTCTGAGCTATTCCAGAAGTAGTGAGGAGGCTTTTTTGGAGGCCTAGG
GACGTACCCAATTCGCCCTATAGTGAGTCGTATTACGCGCGCTCACTGGCCGTCGTTTTACAACGTCGT
GACTGGGAAAACCCTGGCGTTACCCAACTTAATCGCCTTGCAGCACATCCCCCTTTCGCCAGCTGGCGT
AATAGCGAAGAGGCCCGCACCGATCGCCCTTCCCAACAGTTGCGCAGCCTGAATGGCGAATGGGACG
CGCCCTGTAGCGGCGCATTAAGCGCGGCGGGTGTGGTGGTTACGCGCAGCGTGACCGCTACACTTGCC
AGCGCCCTAGCGCCCGCTCCTTTCGCTTTCTTCCCTTCCTTTCTCGCCACGTTCGCCGGCTTTCCCCGTC
AAGCTCTAAATCGGGGGCTCCCTTTAGGGTTCCGATTTAGTGCTTTACGGCACCTCGACCCCAAAAAA
CTTGATTAGGGTGATGGTTCACGTAGTGGGCCATCGCCCTGATAGACGGTTTTTCGCCCTTTGACGTTG
GAGTCCACGTTCTTTAATAGTGGACTCTTGTTCCAAACTGGAACAACACTCAACCCTATCTCGGTCTAT
TCTTTTGATTTATAAGGGATTTTGCCGATTTCGGCCTATTGGTTAAAAAATGAGCTGATTTAACAAAAA
TTTAACGCGAATTTTAACAAAATATTAACGCTTACAATTTAGGTGGCACTTTTCGGGGAAATGTGCGCG
GAACCCCTATTTGTTTATTTTTCTAAATACATTCAAATATGTATCCGCTCATGAGACAATAACCCTGAT
AAATGCTTCAATAATATTGAAAAAGGAAGAGTATGAGTATTCAACATTTCCGTGTCGCCCTTATTCCCT
TTTTTGCGGCATTTTGCCTTCCTGTTTTTGCTCACCCAGAAACGCTGGTGAAAGTAAAAGATGCTGAAG
ATCAGTTGGGTGCACGAGTGGGTTACATCGAACTGGATCTCAACAGCGGTAAGATCCTTGAGAGTTTT
CGCCCCGAAGAACGTTTTCCAATGATGAGCACTTTTAAAGTTCTGCTATGTGGCGCGGTATTATCCCGT
ATTGACGCCGGGCAAGAGCAACTCGGTCGCCGCATACACTATTCTCAGAATGACTTGGTTGAGTACTC
ACCAGTCACAGAAAAGCATCTTACGGATGGCATGACAGTAAGAGAATTATGCAGTGCTGCCATAACC
ATGAGTGATAACACTGCGGCCAACTTACTTCTGACAACGATCGGAGGACCGAAGGAGCTAACCGCTTT
TTTGCACAACATGGGGGATCATGTAACTCGCCTTGATCGTTGGGAACCGGAGCTGAATGAAGCCATAC
CAAACGACGAGCGTGACACCACGATGCCTGTAGCAATGGCAACAACGTTGCGCAAACTATTAACTGG
CGAACTACTTACTCTAGCTTCCCGGCAACAATTAATAGACTGGATGGAGGCGGATAAAGTTGCAGGAC
CACTTCTGCGCTCGGCCCTTCCGGCTGGCTGGTTTATTGCTGATAAATCTGGAGCCGGTGAGCGTGGGT
CTCGCGGTATCATTGCAGCACTGGGGCCAGATGGTAAGCCCTCCCGTATCGTAGTTATCTACACGACG
GGGAGTCAGGCAACTATGGATGAACGAAATAGACAGATCGCTGAGATAGGTGCCTCACTGATTAAGC
ATTGGTAACTGTCAGACCAAGTTTACTCATATATACTTTAGATTGATTTAAAACTTCATTTTTAATTTAA
AAGGATCTAGGTGAAGATCCTTTTTGATAATCTCATGACCAAAATCCCTTAACGTGAGTTTTCGTTCCA
CTGAGCGTCAGACCCCGTAGAAAAGATCAAAGGATCTTCTTGAGATCCTTTTTTTCTGCGCGTAATCTG
CTGCTTGCAAACAAAAAAACCACCGCTACCAGCGGTGGTTTGTTTGCCGGATCAAGAGCTACCAACTC
TTTTTCCGAAGGTAACTGGCTTCAGCAGAGCGCAGATACCAAATACTGTTCTTCTAGTGTAGCCGTAGT
TAGGCCACCACTTCAAGAACTCTGTAGCACCGCCTACATACCTCGCTCTGCTAATCCTGTTACCAGTGG
CTGCTGCCAGTGGCGATAAGTCGTGTCTTACCGGGTTGGACTCAAGACGATAGTTACCGGATAAGGCG
CAGCGGTCGGGCTGAACGGGGGGTTCGTGCACACAGCCCAGCTTGGAGCGAACGACCTACACCGAAC
TGAGATACCTACAGCGTGAGCTATGAGAAAGCGCCACGCTTCCCGAAGAGAGAAAGGCGGACAGGTA
TCCGGTAAGCGGCAGGGTCGGAACAGGAGAGCGCACGAGGGAGCTTCCAGGGGGAAACGCCTGGTAT
CTTTATAGTCCTGTCGGGTTTCGCCACCTCTGACTTGAGCGTCGATTTTTGTGATGCTCGTCAGGGGGG
CGGAGCCTATGGAAAAACGCCAGCAACGCGGCCTTTTTACGGTTCCTGGCCTTTTGCTGGCCTTTTGCT
CACATGTTCTTTCCTGCGTTATCCCCTGATTCTGTGGATAACCGTATTACCGCCTTTGAGTGAGCTGATA
CCGCTCGCCGCAGCCGAACGACCGAGCGCAGCGAGTCAGTGAGCGAGGAAGCGGAAGAGCGCCCAAT
ACGCAAACCGCCTCTCCCCGCGCGTTGGCCGATTCATTAATGCAGCTGGCACGACAGGTTTCCCGACT
GGAAAGCGGGCAGTGAGCGCAACGCAATTAATGTGAGTTAGCTCACTCATTAGGCACCCCAGGCTTTA
CACTTTATGCTTCCGGCTCGTATGTTGTGTGGAATTGTGAGCGGATAACAATTTCACACAGGAAACAG
CTATGACCATGATTACGCCAAGCGCGCAATTAACCCTCACTAAAGGGAACAAAAGCTGGAGCTGCAA
GCTT 3′
